# Scalable and deep profiling of mRNA targets for individual microRNAs with chimeric eCLIP

**DOI:** 10.1101/2022.02.13.480296

**Authors:** Sergei A Manakov, Alexander A Shishkin, Brian A Yee, Kylie A Shen, Diana C Cox, Samuel S Park, Heather M Foster, Karen B Chapman, Gene W Yeo, Eric L Van Nostrand

**Affiliations:** Eclipse BioInnovations, San Diego, CA; Department of Cellular and Molecular Medicine, University of California San Diego, La Jolla, CA; Institute for Genomic Medicine, University of California San Diego, La Jolla, CA; Verna & Marrs McLean Department of Biochemistry & Molecular Biology, Baylor College of Medicine, Houston, TX.; Therapeutic Innovation Center, Baylor College of Medicine, Houston, TX

## Abstract

Our expanding knowledge of the roles small regulatory RNAs play across numerous areas of biology, coupled with the promise of RNA-targeted therapies and small RNA-based medicines, create an urgent need for tools that can accurately identify and quantify small RNA:target interactions at scale. MicroRNAs (miRNA) are a major class of small RNAs in plants and animals. The experimental capture of miRNA:mRNA interactions by ligation into chimeric RNA fragments in chimeric CrossLinking and ImmunoPrecipitation (CLIP) provides a direct readout of miRNA targets with high-throughput sequencing. Despite the power of this approach, widespread adoption of chimeric CLIP has been slow due to both methodological technical complexity as well as limited recovery of chimeric molecules (particularly beyond the most abundant miRNAs). Here we describe chimeric eCLIP, in which we integrate a chimeric ligation step into AGO2 eCLIP to enable chimeric read recovery. We show that removal of the cumbersome polyacrylamide gel and nitrocellulose membrane transfer step common to CLIP techniques can be omitted for chimeric AGO2 eCLIP to create a simplified high throughput version of the assay that maintains high signal-to-noise. With the increased yield of recovered miRNA:mRNA interactions in no-gel chimeric eCLIP, we show that simple enrichment steps using either PCR or on-bead probe capture can be added to chimeric eCLIP in order to target and enrich libraries for chimeric reads specific to one or more miRNAs of interest in both cell lines and tissue samples, resulting in 30- to 175-fold increases in recovery of chimeric reads for miRNAs of interest. We further demonstrate that the same probe-capture approach can be used to recover miRNA interactions for a targeted gene of interest, revealing both distinct miRNA targeting as well as co-targeting by several miRNAs from the same seed family. RNA-seq analysis of gene expression following miRNA overexpression confirmed miRNA-mediated repression of chimeric eCLIP-identified targets and indicated that probe-enriched chimeric eCLIP can provide additional sensitivity to detect regulated targets among genes that either contain or lack computationally predicted miRNA target sites. Thus, we believe that chimeric eCLIP will be a useful tool for quantitative profiling of miRNA targets in varied sample types at scale, and for revealing a deeper picture of regulatory networks for specific miRNAs of biological interest.

**Highlights:**

- No-gel chimeric eCLIP improves recovery of miRNA:mRNA interactions by 70-fold
- Probe- and PCR-enrichment deeply profiles mRNA targets of miRNAs of interest
- Chimeric eCLIP targets experimentally identify non-computationally predicted interactions
- Increased depth recovers ∼6 million miRNA:target chimeras in HEK293T

## Introduction

MicroRNAs (miRNAs) are small non-coding RNAs that regulate target genes via complementarity to messenger RNAs (mRNA), resulting in post-transcriptional repression of hundreds of mRNAs. Regulation via miRNA-mediated repression of gene expression has been shown to be involved in nearly every physiological system and misregulation of miRNA biology has been implicated in a broad spectrum of diseases ranging from cancer to neurodegenerative diseases (Quinlan et al., 2017; Rupaimoole and Slack, 2017). Many miRNAs also display tissue-, cell type-, or condition-specific expression patterns and play key roles in the regulation of developmental programs (DeVeale et al., 2021; Manakov et al., 2009). Consequently, miRNAs have become attractive tools and targets for biomedical advancements. Currently several small molecules and antisense oligos that target miRNA biogenesis as well as miRNA mimics themselves are in clinical trials as candidate therapies for diseases such as non-small cell lung cancer, keloid, chronic hepatitis C, cutaneous T-cell lymphoma and Alport’s syndrome (Rupaimoole and Slack, 2017; Zhu et al., 2020). Thus, the repertoire of miRNA targets is a key determinant of the biological role of a given miRNA (Ebert and Sharp, 2012). Active research and development in the area of RNA-targeted therapies creates a need for tools that can accurately profile miRNA:mRNA target interactions in different cell cultures and tissues at scale.

Generally, miRNAs exert their repressive regulatory function by guiding the RNA-induced silencing complex (RISC) to complementary target sites in the 3′ untranslated region (UTR) of target mRNAs resulting in mRNA degradation, translation inhibition, or sequestration (Bartel, 2018). Building upon this principle of sequence complementarity, various algorithms have been developed to predict miRNA:mRNA interactions throughout the transcriptome (Agarwal et al., 2015; Krek et al., 2005). Computational approaches typically focus on a small set of key features, including sequence complementarity particularly in nucleotides 2-8 (commonly referred to as the ‘seed’ region of the miRNA), and sequence conservation across species. However, many verified targets do not meet these standard criteria (Gebert and MacRae, 2019), and the reliance on conservation limits detection of species-specific interactions.

Experimental identification of direct miRNA interactions on a large scale has proven more challenging. Recent efforts have shown success at identifying candidate miRNA target sites *in vitro* using purified RISC complex pre-loaded with a miRNA of interest, followed by high-throughput RNA binding assays (Becker et al., 2019; McGeary et al., 2019). Although these studies allow unprecedented depth in exploring the binding kinetics of individual miRNAs, it is important to pair these approaches with methods to experimentally validate the presence of miRNA:target interactions *in vivo.* Such profiling of miRNA targets in cell culture or tissues typically relies on immunoprecipitation (IP) of Argonaute (AGO) protein components of the RISC, followed by converting associated RNA into libraries that can be subjected to high-throughput sequencing in order to quantify association. Early methods including RNA Immunoprecipitation (RIP) and CrossLinking and ImmunoPrecipitation (CLIP) of AGO proteins provided the first broad view of the miRNA interaction landscape, revealing principles of miRNA regulation mechanisms as well as an overview of mRNAs regulated by miRNAs (Chi et al., 2009; Hafner et al., 2010). Although these approaches do not explicitly identify the miRNA which recruits the RISC complex to identified AGO2 binding sites, further development of computational methods addressed this limitation by utilizing analysis of sequence and altered binding upon miRNA over-expression or knockdown to predict these specific miRNAs (Erhard et al., 2013; Majoros et al., 2013). Although the combination of AGO IP and computational analysis has the advantage of enabling prediction of interactions for all miRNAs in one experiment, there remains many situations where experimental mapping of direct miRNA:target interactions would be preferred.

To enable this direct experimental mapping of microRNA:target interactions, a suite of chimeric CLIP methods including CrossLinking And Sequencing of Hybrids (CLASH), modified iPAR-CLIP, and CLEAR-CLIP (Broughton et al., 2016; Grosswendt et al., 2014; Helwak et al., 2013; Moore et al., 2015) were developed. These methods use computational analysis coupled with experimental modification of the standard CLIP method to generate and identify miRNA:mRNA ‘chimeric’ reads which reflect ligation of a miRNA with the target RNA that the miRNA is bound to. This snap shot of *in vivo* miRNA:mRNA interactions provided a unique ability to characterize the principles of miRNA:target binding through both seed and auxiliary region base-pairing, and provided insight into how these rules can impact functional regulation. However, despite the obvious power of direct experimental identification of miRNA targets and recent efforts to improve the efficiency of these approaches in capturing additional chimeric molecules (Bjerke and Yi, 2020; Gay et al., 2018), widespread adoption of these methods remains limited, as both the poor recovery of AGO-crosslinked RNA as well as the low fraction of chimeric reads make it difficult to deeply profile the interactome of a miRNA of interest at a reasonable cost. We reasoned that the recent development of improved CLIP methods that increase the recovery of protein-bound RNA by more than a thousand-fold over prior CLIP methods (Lee et al., 2021; Van Nostrand et al., 2016; Zarnegar et al., 2016) represented an opportunity to expand the utility of chimeric CLIP approaches.

Here we describe chimeric eCLIP coupled with AGO2 immunoprecipitation, which builds upon the improvements we described in eCLIP by modifying the standard eCLIP assay with an added ligation step to encourage generation of chimeric reads from both cell lines (HEK293T) and tissues (mouse liver). Further, we find that in the case of chimeric AGO2 CLIP, omitting the SDS-PAGE and nitrocellulose membrane transfer steps dramatically increases recovery of miRNA chimeric reads and simplifies the workflow, and rigorously validate that this approach maintains high signal-to-noise ratio. With this increased recovery, we show that chimeric eCLIP can be combined with PCR-based or antisense oligonucleotide probe capture to enrich libraries for chimeric reads specific to one or more miRNAs or genes of interest, enabling deep profiling of miRNA regulatory networks of interest with a robust and simplified chimeric CLIP protocol.

### Design

Here, we combine chimeric CLIP methods, where chimeric fragments directly link miRNA and target RNA transcript within the same sequencing read to unambiguously identify miRNA targets, with the methodological (library preparation) improvements in eCLIP and library capture/enrichment approaches to develop technologies that enable scalable and deep profiling of miRNA targets, particularly for individual miRNAs of interest.

### Integration of chimeric ligation with eCLIP

Although chimeric ligation of small RNAs with their targets can occur at low frequency during standard CLIP (Broughton *et al*., 2016; Grosswendt *et al*., 2014), chimeric CLIP-seq approaches (including CLASH, CLEAR-CLIP, and modified iPAR-CLIP) have modified this approach by incorporating both a phosphorylation step (using 3’ phosphatase minus T4 Polynucleotide Kinase) as well as an additional ligation step (without adapters) to encourage proximity-based ligation and increase the frequency of chimeric fragments to as much as 5.3%, although it often remains less than 2% (Bjerke and Yi, 2020; Gay *et al*., 2018; Grosswendt *et al*., 2014; Helwak *et al*., 2013; Moore *et al*., 2015). However, the fact that the number of chimeras recovered per miRNA correlates well with miRNA abundance (Moore *et al*., 2015) coupled with the generally low efficiency of recovery of protein-crosslinked RNA with prior CLIP methods makes it challenging to obtain sufficient signal for individual miRNAs to perform traditional peak calling, particularly past the few most abundant miRNAs.

Thus, we set out to create a chimeric eCLIP method that combines the improved library preparation steps we developed in the enhanced CLIP (eCLIP) procedure (Van Nostrand *et al*., 2016) with these optimizations for recovery of miRNA:target chimeras. First, we incorporated the 3’ phosphatase minus T4 PNK and no-adapter ligation steps into a standard AGO2 eCLIP experiment. Next, as recovery of chimeric fragments requires that the reverse transcription read through the protein:RNA crosslink site (rather than terminate at this adduct, as often occurs in CLIP (Konig et al., 2010)), we utilized an altered Mn^2+^ buffer for reverse transcription that is common in RNA structure probing experiments (Siegfried et al., 2014) and that we previously showed to increase crosslink site readthrough in eCLIP (Van Nostrand et al., 2017b). Finally, as successful mapping of chimeric miRNA:target reads requires at least 40 nt total length (including both a 20-22nt miRNA and a sufficient target sequence to uniquely map), miRNA-only and other undesired smaller fragments could be further depleted from the final sequencing library by increasing the lower bound of the size selection performed at both the nitrocellulose membrane isolation and final library purification step (either by agarose gel purification or bead cleanup).

### Increased recovery of AGO2-associated RNA with no-gel chimeric eCLIP

In addition to being experimentally intricate and difficult to scale or automate, the SDS-PAGE, nitrocellulose transfer, and RNA extraction steps are some of the major points of sample loss during CLIP protocols. Thus, developing approaches to transition away from these steps towards a ‘no-gel’ approach, while avoiding an increase in background (either in co-immunoprecipitated proteins, or co-purification of non-crosslinked RNA), has been a major point of emphasis in the CLIP field (Ilik et al., 2020). Use of denaturing washes (typically coupled with the use of HIS, biotin, HALO, or other peptide tags that have high-affinity interaction with bead or other support structures) is one such avenue, though these stringent washes often result in decreased yields (Helwak *et al*., 2013; Moore *et al*., 2015). An appealing alternative approach as described in the qCLASH method is simply to remove the SDS-PAGE and nitrocellulose transfer steps (Gay *et al*., 2018). However, this modification will lead to at least some inclusion of additional background signal from non-crosslinked RNA or protein-crosslinked RNA outside of the excised membrane region in with-gel experiments, and to what degree this alteration increases background, particularly among potential post-lysis interactions, was not fully explored. To explore this, we performed a variety of analyses to compare both signal and background observed in no-gel versus with-gel chimeric eCLIP in HEK293T cells, and observed that no-gel chimeric AGO2 eCLIP recovered both non-chimeric peaks as well as chimeric reads that had similar enrichment for 3’ UTR and miRNA seed sequence motifs compared to with-gel chimeric eCLIP. Further, we observed an increased recovery of unique cDNA fragments (measured as a decrease in required PCR amplification), making no-gel chimeric eCLIP a robust and scalable way to generate miRNA interactome maps.

### Enrichment of miRNA or gene of interest by PCR or probe-based capture

The first discovered microRNA (lin-4) was initially characterized for its key role in *C. elegans* development (Chalfie et al., 1981), and since then individual microRNAs have been shown to play critical roles in cancer (Peng and Croce, 2016), stem cell self-renewal and differentiation (Gangaraju and Lin, 2009), and obesity and insulin homeostasis (Ying et al., 2017) among numerous other diseases, as well as serve as potential therapeutics (Rupaimoole and Slack, 2017). Thus, deep profiling of regulatory maps for individual miRNAs is needed to study the molecular mechanisms of how miRNA regulation alters physiology and disease. However, as the recovery of chimeras per miRNA was still correlated to miRNA abundance, even with the increased number of unique (non-PCR duplicate) fragments in no-gel chimeric eCLIP the necessary sequencing to generate robust maps for a miRNA of interest would be cost-prohibitive for most miRNAs. Thus, a method to specifically enrich the sequenced library for individual miRNAs of interest would enable us to make full use of this improved yield by allowing deeper profiling for biologically relevant miRNAs in a particular cell line or tissue system under study. Further, enrichment for reads at least 40 nt in length (thus potentially containing both a miRNA of interest and a fragment of target RNA of mappable length (>18nt)) could provide further enrichment for chimeras specifically.

PCR-based enrichment has long been used for targeted genomic sequencing approaches (Tewhey et al., 2009), and has the advantage of experimental simplicity. The most straightforward such approach to enrich for desired miRNA-containing fragments is to simply utilize a two-step PCR process, with the first step utilizing a PCR primer within the miRNA of interest, followed by a second round of PCR to incorporate full sequencing adapters. We found that this could be easily adapted to targeted amplification of chimeric eCLIP libraries, and successfully enabled selective enrichment for miRNAs of interest while only requiring the purchase of a single additional PCR primer per miRNA. However, although PCR enrichment is an easy way to allow for in-depth profiling of miRNA targets, the nature of using PCR introduces limitations. First, many mammalian miRNAs have extremely high or low GC content, making design of primers with reasonable melting temperatures challenging. Next, because of the use of a primer targeted to the miRNA, the final reads reflect amplified products and have lost the original miRNA sequence present in the miRNA:mRNA chimeric molecule, leading to loss of information about differential targeting of miRNA family members (which often only contain a single nucleotide difference), and isomiRs or other variations which have been described to play key roles in the processing and function of microRNAs (Ameres and Zamore, 2013; Cloonan et al., 2011). Additionally, false-positive chimeras can be introduced due to the PCR primer annealing to similar sequences elsewhere in the transcriptome.

To address this limitation, we leveraged hybrid capture enrichment as an alternative that preserves the native miRNA and target sequence. Enrichment of desired regions by annealing biotin- or surface-attached antisense oligonucleotide probes followed by stringent washing was a key advance in the development of whole-exome and other targeted genome sequencing approaches (Hodges et al., 2007; Okou et al., 2007). We found that hybrid capture is adaptable to the eCLIP procedure at the post RNA adapter-ligation stage by annealing to commercially synthesized biotinylated DNA oligonucleotide probes followed by capture on standard streptavidin beads, light washes, and recovery of enriched RNA by DNase treatment. For miRNA-targeted enrichment, we initially designed probes to include multiple copies of the same miRNA to obtain higher yield, but found that concatamers of distinct miRNAs could also be used to enrich for multiple miRNAs simultaneously. Similarly, we designed probes antisense to transcript 3’ UTR regions and observed that this could also enrich for chimeric miRNA:mRNA reads that deeply profile miRNAs interacting with a gene of interest. These approaches allow an unparalleled ability to deeply map candidate miRNA interactions for either a miRNA or gene of interest.

## Results

### Chimeric eCLIP recovers miRNA:mRNA chimeras

To confirm that chimeric eCLIP successfully recovers chimeric miRNA:target reads in a manner similar to prior CLASH approaches, we performed chimeric eCLIP on HEK293T cells using an AGO2 antibody previously validated for CLIP (Sternburg et al., 2018) and the additional 3’ phosphatase minus T4 PNK and no-adapter ligation steps described above along with standard eCLIP immunoprecipitation, adapter ligation, SDS-PAGE electrophoresis, nitrocellulose membrane transfer and RNA isolation, reverse transcription, and PCR amplification (Fig. 1A) (see Methods). IP-western blotting indicated successful immunoprecipitation of AGO2 (Sup. Fig. 1A), and visualization using a biotin on the RNA adapter indicated pulldown of crosslinked RNA that resolved to the AGO2 size in high RNase conditions (Sup. Fig. 1B). To confirm that we successfully enriched for AGO2 interactions, we first performed standard (non-chimeric) CLIP analysis, including adapter trimming, repetitive element removal, genomic mapping, PCR duplicate removal, and peak calling (see Methods). In replicate experiments sequenced to 144 and 145 million reads respectively, an average of 57.6% of peaks significantly enriched in IP versus paired input were in 3’ UTRs (11,282 out of 20,038 total and 10,502 out of 17,819 total in each replicate respectively), with another 17.4% (4,024 and 2,633 in two replicates, respectively) in coding sequence (CDS) (Fig. 1B). Although only an average of 2.9% of peaks overlapped microRNAs due to their limited number, weighting peaks by information content in IP versus input revealed that an average of 38.8% of total peak information was at microRNAs, confirming substantial enrichment (Fig. 1C). Our results indicate successful enrichment of both miRNAs and putative targets in 3’ UTR and CDS regions with AGO2 eCLIP.

**Figure 1.**
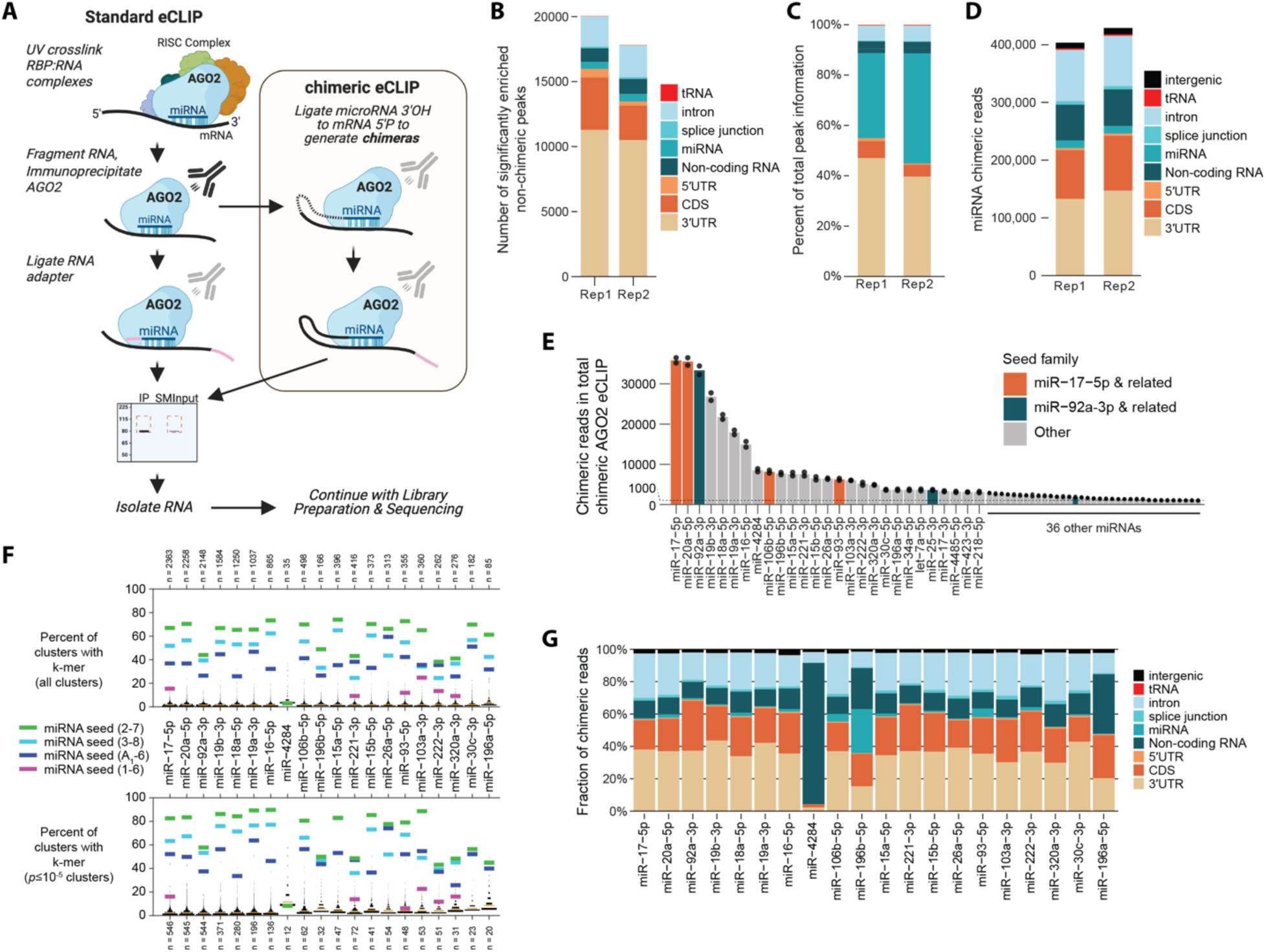
Identification of miRNA targets with chimeric eCLIP in HEK293T cells. (A) Schematic of alterations in chimeric eCLIP versus standard eCLIP of AGO2. (B-C) Chimeric eCLIP experiments analyzed with standard eCLIP peak calling using non-chimeric reads. Stacked bars indicate (B) the number of significantly enriched peaks (*p* ≤ 10^-3^ and fold-enrichment ≥ 8 in immunoprecipitation (IP) versus size-matched input) or (C) distribution of information content in significantly enriched peaks, with peaks annotated based on overlapping transcript regions (indicated with colors). (D) Stacked bars indicate the number of chimeric reads (miRNA plus a uniquely aligned target sequence) separated based on transcript regions overlapping chimeric read alignments. (E) Points indicate the number of chimeric reads observed per miRNA (with bar representing average across two replicates). Shown are all 62 miRNAs with >1000 average chimeric reads. Color indicates miRNAs from two abundant families in HEK293T cells. (F) For each of the top 20 miRNAs, lines indicate the percent of clusters containing either (colors) the indicated five 6-mers, or (black histogram) all other 6-mers. Orange line indicates mean of all other 6-mers. Data shown are from one representative replicate. Plots were generated using the MATLAB ‘distributionPlot’ package (v1.15) with option ’histOpt=0’ to plot histograms only. (G) Stacked bars indicate the number of chimeric reads per miRNA identified (average across 2 replicates) separated based on overlapping transcript regions.

Next, we developed an analysis workflow to identify chimeric reads based on a previously published ‘reverse mapping’ strategy (Moore *et al*., 2015). Although we started with all annotated miRNAs in miRbase, as part of this process we removed 15 miRNAs from analysis (corresponding to an average of 35% of potential chimeras) because both the miRNA and chimeric fragments mapped to rRNA, suggesting they likely represent annotation errors rather than true miRNAs (Supplementary Table 1). Confirming the ability of eCLIP to improve the recovery of unique (non-PCR duplicate) fragments, for the two replicates we identified a total of 403,532 and 428,936 unique chimeric reads (2.5% and 2.5% of uniquely mapped deduplicated reads, or 0.28% and 0.30% of 144 and 145×10^6^ initial sequenced reads respectively for the two replicates), including 132,801 and 146,966 3’ UTR chimeras (Fig. 1D), a dramatic increase over non-chimeric eCLIP or prior chimeric CLIP approaches. We observed high correlation between miRNA-only and both miRNA:chimera reads and independent small RNA-seq (Sup. Fig. 1C-E), consistent with previous chimeric CLIP methods (Moore *et al*., 2015). Separating chimeras by miRNA, these experiments yielded ∼7,000 to ∼36,000 chimeric reads per miRNA for the top 10 identified miRNAs, rapidly declining to less than 1,000 chimeric reads for the 63^rd^ most abundant miRNA (Fig. 1E). Notably, the six miRNAs processed from the mir-17-92 polycistronic cluster (He et al., 2005) were the six miRNAs with the most chimeric reads: the most abundant miRNA was miR-17-5p, which has previously been shown to regulate cell cycle progression in 293T cells (Cloonan et al., 2008), while miR-20a-5p, miR-92a-5p, miR-19b-3p, miR-18a-5p, and miR-19a-3p were second through sixth most abundant. In the top 25 miRNAs were also three miRNAs from the mir-106b-25 cluster, of which two (miR-106b-5p and miR-93-5p) share seed sequence with miR-17-5p and a third (miR-25-3p) shares seed sequence with miR-92a-3p. miR-25 was also proposed to be involved in the regulation of cancer (Sarkozy et al., 2018), suggesting that the most abundantly recovered miRNAs likely reflect important functional regulatory RNAs.

To confirm whether chimeric reads likely reflect true miRNA targets, we considered a variety of properties. First, we utilized the CLIPper algorithm (Lovci et al., 2013) to find clusters of chimeric reads for the 20 miRNAs with the greatest number of chimeric reads, identifying 15,222 and 15,805 chimeric clusters in two replicates, respectively. Motif analysis indicated that for the top 20 miRNAs an average of 58.7% of clusters contained 6-mers complementary to the cognate miRNA 6-mer seed region in positions [2:7] (versus an average of 1.7% for non-seed 6-mers), and for 19 of the top 20 the miRNA seed was the most commonly observed 6-mer (Fig. 1F). With a more conservative threshold for clusters (CLIPper *p*≤10^-5^) this was further increased to 68.4% of clusters containing the miRNA 6-mer seed (versus 3.5% for non-seed k-mers), albeit with a low number of clusters for some miRNAs (Fig. 1F). One exception, miR-4284, had chimeras that predominantly mapped to mitochondrial transcripts and lacked seed matches, suggesting that this may not reflect a bona fide microRNA in 293T cells. Next, location analysis of chimeric read alignments again indicated an enrichment for expected target regions except for miR-4284, with an average of 33% of chimeric reads mapped to 3’ UTRs and additional 21% to CDS (Fig. 1G). Thus, these results suggest that the properties of chimeric reads obtained with chimeric eCLIP modifications are consistent with previous chimeric CLIP-seq approaches.

### No-gel chimeric eCLIP increases depth of miRNA chimeras recovered

The standard eCLIP protocol that chimeric eCLIP is based on includes SDS-PAGE protein gel electrophoresis, Western blot-like nitrocellulose membrane transfer, and manual cutting of the membrane to isolate protein-crosslinked RNA. These steps are performed for two purposes: first, non-crosslinked RNA does not transfer to nitrocellulose and is thus removed (Grosswendt *et al*., 2014), and second, denaturation allows removal of RNA crosslinked to co-immunoprecipitated unwanted proteins of different sizes than the targeted protein. However, in addition to being complex for novice users and limiting scalability and automated handling, we and others have observed that this transfer and isolation step by itself drives a dramatic reduction in experimental yield, and multiple recent modified RBP immunoprecipitation protocols have been described which leave out this step (Gay *et al*., 2018; Ilik *et al*., 2020; Patton et al., 2020). As our experience with other RBPs suggested that co-immunoprecipitation artifacts were heavily protein- and antibody- dependent, we set out to rigorously test whether removing these steps altered the composition of chimeric eCLIP-reads. To do this, we tested a simplified protocol that removes the SDS-PAGE and membrane transfer steps and replaces it with a simple Proteinase K treatment to isolate the crosslinked RNA (“no-gel” variant of chimeric eCLIP, Fig. 2A). We observed that removal of the gel transfer steps required on average 6.3 fewer PCR cycles of amplification, suggesting a >70-fold increased experimental yield (Fig. 2B). Using previous estimates of conversion of eCT to non-PCR duplicate reads (Van Nostrand et al., 2020a), this suggests that no-gel AGO2 eCLIP recovers billions of unique RNA fragments.

**Figure 2.**
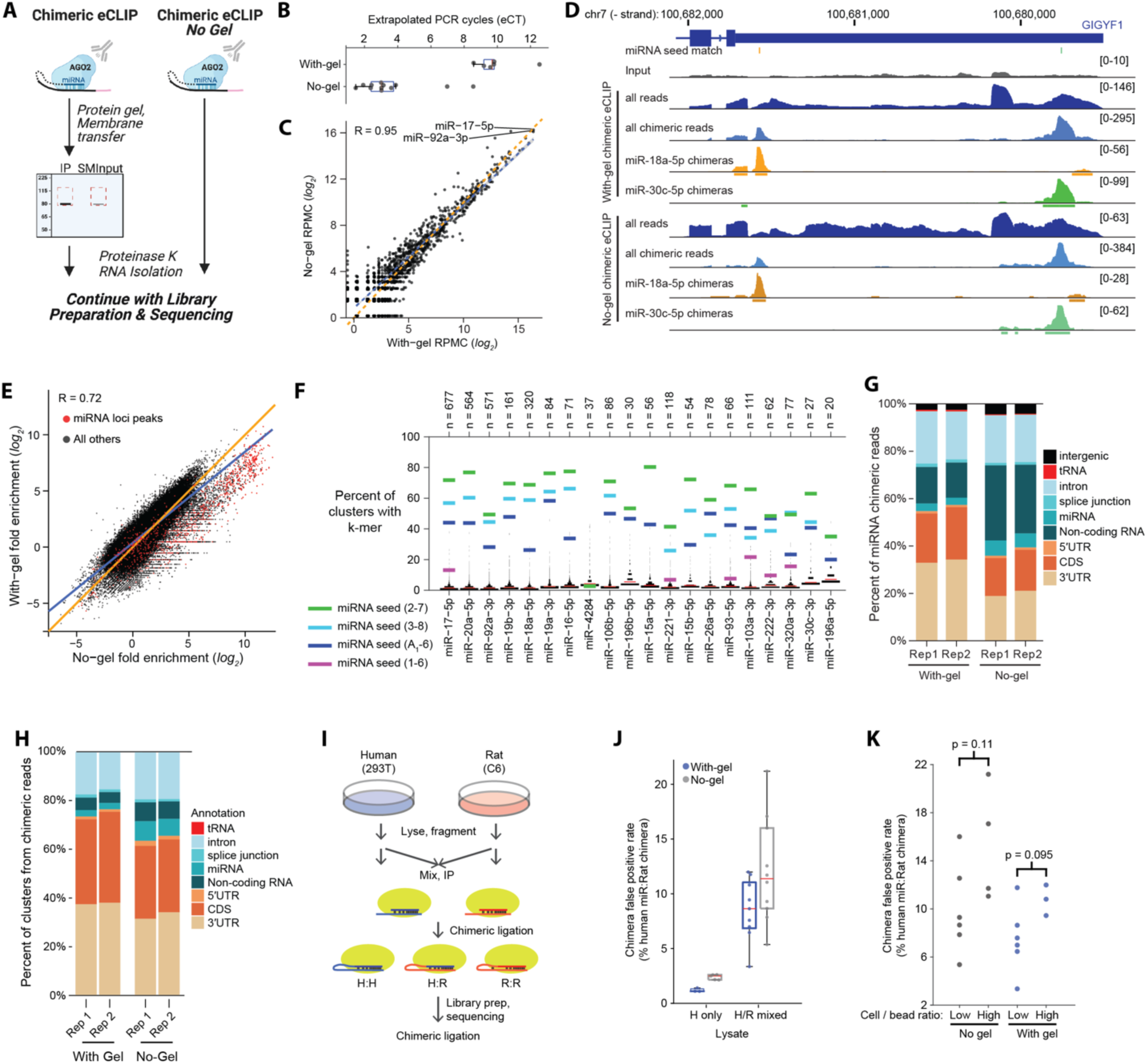
Improved recovery of chimeric miRNA fragments with no-gel chimeric eCLIP. (A) Schematic for no-gel versus standard chimeric eCLIP. (B) Points indicate yield (represented as extrapolated number of PCR cycles required to obtain 100 fmoles of amplified library, assuming 2-fold amplification per cycle). Experiments shown include standard and no-gel datasets described here as well as human-only samples from mixing experiments below. (C) Points indicate miRNA abundance among chimeric reads for standard with-gel (x-axis) versus no-gel (y-axis) experiments, displayed as log_2_(Reads Per Million Chimeras (RPMC)). Pearson correlation (R) is indicated. Blue line indicates least squares linear regression, and orange line indicates equality. (D) Genome browser figure showing with- and no-gel chimeric eCLIP data for the GIGYF1 3′UTR. Boxes underneath tracks indicate CLIPper-identified clusters (with *p* < 0.05), and miRNA seed 2-7 sequence matches in the region are indicated above. Input and ‘all reads’ are shown as reads per million, ‘all chimeric reads’ is shown as reads per million chimeras, and miR- specific tracks are shown as the number of reads. (E) For each of 250,339 clusters identified from non-chimeric eCLIP analysis of no-gel chimeric eCLIP replicate 1, points indicate fold- enrichment in (x-axis) no-gel chimeric eCLIP replicate 1 versus (y-axis) with-gel chimeric eCLIP replicate 1, with Pearson correlation indicated. Points indicated in red reflect peaks overlapping miRNA loci. (F) For each of the top 20 miRNAs, lines indicate the percent of clusters containing either (colors) the indicated five 6-mers, or (black histogram) all other 6-mers. Red line indicates mean of all other 6-mers. Data shown are from one representative replicate. (G-H) Stacked bars indicate the fraction of (G) chimeric reads or (H) clusters identified from chimeric reads from standard and no-gel chimeric eCLIP, separated by overlapping transcript annotation (indicated by colors). (I) Schematic of mixing experiments between human (HEK293xT) and rat (C6) cell lines. (J-K) Points indicate ‘false positive’ rate per mixing experiment, determined by identifying the subset of chimeras where the target fragment maps uniquely either to human or rat and calculating the ratio of miRNA_Human_:target_Rat_ target chimeras divided by miRNA_Human_:target_Human_ + miRNA_Human_:target_Rat_. Standard (with-gel) experiments are indicated in blue and no-gel experiments are indicated in grey. (J) All experiments are shown, with red line indicating median, and boxes indicating 25^th^ to 75^th^ percentiles. (K) Points indicate datasets split by whether the ratio of cell input to secondary beads was high (≥10 million cells per 100 uL of beads) or low (≤1.5 million cells per 100 uL of beads). Significance was determined by two- tailed Kolmogorov-Smirnov test.

To query whether the no-gel approach faithfully recapitulated with-gel targets, we first considered the frequency of miRNA-only (non-chimeric) and miRNA-containing (chimeric) reads. We observed a high correlation between with-gel and no-gel approaches for both miRNA-only (Pearson correlation 0.950, P.Value < 2.2×10^-16^) (Sup. Fig. 2A) and miRNA-chimeric read counts (Pearson correlation 0.953, P.Value < 2.2×10^-16^) (Fig. 2C). Additionally, the correlation between miRNA non-chimeric and chimeric read counts within an experiment was similar between with- gel (R = 0.87 and R = 0.85, *p* < 6.1×10^-183^ and < 9.2×10^-182^) (Sup. Fig. 1C-D) and no-gel variants (Pearson R = 0.78 & R = 0.81, p < 3.8×10^-145^ and < 3.9×10^-165^ for replicate 1 and 2 respectively) (Sup. Fig. 2B-C)). These results suggest that the no-gel approach does not alter the observed pattern of miRNA enrichment. However, we did note that information at non-chimeric peaks overlapping miRNAs (as well as the frequency of miRNA-only reads generally) was higher in the no-gel experiments (Sup. Fig. 2D), and we replicated this observation even when explicitly size- separating large (>30nt) and small (<30) library fractions (Sup. Fig. 2E), suggesting that miRNAs might have lower efficiency of UV crosslinking to AGO2 than target regions due to their smaller size and location buried within AGO2, leading to decreased recovery in with-gel approaches that include denaturing steps that would remove non-crosslinked miRNAs.

Next, we considered candidate interactions, defined from both non-chimeric as well as chimeric analyses. Manual inspection suggested similar read density distributions of non-chimeric as well as chimeric eCLIP reads between with-gel and no-gel libraries at many significantly enriched peaks (Fig. 2D). Expanding this analysis transcriptome-wide among non-chimeric reads, we observed a high correlation in IP versus input enrichment between with-gel and no-gel libraries considering either all 250,339 clusters identified by CLIPper in the no-gel libraries (Pearson correlation 0.72, P.Value < 2.2×10^-16^) (Fig. 2E) or all 353,280 clusters identified in the with-gel libraries (Pearson correlation 0.80, P.Value < 2.2×10^-16^) (Sup. Fig. 2F), suggesting that no-gel chimeric eCLIP generally recapitulates with-gel enrichments (with clusters at miRNA loci showing higher enrichment due to the increased frequency of miRNA-only reads noted above). The enrichment for 3’ UTR and CDS regions and increase in frequency of seed matching *k-*mers was also preserved between with-gel and no-gel chimeric reads. Analysis of clusters identified from no-gel chimeric reads for the most abundant 20 miRNAs showed an average of 60.0% containing the miRNA seed match relative to an average of 2.5% for non-seed 6-mers. The miRNA position 2-7 seed match was the most commonly found 6-mer for 16 of the 20 miRNAs (Fig. 2F), and the percent of clusters containing the seed match was consistent between with- and no-gel experiments (Sup. Fig. 2G). Similarly, we observed high frequencies of 3’ UTR coverage for non-chimeric peaks (33.6% and 36.5% for replicates 1 and 2 respectively) (Sup. Fig. 2H), chimeric reads (18.8% and 21.1%) (Fig. 2G), and clusters called from all chimeric reads (31.5% and 34.1%) (Fig. 2H, Sup. Fig. 2I) in no-gel libraries as with-gel libraries. Similar results were seen considering no-gel chimeric reads for the most abundant 20 miRNAs individually (Sup. Fig. 2J). These results indicate that no-gel chimeric eCLIP is maintaining robust recovery of miRNA targets.

One distinction between the no-gel and with-gel experiments was increased signal in non-mRNA regions, particularly non-coding transcripts (where the percent of chimeric reads doubled from 15.4% and 14.8% to 31.7% and 29.0%) (Fig. 2G). The effect on the number of chimeric clusters was far less dramatic, with only slight increases for non-coding transcripts (average 4.7% in with- gel to 7.4% in no-gel) and intronic (average 16.4% of with-gel to 19.3% of no-gel clusters) (Fig. 2H and Sup. Fig. 2I). Although previous AGO2 CLIP and chimeric CLIP studies have sometimes yielded a significant number of peaks in introns or linked to rRNA, tRNA, or other non-coding RNAs (Chu et al., 2021; Helwak *et al*., 2013), the particular emergence of non-coding signal only in no-gel experiments here suggests that it likely represents false positive signal. Previous CLASH studies utilizing spike-ins of *E. coli* RNA and *Drosophila* S2 cells (Moore *et al*., 2015) or yeast lysate (Helwak *et al*., 2013) into human cell lysate estimated a low (1-5%) rate of interactions formed *in vitro* after cell lysis. However, recent work performing *in vitro* RNA binding assays using pre-formed miRNA-loaded Argonaute complexes has shown that these complexes readily bind RNA with similar targeting principles as *in vivo*-identified miRNA targets (Becker *et al*., 2019; McGeary *et al*., 2019), suggesting that it would not be surprising that such post-lysis interactions could occur.

To query this we performed chimeric eCLIP upon mixing of human (H; HEK293T) and rat (R; C6) cell lysates (Fig. 2I). Although the sequence similarity between human and rat means that only a subset of chimeric reads can be uniquely resolved, the cross-reactivity of the AGO2 antibody for successful IP of AGO2 protein in rat (Sup. Fig. 2K-L) enabled us to perform chimeric eCLIP in both species simultaneously, validating that both samples were optimally fragmented for successful ligation and incorporation into chimeric reads that had similar enrichment for 3’ UTR regions (with a higher intergenic fraction likely driven by less accurate bioinformatic removal of rRNA and other repetitive elements as well as poorer annotation of non-coding transcripts) (Sup. Fig. 2M-N). Next, using 3 paired human- versus rat-only with-gel chimeric eCLIP samples we identified 108 human-enriched and 28 rat-enriched miRNAs with at least 10-fold differential expression (Sup. Fig. 2O). Independent no-gel experiments showed similar sample-specific expression for these miRNAs, whereas intermediate miRNA frequencies were observed in mixed lysate samples (Sup. Fig. 2O). Next, we isolated species-specific chimeric fragments by discarding chimeras that mapped equally well to human and rat, identifying thousands which were chimeric for a human-specific microRNA and were uniquely aligned to either the human or rat transcriptomes (including exons and introns for all protein-coding genes) (Sup. Fig. 2P). We observed that in with-gel experiments, an average of 8.6% of human miRNA chimeras aligned to rat in human-rat mixtures, whereas only 1.2% were observed in human-only experiments (reflecting the baseline mapping error rate) (Fig. 2J, Sup. Fig. 2P). This rate was slightly increased in no-gel human-rat mixture experiments, where 12.1% of human miRNA chimeric regions aligned to rat (Fig. 2J). Similar rates were observed if only 3’ UTR-mapping chimeras were considered (Sup. Fig. 2Q). To query whether this was driven by post-lysis interactions of miR:AGO2 complexes or simply cross-ligation of nearby RNAs due to crowding of the beads, we compared the false-positive rate in high (≥10 million cells per 100 uL of beads) and low (≤1.5 million cells per 100 uL of beads), and observed a trend towards decreased rates with increased dilution, with diluted ligation decreasing the false-positive rate from 10.7% to 7.5% in with-gel and 15.3% to 10.0% in no-gel (Fig. 2K). Although this did not reach significance, we utilized this lower ratio for the experiments described above and all probe capture experiments below as it suggested that some fraction of these artifacts might be due to on-bead crowding. As the half of potential post-lysis interactions that pair a human miRNA with human target sequence are undetectable by these lysate mixing approaches, these results suggest that a substantial rate of false positives are present even in standard with-gel chimeric approaches and are moderately increased in the no-gel approach. This may explain the significant number of rRNA and other non-coding RNA chimeras often seen for many miRNAs in our and prior chimeric CLIP studies (Helwak *et al*., 2013), and provides further evidence that (as with any immunoprecipitation-based protocol) it is important to validate chimeric CLIP-based miRNA interactions with orthogonal approaches (see further discussion in transfection section below).

### Targeted enrichment of miRNA chimeras by PCR

As noted above, the number of per-miRNA chimeric reads in total chimeric eCLIP correlates with non-chimeric miRNA abundance (Sup Fig. 1C-D, Sup Fig. 2B-C), leading to low coverage beyond the first few miRNAs (Fig. 1E). As standard sequencing depth of with-gel chimeric eCLIP often saturated sequencing of non-PCR duplicate reads, there would be no benefit to deeper sequencing or miRNA-targeted enrichment. However, the ∼70-fold improved yield in the no-gel variant (Fig. 2B) suggested that deeper profiling of individual miRNA interactomes might now be achievable. To apply chimeric-eCLIP to directly profile individual miRNAs that were not adequately captured using total chimeric using targeted PCR amplification, we altered the standard PCR amplification in chimeric eCLIP to a two-step approach which first utilizes one universal primer and one primer targeting the miRNA of interest, followed by a second PCR with multiplexing sequencing primers (Fig. 3A). By requiring that this PCR yields at least a 40 nt insert (reflecting miRNA plus 18 nt of additional sequence), this approach allows enrichment not only of a miRNA of interest but also provides selection of chimeric rather than miRNA- or mRNA-only fragments.

**Figure 3.**
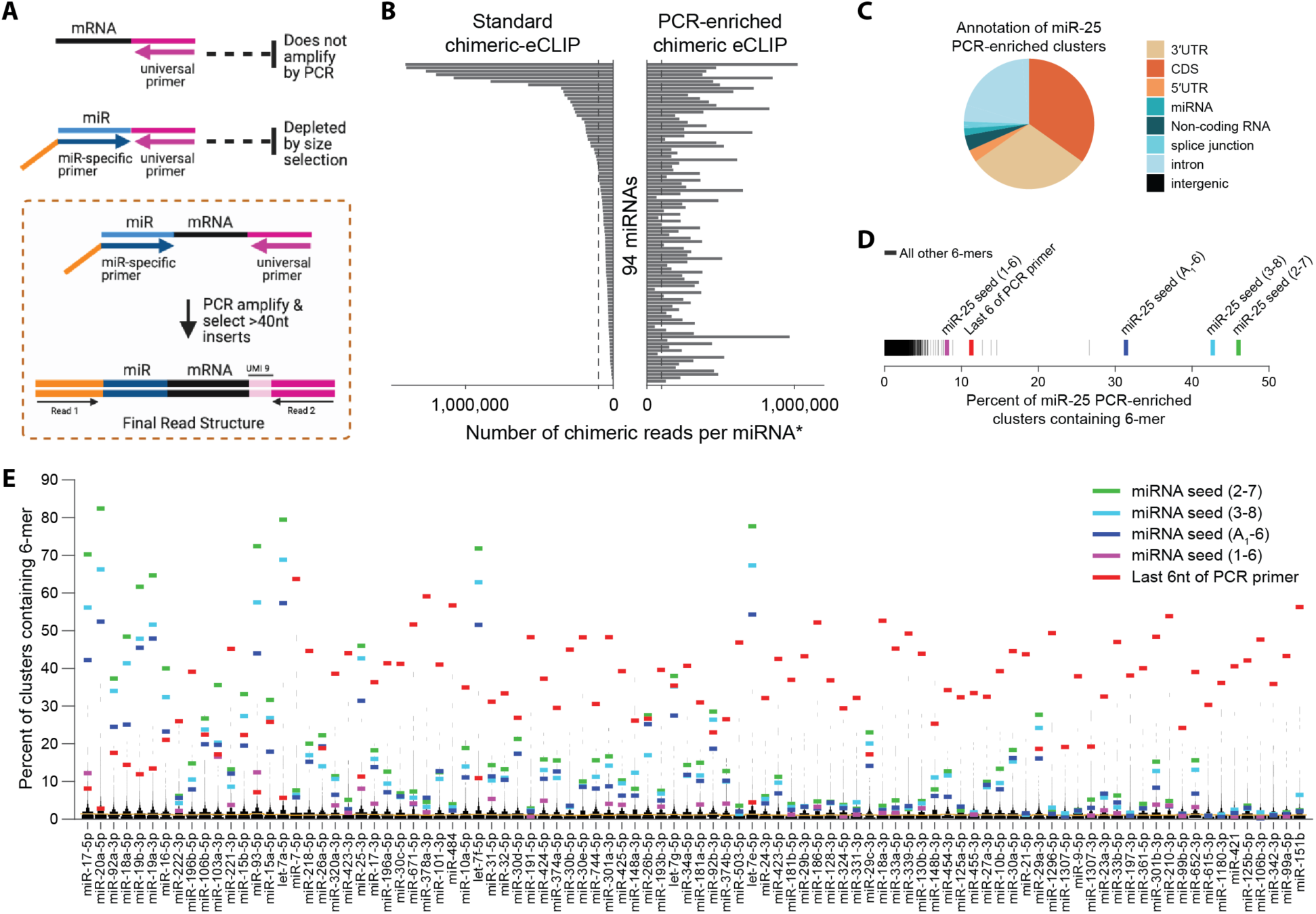
PCR enrichment of miRNAs of interest. (A) Schematic of enrichment of chimeras with a desired miRNA by targeted PCR amplification, using a miRNA-specific primer. (B) Bars indicate the number of chimeric reads per miRNA from (right) PCR enrichment for 94 separate miRNAs, or (left) extrapolated numbers for a paired total with-gel experiment (sequenced to 50.4×10^6^ reads) if it were sequenced to the same total depth (736×10^6^ reads) as the sum of all 94 separate PCR-enriched libraries. (C-D) Analysis of significant clusters (CLIPper *p* ≤ 10^-5^) from miR-25 PCR enrichment. (C) Pie chart indicates the fraction of clusters overlapping indicated transcript region annotations. (D) For all 6-mer sequences, lines indicate the percent of significant clusters containing each. Colors indicate 6-mers matching miR-25 seed sequences and the last 6 nt of the PCR primer. (E) For each of 94 human (hsa-) miRNAs profiled by PCR-enriched chimeric eCLIP, lines indicate the percent of significant clusters containing either (colors) the indicated five 6-mers, or (black histogram) all other 6-mers. Orange line indicates mean of all other 6- mers.

To test this approach, we performed PCR-enriched miRNA targeted no-gel chimeric-eCLIP on the most abundant 94 miRNAs in chimeric eCLIP in HEK293T cells, paired with a separate standard (non-selected) with-gel chimeric eCLIP. Sequencing to an average of 7.8 million reads per miRNA library, we obtained at least 100,000 chimeras for 89 of the 94 targeted miRNAs; extrapolating sequencing the non-enriched library to similar total depth (∼736 million reads) would yield 100,000 chimeras for only the most abundant 32 (Fig. 3B). Moreover, if one were interested in an individual miRNA, this approach increased the fraction of uniquely mapped chimeric reads from an average of 0.02% to 4.0% (a 175-fold increase) (Sup. Fig. 3A), which could likely be further improved by optimization of PCR amplification conditions. This leads to an increase in identified clusters (Sup. Fig. 3B), and a dramatic drop in required sequencing, as for example miR-25-3p went from 12,045 chimeric reads (out of 50.4 million) in total chimeric eCLIP to 253,151 (out of 9.0 million) in PCR-enriched chimeric eCLIP. For miR-25-3p, cluster identification on these chimeric reads yielded 2,877 significant clusters (CLIPper *p* ≤ 10^-5^). These clusters showed high enrichment for 3’ UTR (30.4%) and CDS (34.8%) regions (Fig. 3C), and motif analysis indicated that 46.0% of clusters contained the canonical miR-25 seed sequence (complementary to miR- 25 positions 2-7) (Fig. 3D). Expanding to all 94 miRs showed a similar annotation distribution, with average 39.9% 3’ UTR and 31.9% CDS clusters (Sup. Fig. 3C), and the 2-7 seed match sequence was the most abundant 6-mer for 21 (and in the top 10 for 47) out of the 94 profiled miRNAs (Fig. 3E).

However, we noted that many datasets also showed strong enrichment for the 6-mer at the end of the PCR primer (Fig. 3E), and manual inspection confirmed that these regions often contained stretches of homology to the miRNA that likely cause mis-priming of the PCR primer. Thus, although PCR-based enrichment can successfully yield deep profiling for some miRNAs, these primer-specific artifacts require careful consideration during analysis. To experimentally avoid these artifacts, we developed an alternative non-PCR based approach described below. However, we also hypothesized that as these artifacts were due to the miRNA-specific primer amplifying non-chimeric fragments, the false-positive clusters would likely be created in regions that were not enriched in the non-chimeric AGO2 eCLIP analysis. Consistent with this, motif analysis performed only on PCR-enriched clusters that overlapped reproducible AGO2 eCLIP peaks indicated that 70 of the 94 had an increase in the percent of clusters with the miRNA 2-7 seed match and 80 of the 94 had a decrease in the 6-mer at the 3’ end of the primer (Sup. Fig. 3D-E), suggesting that overlapping PCR-enriched datasets with standard AGO2 peaks can help remove false-positive signal and can help enable identification of which miRNA is causing a putative AGO2 interaction.

### Targeted enrichment of miRNA chimeras by probe-capture

As discussed in Design above, while PCR enrichment is an easy way to allow for in-depth profiling of miRNA targets, the replacement of the actual RNA fragment sequence with the primer sequence in the final read (and subsequent loss of information about related miRNAs, including miRNA family members which often only contain a single nucleotide difference, and miRNA 5’ ends) represents a clear limitation. Further, our results above suggested that PCR amplification artifacts could be a common source of false positives.

To address these concerns, we tested a probe-capture enrichment technique with modified oligonucleotides to increase the depth of chimeric read enrichment (Fig. 4A). First, we tested specificity of enrichment of chimeric reads for miRNAs of interest in HEK293T cells, choosing five miRNAs of interest (miR-221-3p, miR-34a-5p, miR-186-5p, miR-21-5p and miR-222-3p) that span a range of miRNA abundances from 15^th^ to 59^th^ most highly expressed miRNA in HEK293T (according to small RNA-seq profiling, Supplementary Table 2). We applied chimeric eCLIP to enrich libraries for chimeras of these miRNAs and compared it to libraries generated using with- gel chimeric eCLIP without enrichment. Probe capture-enriched chimeric eCLIP revealed unique targeting by the distinct miRNAs (Fig. 4B), and successfully enriched chimeras for each of the targeted miRNAs by more than 19-fold (Fig. 4C). The frequency of chimeras for the targeted miRNAs was increased from 0.01% to 0.48% of sequenced reads (Fig. 4D) resulting in a more than a 12-fold increase in the number of genes with reproducible 3’ UTR clusters (Fig. 4E). Out of all identified chimeras, the 5 targeted miRNAs went from 4.9% in standard chimeric eCLIP to 93.6% in the enriched pool (Fig. 4F), indicating a high specificity in recovering the desired miRNAs. Of note, we observed from 25,441 to 205,723 chimeric reads for each of the enriched miRNAs (Fig. 4C); achieving 25,000 chimeric reads for each would have required sequencing the non-enriched with-gel chimeric library 35-fold deeper (assuming there were sufficient non-PCR duplicates to make this possible). Clusters identified from chimeric reads in the probe-enriched experiment recapitulated specific enrichment for the miRNA seed sequence (Fig. 4G) and the read density within clusters was highly correlated with (while providing far deeper coverage compared to) standard chimeric eCLIP (R = 0.63, *p* = 9.8×10^-194^) (Sup. Fig. 4A), indicating that probe-capture enrichment maintained the same high signal-to-noise in identifying candidate miRNA targets.

**Figure 4.**
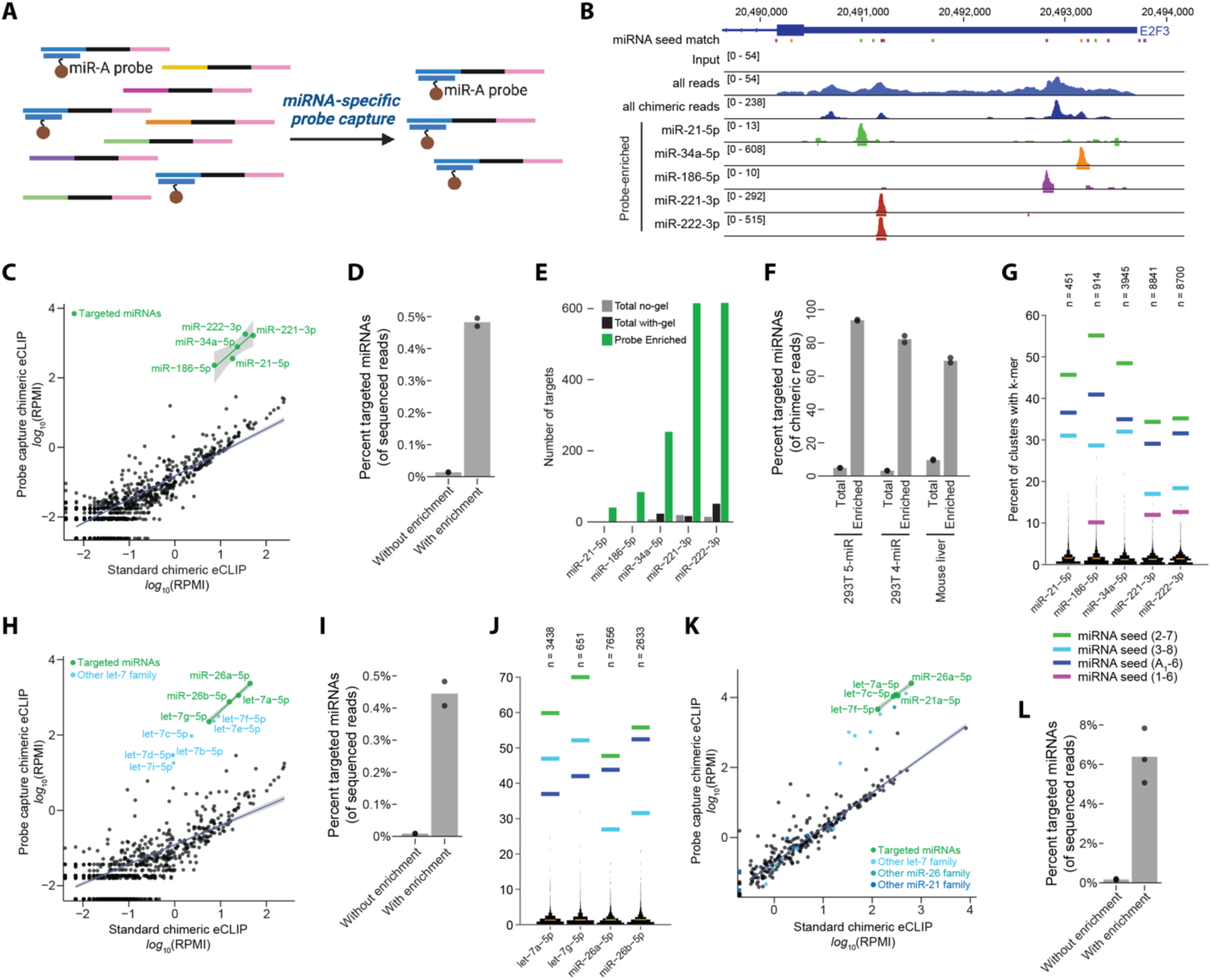
Probe-capture enrichment of miRNAs of interest. (A) Schematic of probe-capture enrichment for chimeras containing a targeted miRNA. (B) Genome browser image shows read density for the E2F3 3’UTR for non-chimeric analysis (‘all reads’), all chimeric reads from with-gel chimeric eCLIP, or miRNA-specific chimeric reads from probe-enriched chimeric eCLIP. Bars underneath tracks indicate CLIPper-identified clusters. (C-L) Shown are (C-G,F) enrichment of 5 miRNAs (hsa-) in HEK293T, (F,H-J) enrichment of 4 miRNAs (hsa-) in HEK293T, and (F,K-L) five indicated miRNAs (mmu-) in mouse liver tissue. (C,H,K) Points indicate chimeric reads in Reads Per Million Initial (RPMI) for all miRNAs in (x-axis) standard chimeric eCLIP, or (y-axis) probe capture-enriched chimeric eCLIP. Green points and line indicate targeted miRNAs, with a least squares linear model fit showing 0.95 confidence intervals. Light blue, dark blue and gray points indicate miRNAs that were not specifically targeted but are in the same family (share seed sequence) with targeted miRNAs. Black points and a robust linear regression model fit showing 0.95 confidence intervals represent all other miRNAs (except (K), where miRNAs with fewer than 20 chimeric reads in liver standard chimeric eCLIP experiment were omitted). (D,I,L) Points indicate percent of chimeric reads that represent the targeted miRNAs out of all sequenced reads, with bar indicating mean of replicate experiments. (E) Bars indicate the number of genes with reproducible clusters of chimeric reads in their 3’UTR regions for the indicated enriched miRNAs. (F) Points indicate percent of chimeric reads that represent the targeted miRNAs out of all chimeric reads, with bar indicating mean of replicate experiments. (G,J) For each of the targeted miRNAs, lines indicate the percent of clusters of probe capture enriched chimeric reads containing either (colors) the indicated five 6-mers, or (black histogram) all other 6-mers. Orange line indicates mean of all other 6-mers. Data shown are from one representative replicate.

Probe-capture chimeric-eCLIP can enrich for entire miRNA families while preserving the exact sequence of the specific miRNA bound to each target mRNA, enabling deep profiling of miRNA families with highly related sequences. Since many investigators are interested in studying families of miRNAs, we next tested whether probe capture could enable enrichment and subsequent separation of chimeric reads for members of the same miRNA family. First, we targeted six members of the miR-17 family that share the same seed sequence (miR-17-5p, miR- 93-5p, miR-20a-5p, miR-20b-5p, miR-106a-5p, miR-106b-5p) along with two miRNAs with related seed sites (miR-18a-5p, miR-18b-5p). This included two miRNAs (miR-17-5p and miR-20a-5p) that were highly over-represented among chimeric reads (Fig. 1E) as well as in small RNA-seq in HEK293T (Supplementary Table 2), and three miRNAs (miR-20b-5p, miR-106a-5p and miR-18b- 5p) that were ranked outside of the 80 most abundant miRNAs by chimeric frequency and outside the 200 most abundant miRNAs in small RNA-seq. Even though miRNAs that were selected in the miR-17 family experiment accounted for an average of 26.3% of total chimeric reads without enrichment, probe capture further increased representation of the selected miRNAs by 3.8-fold to 98.9% of chimeric reads and by 32.4-fold among all sequenced reads (from 0.07% to 2.4%) (Sup. Fig. 4B-D). Next, we designed probes against 2 members of the let-7 family (let-7a-5p and let-7g- 5p) along with two unrelated miRNAs (miR-26a-5p and miR-26b-5p). Chimeric reads for the targeted miRNAs in the let-7 family experiment were less common, accounting for only 0.01% before enrichment but increasing 49.4-fold to 0.4% following enrichment, with an increase from 3.2% of chimeric reads without enrichment to 82.2% after enrichment (Fig. 4F,H-I). We note that although other non-targeted let-7 members were also enriched, they were distinguishable by sequencing (Fig. 4H). Again, the increased read coverage led to a dramatic increase in the number of reproducible 3’ UTR clusters observed for the targeted miRNAs (Sup. Fig. 4E-F). In each case, the read density within clusters identified from chimeric reads was correlated with that observed from standard chimeric eCLIP (Sup. Fig. 4G-H), and clusters showed specific enrichment for sequences matching the seed region of the cognate miRNA (Fig. 4J, Sup. Fig. 4I), indicating that the more extensive set of probe-enriched targets retains the high signal-to-noise observed for non-enriched no-gel chimeric eCLIP (Fig. 2E).

Finally, we tested chimeric CLIP with probe capture enrichment of miRNA:mRNA chimeras in mouse liver tissue. Two sets of enriched libraries were prepared, one enriched for a selection of five miRNAs (miR-26a-5p, miR-21a-5p, let-7a-5p, let-7c-5p, let-7f-5p) and another enriched specifically for miR-122-5p. As above, probe-based enrichment further increased representation of chimeras for miRNAs of interest among chimeric reads, going from 9.6% to 69.2% of chimeras and from 0.2% to 6.4% of total reads for the 5-miRNA pool (Fig. 4F,K-L) and from 34.2% to 66.6% of chimeras and from 0.8% to 5.9% of total reads for miR-122-5p (Sup. Fig. 4J-L). As before, the number of clusters was dramatically increased in probe-capture experiments (Sup. Fig. 4M-N), and seed matching sites for miRNAs were over-represented in clusters called from chimeric reads (Sup. Fig. 4O-P). In summary, probe-enriched chimeric eCLIP libraries enable profiling of hundreds of thousands to several million chimeric reads per miRNAs of interest, allowing deep recovery of interactions for a miRNA of interest at substantially decreased sequencing depth.

### Profiling of miRNAs targeting a gene of interest

In addition to focused profiling of genes targeted by miRNAs of interest, it is increasingly important to identify miRNAs that target a specific gene of interest. To do this, we generated enrichment probes complementary to a gene of interest by ordering dsDNA gene fragments and performing T7 transcription with biotinylated nucleotides (see Methods), and performed capture of no-gel chimeric eCLIP cDNA as described above for miRNA-specific probes. As a proof of concept, we tested capture of two 3’ UTR regions: Unc-51 Like Autophagy Activating Kinase 1 (*ULK1*) and Amlyoid Beta Precursor Protein (*APP*). Although the frequency of chimeric reads obtained for a gene of interest was lower than for probe-capture enrichment of miRNAs, we observed 358-fold and 61-fold increased representation of chimeric reads for the gene of interest for ULK1 and APP enrichment respectively, going from less than 200 chimeric reads (out of 144 and 145 million sequenced reads) to over 1000 (out of less than 20 million reads) (Fig. 5A-B).

**Figure 5.**
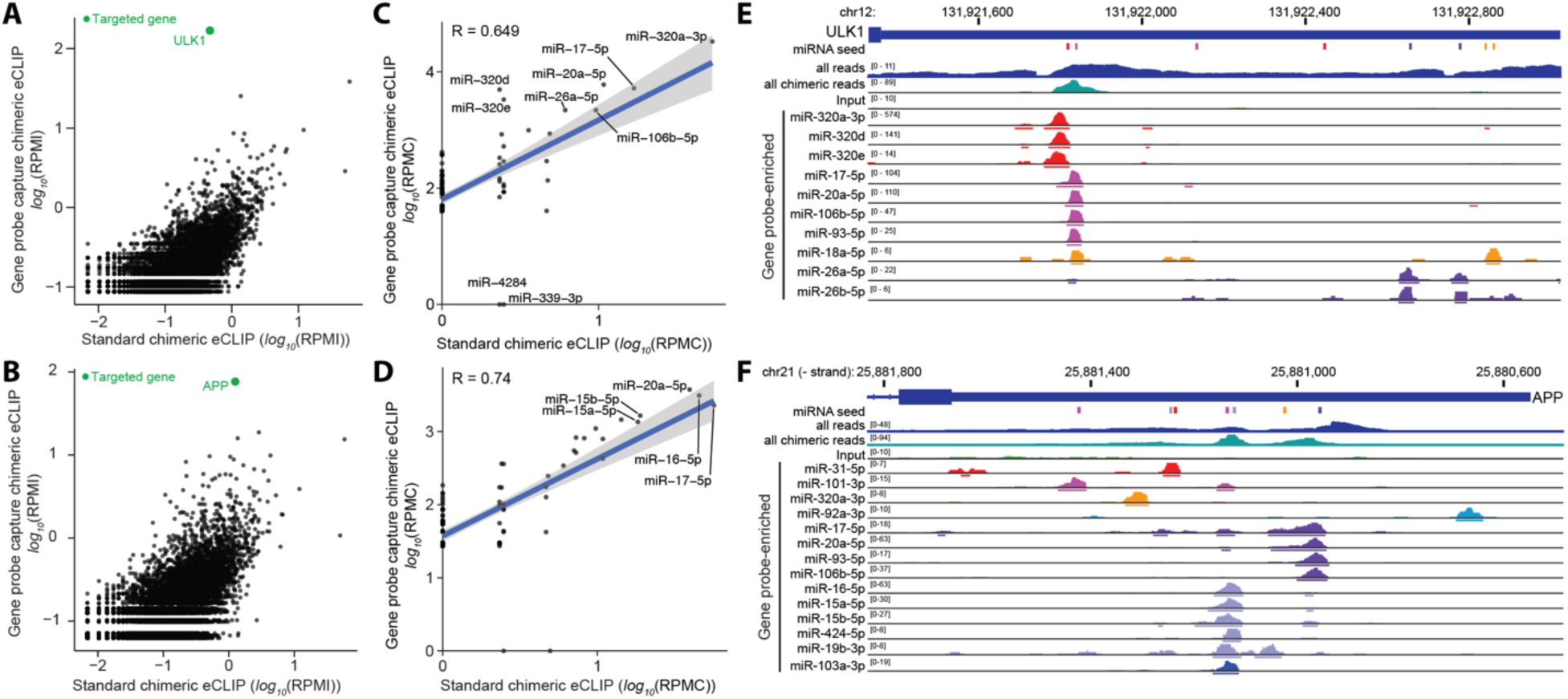
Probe-capture enrichment of genes of interest. (A-B) Points indicate chimeric read counts (per million initial reads) per gene in (x-axis) standard chimeric eCLIP, or (y-axis) probe capture- enriched chimeric eCLIP. Targeted genes (A) ULK1 or (B) APP are indicated in green. (C-D) Points indicate human (hsa-) per-miRNA chimeric read counts (per million chimeras) in standard chimeric eCLIP (x-axis) and probe capture-enriched chimeric eCLIP (y-axis) targeting 3’UTRs of (C) ULK1 and (D) APP. (E-F) Genome browser tracks show per-miRNA chimeric read density for 3’UTR regions of (E) ULK1 and (F) APP. ‘All reads’ and ‘all chimeric reads’ are from with-gel chimeric eCLIP, and miR-specific chimeric read tracks below are from gene-enriched chimeric eCLIP. All miRNA seed (2-7) matches are indicated, with colors indicating miRNA families sharing the same seed sequence.

Despite differences in chimeric read abundance with and without enrichment, counts of chimeric reads per miRNA were highly correlated between gene-enriched (no-gel) chimeric eCLIP and matched with-gel non-enriched chimeric eCLIP (Pearson correlation 0.65 and 0.75 for ULK1 and APP respectively) (Fig. 5C-D), indicating that the probe enrichment did not dramatically bias miRNA representation among gene specific chimeras. Upon visual inspection, we observed that individual target sites were well separated from each other, identifying numerous putative miRNA target sites in 3’ UTRs of ULK1 and APP (Fig. 5E-F). As expected, these sites commonly overlapped sequences complementary to the miRNA seed region. Notably, different miRNAs with the same or related seed sequences often (but not always) showed similar patterns of enrichment (Fig. 5E-F), reflecting the ability of probe capture enrichment to separate interactions of highly related miRNAs.

### miRNA:mRNA chimeras identify functional miRNA targets

As microRNAs often regulate gene expression by inducing RNA degradation, a common way to validate miRNA targets at scale is to show downregulation following miRNA overexpression (Lim et al., 2005). Indeed, targets identified using CLASH or similar chimeric ligation approaches showed enrichment for miRNA-dependent changes in RNA decay and specific functional categories, confirming that these methods yield high-quality sets of miRNA targets (Moore *et al*., 2015). To confirm that chimeric eCLIP also identifies functional miRNA targets, we chose two miRNAs with low endogenous expression in HEK293T cells (miR-1 and miR-124, ranked 66^th^ and 274^th^ most expressed miRNAs respectively (Supplementary Table 2) that have previously been used to study miRNA overexpression (Lim *et al*., 2005) and performed overexpression followed by small RNA-seq to confirm miRNA expression, mRNA-seq to assess the effect of miRNA overexpression on global gene expression, and chimeric eCLIP to identify targets (Fig. 6A).

**Figure 6.**
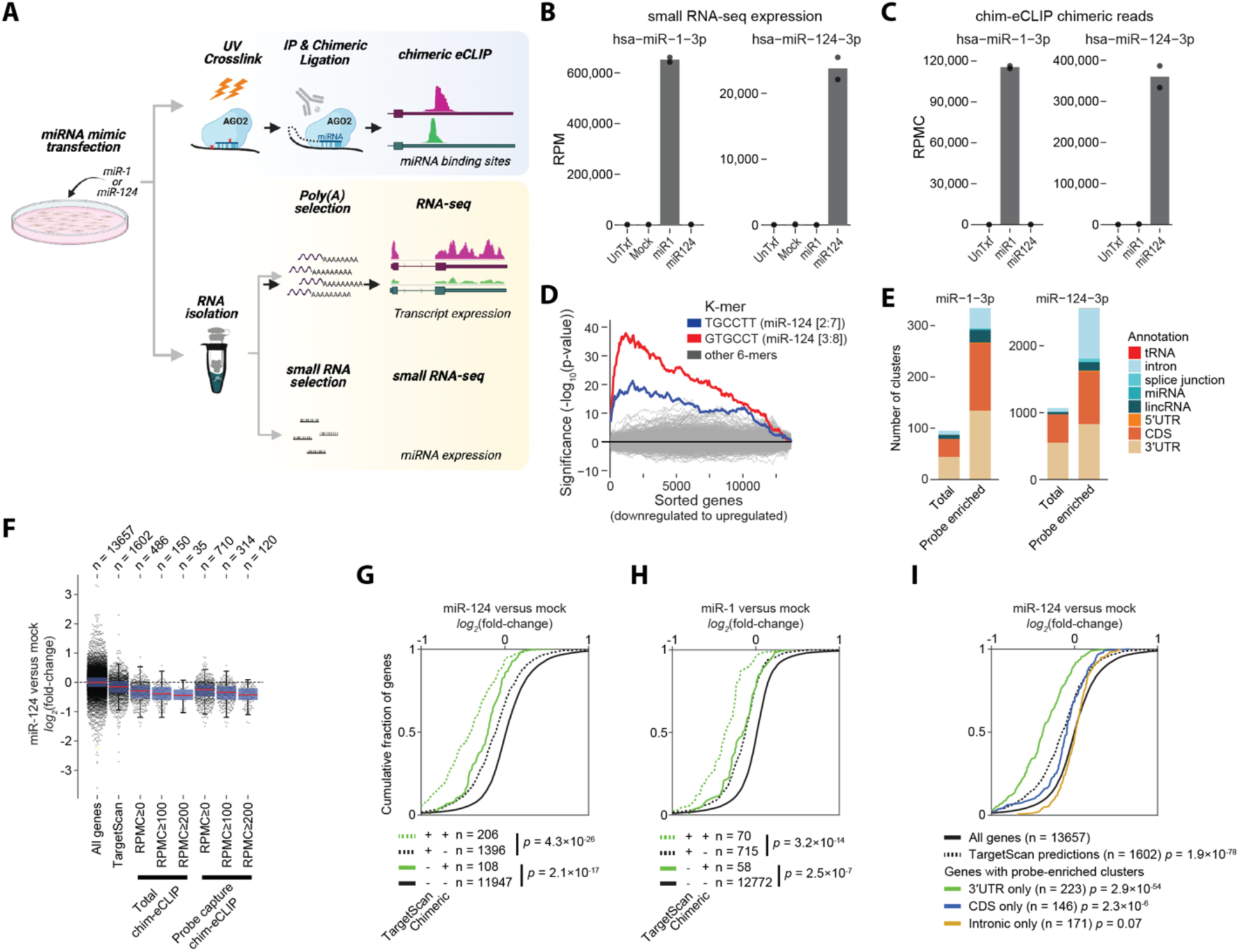
Validation of regulated miRNA targets with miRNA over-expression. (A) Schematic of miRNA over-expression experiments. (B-C) Points indicate abundance of two over-expressed miRNAs (miR-1-3p and miR-124-3p) in 2 replicate experiments, with bars indicating mean, for (B) reads per miRNA (RPM) from small RNA-seq and (C) chimeric reads per miRNA (RPMC) from chimeric eCLIP. (D) Lines show -log_10_ transformed hypergeometric p-values of enrichment of each of 2,332 6-mers complementing miRBase human miRNAs in [2:7] and [3:8] positions in growing bins of 3’UTR sequences of all expressed genes. Genes were sorted from down- to up- regulated upon miR-124 over-expression using test statistic associated with expression fold- change. (E) Stacked bars indicate number of reproducible (IDR > 540 across two replicates) clusters of chimeric reads from (left) miR-1 chimeric eCLIP in miR-1 over-expression or (right) miR-124 chimeric eCLIP in miR-124 over-expression, with overlapping transcript region annotation indicated by color. (F) For indicated gene classes, points indicate fold-enrichment in miR-124 over-expression. Red line indicates median, and blue boxes indicate 25^th^ to 75^th^ percentiles. Gene classes include those with TargetScan7.2-predicted miR-124 targets and genes containing chimeric eCLIP clusters meeting indicated chimeric reads per million (RPM) cutoff (average across two replicates) for either total (non-enriched) or miR-124 probe-enriched chimeric eCLIP. See Sup. Fig. 6I for miR-1 over-expression. Plots were generated using the MATLAB ‘plotSpread’ option in the “distributionPlot” package (v1.15). (G-I) Cumulative empirical distribution plots show fold-enrichment in miRNA over-expression for (G,I) miRNA-124 and (H) miRNA-1. Significance was determined by two-tailed Kolmogorov-Smirnov test. (G,H) Lines indicate sets of genes (green) with or (black) without TargetScan7.2-predicted miRNA interaction sites versus sets of genes (dotted lines) with or (solid lines) without clusters from chimeric eCLIP (using a cutoff of average RPMC≥100). (I) Lines indicate sets of genes with chimeric miR-124 clusters only in (green) 3’UTR, (blue) coding sequence, or (orange) intronic regions, versus (black) all genes or (dotted black) all genes with TargetScan7.2 predicted miR- 124 targets.

Both small RNA-seq and total chimeric eCLIP confirmed reproducible and specific enrichment of the overexpressed miRNA (Sup. Fig. 6A-C) as well as incorporation into mRNA-bound AGO2 (Sup. Fig. 6D-E), with both miR-1 and miR-124 showing dramatic up-regulation in both small RNA-seq (average 446- and 311-fold respectively versus mock transfection) and chimeric eCLIP (48,021- and 1,239-fold respectively) (Fig. 6B-C). Interestingly, although miR-1 was more abundant after over-expression in small RNA-seq (contributing more than 66% of reads (Fig. 6B)), miR-124 was more frequent in chimeras (more than 35% (Fig. 6C)), suggesting that the miR-124 mimic may be more easily incorporated into functional AGO2 complexes. Next, we used DESeq2 (Love et al., 2014) to quantify differential gene expression upon miRNA overexpression (Sup. Fig. 6F). Confirming that transfection of miR-1 and miR-124 mimics successfully induced repression of miR-1 and miR-124 targets respectively, 3’ UTRs of downregulated genes contained miR-1 and miR-124 seed matches more often than 3’ UTRs of upregulated genes (Sup. Fig. 6G), and genes containing complemented seed sites in 3’ UTRs showed the strongest shift towards gene repression for the two transfected miRNAs (Fig. 6D and Sup. Fig. 6H), consistent with prior observations (Lim *et al*., 2005).

To identify reproducible targets for miR-124 and miR-1, we first called clusters using miR-124 and miR-1 chimeric reads in each of the two replicates, and then used a modified IDR pipeline to identify reproducible clusters, which identified 549 miR-124 and 44 miR-1 3’ UTR clusters (Fig. 6E). Of genes also profiled in the miRNA over-expression RNA-seq, we observed a significant shift towards decreased expression for the 486 and 43 genes containing at least 1 chimeric 3’ UTR cluster for miR-124 and miR-1 respectively (*p* = 1.7×10^-83^ and *p* = 5.6×10^-18^ by two-tailed Kolmogorov-Smirnov test) (Fig. 6F, Sup. Fig. 6I). The magnitude of repression generally increased among chimeric eCLIP targets when more chimeric reads were identified (RPMC≥0, ≥100, or ≥200 chimeric reads per cluster), suggesting that the count of chimeric reads per target provides a metric that correlates with the regulatory impact of a particular miRNA:mRNA interaction.

To more deeply profile miR-1 and miR-124 targets, we additionally performed probe-enriched chimeric eCLIP, which expanded the set to 829 miR-124 and 134 miR-1 3’ UTR clusters (Fig. 6E). The 710 and 128 genes with probe-enriched 3’ UTR clusters for both miR-124 and miR-1 profiled in the over-expression RNA-seq showed enrichment for decreased expression upon miR over- expression (*p* = 1.6×10^-103^ and *p* = 6.2×10^-31^ respectively) (Fig. 6F, Sup. Fig. 6I). Moreover, genes (174 for miR-124 and 109 for miR-1) with reproducible chimeric clusters by probe capture but not identified in total chimeric still showed a significant shift towards repression upon miRNA over- expression (*p* = 1.6×10^-32^ and *p* = 5.4×10^-23^ respectively), which was only marginally less than those (140 for miR-124 and 19 for miR-1) detected by both methods (Sup. Fig. 6J-K), suggesting that the additional candidate targets recovered by deeper probe capture profiling remain enriched for functional miR targets.

Next, we compared these results against TargetScan (v 7.2) computationally predicted targets (Agarwal *et al*., 2015). TargetScan-predicted targets were more numerous than genes with chimeric eCLIP clusters and showed significant repression upon miRNA over-expression for both miR-124 and miR-1 (Fig. 6F, Sup. Fig. 6I), although the magnitude was more similar to only the low-confidence (RPMC≥0 read) chimeric eCLIP set. Notably, we observed that for both genes with or lacking TargetScan-predicted miRNA interactions, the subset of genes with chimeric eCLIP clusters showed greater repression upon miRNA over-expression (Fig. 6G-H, Sup. Fig. 6L-M), matching previous findings that chimeric CLIP can both refine computationally predicted targets as well as reveal new targets not captured by prediction alone (Bjerke and Yi, 2020).

MicroRNA sites have also been validated in coding sequence (CDS) regions, albeit with typically weaker repressive activity (Baek et al., 2008; Grimson et al., 2007). Consistent with these prior findings, we observed that the number of CDS clusters was roughly similar to that of 3’ UTR clusters (Fig. 6E), and that genes with chimeric clusters only in CDS for miR-124 (146) and miR- 1 (93) showed significant repression in miR over-expression (*p* = 2.3×10^-5^ a6d *p* = 7.3×10^-4^ respectively), albeit weaker than the repression observed for genes with 3’ UTR only clusters (Fig. 6I, Sup. Fig. 6N). This decreased effect for CDS targets versus 3’ UTR targets is consistent with prior observations, although notably for miR-124 the median *log_2_* fold-change for CDS-only chimeric targets (-0.093) was approaching the change observed for TargetScan-predicted 3’ UTR targets (-0.149) (Fig. 6I). As prediction of CDS targets remains challenging due to the high baseline conservation of coding regions, these results suggest that chimeric eCLIP represents a unique approach to enable exploration of functional CDS-region targeting by miRNAs.

### A resource of chimeric eCLIP data in 293T cells

Overall, the experiments described in our study profiled nearly six million non-PCR duplicate, uniquely mapped chimeric reads in HEK293T, including 2.6 million across 6 with-gel and 3.4 million across 12 no-gel experiments (Sup. Fig. 7A, Supplemental Table 3). To further aid the utilization of this resource, we merged these replicates together, yielding 963,813 and 856,953 3’ UTR chimeras for with-gel and no-gel respectively, and performed cluster identification for each miRNA with at least 1,000 chimeric reads, enabling identification of 31,494 putative high-confidence miRNA interaction sites in with-gel and 36,559 in no-gel respectively (CLIPper *p ≤* 10^-5^). As expected, coverage remained dependent on miRNA abundance, with miR-20a-5p and miR-17-5p yielding over 210,000 with-gel and 340,000 no-gel chimeras each; however, 46 miRNAs in the merged with-gel and 52 in the merged no-gel had at least 10,000 chimeras (Fig. 7A, Sup. Fig. 7A). These typically had hundreds to thousands of significant clusters located within the 3’ UTR or CDS of dozens to over a thousand unique genes (Fig. 7A, Sup. Fig. 7B), and these clusters were enriched for miRNA seed matches as expected, with the miRNA 2-7 seed being the most frequently observed 6-mer for 44 of the 46 miRNAs (Fig. 7B). Thus, this resource represents a unique opportunity to further explore the regulatory roles and networks of miRNAs in the well-characterized 293T cell line.

**Figure 7.**
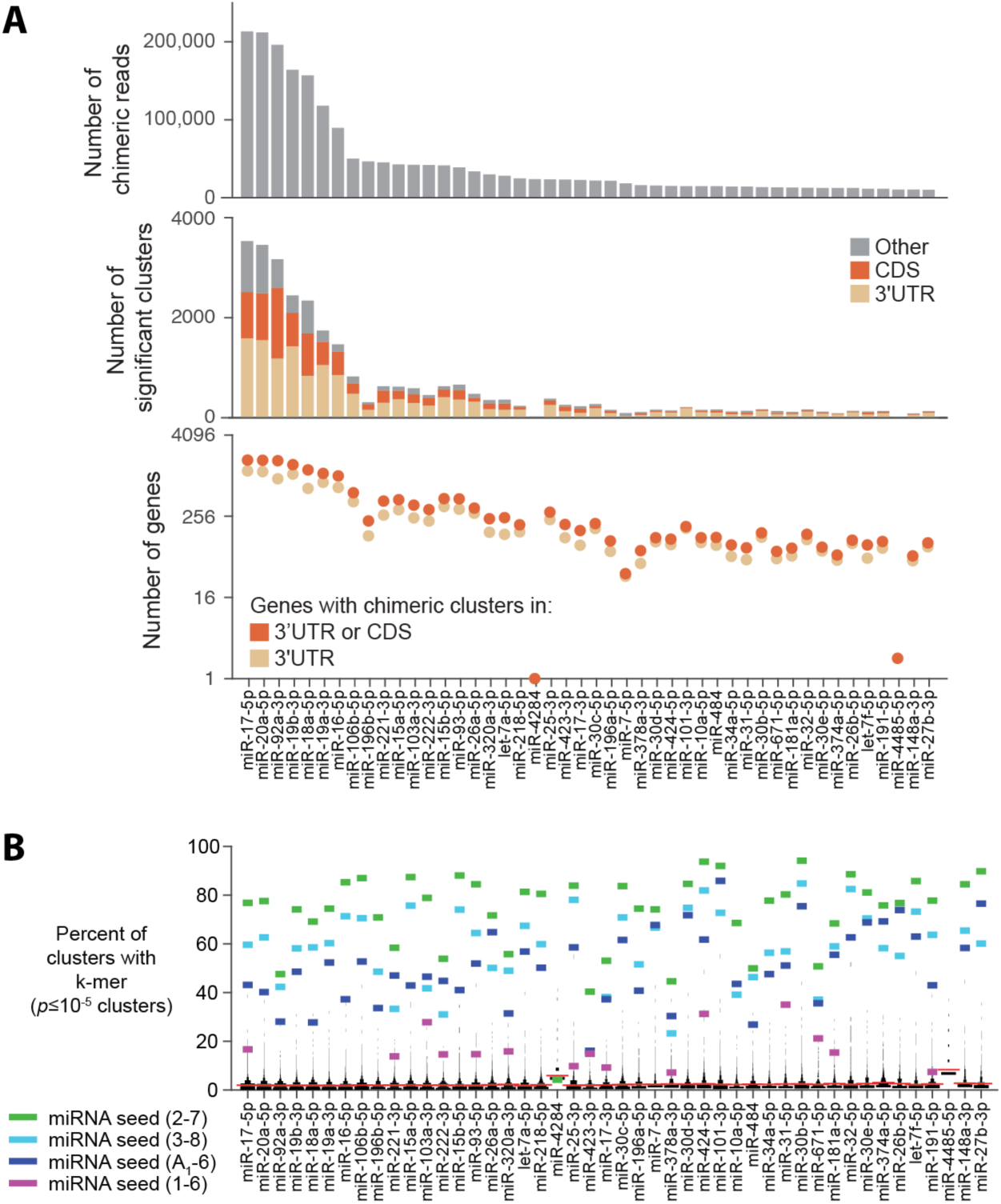
Deep profiling of miRNA interactomes in HEK293T. (A) Properties of the merged with-gel chimeric eCLIP data for the top 46 human (hsa-) miRNAs which each had more than 10,000 chimeric reads. (top) Bar indicates number of chimeric reads for each miRNA. (middle) Stacked bar indicates the number of significant (*p* ≤ 10^-5^) CLIPper clusters from chimeric reads for each miRNA, separated by overlapping annotation region. (bottom) On logarithmic scale, points indicate the number of unique genes containing significant clusters in either (tan) 3’UTR regions only or (orange) 3’UTR or CDS regions. See Supplementary Fig. 7 for merged no-gel data. (B) For each of the top 46 miRNAs, lines indicate the percent of clusters containing either (colors) the indicated five 6-mers, or (black histogram) all other 6-mers. Red line indicates mean of all other 6-mers.

## Discussion

The experimental identification of miRNA targets with chimeric CLIP-seq approaches provided unique insights into the rules for miRNA:target interactions (Broughton *et al*., 2016; Grosswendt *et al*., 2014; Helwak *et al*., 2013; Moore *et al*., 2015). However, the limited yield of non-PCR duplicate fragments, coupled with the low number of chimeric reads outside of the most abundant miRNAs, has hindered the widescale adoption of this powerful approach. With the ever-increasing catalog of miRNAs associated with human disease, coupled with the characterization of small RNAs produced by RNA viruses including SARS-CoV-2 that appear to utilize the host miRNA regulatory machinery (Pawlica et al., 2021), the ability to deeply profile targets for individual miRNAs of particular interest represents a novel opportunity to better understand the biological regulatory networks driven by individual microRNAs in a variety of context.

In this work, we describe both PCR- and probe-based strategies for targeted enrichment of a miRNA of interest. Although PCR-based enrichment is experimentally simpler and can yield robust profiling for many miRNAs, the common amplification of transcriptomic sites with sequence complementary to the primer coupled with the fact that the PCR primer replaces the sequence of the original miRNA (making it impossible to separate highly similar family members) led us to favor the probe-capture approach for general use. Because this approach simply enriches for (but does not alter) the original chimeric RNA fragments, it is thus possible to separate chimeras for miRNA family members differing only by a single nucleotide in order to reveal novel roles and interactions. Indeed, our analysis of miRNAs targeting two individual genes suggested a large degree of co-targeting among family members, which may help to explain the robustness of miRNA regulatory networks and their resilience to mutations of individual miRNAs in knockout experiments (Miska et al., 2007). Although we only performed analysis on annotated miRNAs in this work, this property may also enable querying of how alternative miRNA processing alters targeting, as recent studies indicated that alternative 3’ end tailing in particular can play critical roles (Kingston and Bartel, 2019). We note that although prior reports suggest this can also enable characterization of microRNA 5’ end isomiRs (Bjerke and Yi, 2020), in preliminary analyses we observed the addition of non-templated nucleotides after reverse transcription that may complicate this analysis (data not shown), and further work may be required to explore this question from the chimeric eCLIP data described here.

The utilization of an anti-AGO2 antibody to enrich for miRNA interactions has the benefit of enabling this approach to be utilized across cell lines and tissues with only limited experimental modification (optimization of lysis and RNase fragmentation conditions). However, the development of additional approaches, such as more stringent purification of AGO2:miRNA:mRNA complexes utilizing a transgenic AGO2:HaloTag fusion, could provide further opportunities for optimization and removal of background by enabling denaturing wash conditions (Li et al., 2020). In addition to microRNAs, there exist many other small regulatory RNAs that play critical roles in regulating RNA processing through interaction with complementary sequences, including piRNAs, snoRNAs, and others (Gumienny et al., 2017; Shen et al., 2018). Indeed, early CLASH publications showed the utility of the chimeric analysis approach in mapping snoRNA interactions in yeast (Kudla et al., 2011) and more recently was utilized to generate an extensive catalog of putative snoRNA interactions and reveal novel targets of orphan snoRNAs (Gumienny *et al*., 2017). Similarly, CLASH with PIWI protein has been used to reveal mechanisms of piRNA targeting (Shen *et al*., 2018), and CLASH using *E. Coli Hfq* to enrich for sRNA-mRNA interactions revealed principles of small RNAs in bacteria (Iosub et al., 2020). Thus, although we focus this description of chimeric eCLIP on miRNA target identification using immunoprecipitation of AGO2, we anticipate that the principles described here in both increasing experimental yield, as well as targeted enrichment for individual small RNAs of interest, should similarly enable deep profiling of other small RNA interactomes of interest.

## Limitations

Chimeric eCLIP, like all CLIP-based approaches, attempts to capture interactions as they occur in living cells. As such, whether observed interactions are truly occurring *in vivo* or are instead occurring after lysis is a major concern. In a traditional CLIP experiment, the denaturing SDS-PAGE and nitrocellulose transfer removes non-crosslinked RNA, providing a mechanism for removing potential post-lysis interactions. However, the additional chimeric ligation step in chimeric CLIP experiments (including with-gel chimeric eCLIP described here) weakens this limitation, as only one of the miRNA or target would be required to crosslink in a chimeric fragment. Prior chimeric CLIP experiments using spike-in of *E. coli*, Drosophila, or yeast lysate (Helwak *et al*., 2013; Moore *et al*., 2015) suggested a low (1-5%) rate of interactions formed *in vitro* after cell lysis; however, our work using fragmentation size-optimized human and rat lysate suggests this rate may be substantially higher (up to nearly 20%, once accounting for human-human post-lysis interactions that are not quantified by this approach). Thus, although it is always beneficial to validate CLIP-identified candidate interactions with orthogonal information (e.g., identifying overlap with *in vitro* motifs or regulation upon miRNA or RNA binding protein over- expression or knockdown), such orthogonal validation is particularly important if one wants robust confidence in individual microRNA targets identified by chimeric CLIP. Requiring crosslinking of both miRNA and mRNA target would reduce post-lysis interactions, but low efficiency of UV crosslinking currently limits such efforts (Moore *et al*., 2015). However, it is possible that this may be alleviated with improved crosslinking efficiency and the ability to utilize optimized denaturing wash conditions for Halo or other tag-based approaches.

Although we performed validation with miRNA overexpression, the high level of over-expression may not represent native miRNA targeting as recent studies have suggested that limited abundance of AGO2 itself may drive competition between miRNAs for proper loading and function (Khan et al., 2009; Tan et al., 2019). Further, while the PCR- and probe-based enrichment approaches described here enable deeper profiling of miRNAs of interest, the efficiency of these methods does remain constrained by both sequence properties of the miRNA or gene of interest, as well as the baseline expression level. Thus, in the implementation we describe here we do not directly address the stoichiometry of binding between miRNAs and their targets, and it remains an open question whether certain interactions identified by probe-enriched chimeric eCLIP reflect a high enough fraction of expressed copies of that mRNA to drive functionally relevant expression changes.

## EXPERIMENTAL MODEL AND SUBJECT DETAILS

### Cell culture

Human HEK293xT cells were acquired from Takara Bio and cultured in DMEM media (GIBCO) with 10% FBS 1% penicillin/streptomycin at 37°C with 5% CO_2_. C6 cells were purchased from Sigma and cultured in F-12 media (GIBCO) with 10% FBS, 2mM Glutamax, and 1X penicillin/streptomycin at 37°C with 5% CO_2_. For each experiment, 10 cm plates (∼15 million cells) were washed once with cold 1X phosphate buffered saline (PBS) and overlaid with minimal (3 mL per 10 cm plate) cold 1X PBS, and UV crosslinked (254 nm, 400 mJ/cm^2^) on ice. After crosslinking, cells were scraped and spun down, supernatant removed, and washed with cold 1X PBS. Cell pellets were flash frozen on dry ice and stored at -80 °C.

### Mouse liver tissue

8-week-old C57BL/6J mice were purchased from Jackson Labs. Mice were anesthetized with tribromoethanol and perfused with 0.9% saline. Tissues were collected, snap frozen, and stored at -80 °C for further analysis. Animal experiments were conducted under a protocol approved by the Institutional Animal Care and Use Committee (IACUC) of the University of California San Diego.

### miRNA mimic transfection

Human HEK293T cells grown to ∼75% confluency in antibiotic-free media were transfected with a final concentration of 100 nM of miR-1 or miR-124 miRNA mimics (IDT) or no mimic (mock) with Lipofectamine™ RNAiMAX (Invitrogen). Cells were incubated with mimics for 16 hours and then harvested (viability 70 - 80%). To harvest, cells were washed with cold 1X PBS, and either UV crosslinked as described above (for chimeric-eCLIP), or flash frozen without crosslinking (for RNA-seq and sRNA-seq).

### Chimeric-eCLIP

Chimeric eCLIP was based off the previously described seCLIP protocol (Van Nostrand et al., 2017a; Van Nostrand *et al*., 2016) with modifications to enhance chimera formation described below. As in eCLIP, lysis was performed in eCLIP lysis buffer, followed by sonication and digestion with RNase I (Ambion). Immunoprecipitation of AGO2-RNA complexes was achieved with a primary mouse monoclonal AGO2/EIF2C2 antibody (eIF2C2 4F9 (sc-53521) (Santa Cruz Biotechnology) overnight at 4°C using magnetic beads pre-coupled to the secondary antibody (M-280 Sheep Anti-Mouse IgG Dynabeads, Thermo Fisher 11202D). Initial experiments used standard eCLIP conditions (10 ug of antibody and 125 uL of Dynabeads for 20×10^6^ cells), but most experiments used decreased antibody and increased bead amounts based on the trend of decreased cross-species chimeras in those conditions (See Fig. 2K and Supplementary Table 4). Where indicated, 2% of each immunoprecipitated (IP) sample was saved as input control. For human/rat mixing experiments, cell pellets were lysed, sonicated, and RNase digested separately, and then mixed during addition of antibody and beads prior to overnight incubation. Western blot visualization used anti-AGO2 primary antibody (50683-RP02, SinoBiological) at 1:2000 dilution, with TrueBlot anti-rabbit secondary antibody (18-8816-31, Rockland) at 1:6000 dilution.

To phosphorylate the cleaved mRNA 5’-ends, beads were washed and treated with T4 polynucleotide kinase (PNK, 3’ -phosphatase minus, NEB) and 1 mM ATP. Chimeric ligation was then performed on-bead at room temperature for one hour with T4 RNA Ligase I (NEB) and 1 mM ATP in a 150 µl total volume. As in seCLIP, samples were then dephosphorylated with alkaline phosphatase (FastAP, Thermo Fisher) and T4 PNK (NEB), and an RNA adapter (N9RiT22 or CLASHn10RiL19bio) was ligated to the 3′ ends of the mRNA fragments (T4 RNA Ligase, NEB). With-gel chimeric-eCLIP IP and input samples were then denatured with 1X NuPage buffer (Life Technologies) and DTT, run on 4%–12% Bis-Tris protein gels and transferred to nitrocellulose membranes. The region corresponding to bands at the appropriate Ago2 protein size plus 75 kDa was excised and treated with Proteinase K (NEB) to isolate RNA, which was column purified (Zymo). No-gel chimeric eCLIP samples were treated directly with Proteinase K (NEB) to isolate RNA and column purified (Zymo). For both methods, RNA was then reverse transcribed with SuperScript IV Reverse Transcriptase (Invitrogen), 3 mM manganese chloride (to encourage read-through of crosslink sites), and 0.1 M DTT. Following reverse transcription (i16RT or InvAR19 primer), samples were treated as in seCLIP, including treatment with ExoSAP-IT (Affymetrix) to remove excess oligonucleotides, hydrolysis with sodium hydroxide (to degrade RNA) and addition of hydroden chloride (to balance pH). A 5’ Illumina DNA adapter (InvRand3Tr3) was then ligated to the 3′ end of cDNA fragments with T4 RNA Ligase (NEB), and after bead purification (Dynabeads MyOne Silane, Thermo Fisher), qPCR was performed on an aliquot of each sample to identify the proper number of PCR cycles (using D501_qPCR and D701_qPCR primers). The remainder of the sample was PCR amplified with barcoded Illumina compatible primers (Q5, NEB) based on qPCR quantification and size selected using AMPure XP beads (Beckman). Libraries were quantified using Agilent TapeStation and sequenced on the Illumina HiSeq or NovaSeq platform. As previously described, experimental yield was estimated by eCT, defined as the extrapolated number of PCR cycles necessary to obtain 100 femtomoles of library, assuming 2-fold amplification per PCR cycle (Van Nostrand *et al*., 2016). In this work, the eCT calculation also included extrapolation to normalize for cell number input and percent of cDNA used in order to enable comparison across experiments.

RNA visualization experiments were performed as previously described (Van Nostrand et al., 2020b) with the additional chimeric eCLIP steps added. Briefly, lysis of 3×10^6^ cells was performed with either low (3.3U) or high (100U) of RNase I (Ambion), with IP, chimeric (no-adapter) ligation, dephosphorylation, biotinylated adapter ligation, SDS-PAGE electrophoresis, and transfer to nitrocellulose membrane performed as described above. Detection of biotin-labeled RNA was performed using a Chemiluminescent Nucleic Acid Detection Module Kit (Thermo Scientific).

### PCR-based miRNA enrichment

MicroRNA-specific amplification primers were designed to minimize overlap with other miRNAs and limit overlapping with variable miRNA ends (Supplementary Table 5). To perform enrichment, first a qPCR was performed on diluted chimeric-eCLIP cDNA with miRNA-specific forward primer (Supplementary Table 5) and a reverse primer complementary to the RNA adapter sequence (chimeCLIP-7qL_R), and per-primer Ct values were obtained. Next, PCR amplification was performed in two steps. First, 8 cycles of PCR was performed with miRNA-specific forward primer and a reverse primer complementary to the RNA adapter sequence (chimeCLIP-7qL_R), using the standard eCLIP PCR conditions except for primer-specific annealing temperatures (see Supplementary Table 5). After bead cleanup with 2X volume Ampure XP beads (Beckman), a second PCR was performed using standard Illumina multiplexing primers and 8 cycles of amplification with standard eCLIP PCR conditions, using variable amount of first PCR product for each miRNA primer based on the qPCR Ct obtained (Supplementary Table 5). Each library was then purified again with 1.88X volume Ampure XP beads (Beckman), eluted, quantified by TapeStation (Agilent), and pooled for sequencing on the Illumina Nova Seq 6000 platform.

#### Probe-based miRNA capture

Samples were directly treated with Proteinase K (NEB) in place of the SDS-PAGE and membrane transfer steps described above. Biotinylated DNA probes designed (reverse complement) to the miRNA of interest (IDT) were then hybridized (500 picomoles per sample), washed on Silane beads (Dynabeads MyOne Silane, Thermo Fisher) and treated with DNase (Life Technologies). The remaining reverse transcription and library preparation steps were then performed as described above.

#### Probe-based gene capture

Samples were directly treated with Proteinase K in place of the SDS-PAGE and membrane transfer described above. Reverse transcription and cDNA adapter ligation steps were performed as above. Prior to PCR amplification, gblocks Gene Fragments (IDT) designed for the gene of interest were amplified to generate dsDNA templates. Biotinylated RNA probes were generated using T7 RNA Polymerase and biotinylated nucleotides (bio-UTP and bio- CTP). The biotinylated probes were coupled to streptavidin beads (Dynabeads MyOne Streptavidin C1, Thermo Fisher), chimeric library molecules were denatured, and chimeric molecules and probes were hybridized for one hour at 50°C. Beads were then washed, gene-specific probes were degraded with RNase, and enriched DNA fragments eluted from beads. The remaining PCR amplification and library preparation steps were then performed as described above.

### RNA-seq library preparation

15 million HEK293T cells were cells spun down, supernatant removed, and washed with cold PBS. Cell pellets were flash frozen on dry ice and stored at -80°C. Total RNA was isolated using the miRNeasy Mini Kit (Qiagen). Poly(A) RNA was isolated using Dynabeads Oligo (dT)_25_ (Thermo Fisher). RNA integrity and purity was measured using the Agilent 4200 TapeStation. To prepare RNA-sequencing libraries, 50 ng of poly(A) selected RNA was heat fragmented in 2X FastAP buffer and then treated with alkaline phosphatase (FastAP, Thermo Fisher) and T4 PNK (NEB). An RNA adapter was then ligated to the 3′-ends of the mRNA fragments (T4 RNA Ligase, NEB), after which RNA was column purified (Zymo) and reverse transcribed with SuperScript III Reverse Transcriptase (Invitrogen), then treated with ExoSAP-IT (Affymetrix) to remove excess oligonucleotides. A 5’ Illumina DNA adapter (InvRand3Tr3) was ligated to the 3′-end of cDNA fragments with T4 RNA Ligase (NEB) and after on-bead cleanup (Dynabeads MyOne Silane, Thermo Fisher), qPCR was performed on an aliquot of each sample to identify the proper number of PCR cycles. The remainder of the sample was PCR amplified with barcoded Illumina compatible primers (Q5, NEB) and size selected using AMPure XP beads (Beckman). Libraries were quantified using Agilent TapeStation and sequenced on the Illumina NovaSeq platform.

### sRNA-seq library preparation

HEK293fT cell pellets were prepared, and total RNA isolated as described in the RNA-seq library preparation Methods section. Small RNA-sequencing libraries were prepared using the QIAseq miRNA Library Kit (Qiagen). Briefly, a pre-adenylated DNA adapter was ligated to the 3’ ends of the RNA followed by ligation of an RNA adapter to the 5’ ends of the RNA. The adapter-ligated RNA was then reverse transcribed, and on-bead cleanup of cDNA was performed. Library amplification and barcoding was achieved using a universal forward primer and indexing 8-base reverse primers (HT Plate Indices 331565). Libraries were quantified using Agilent TapeStation and sequenced on the Illumina NovaSeq platform.

### Analysis of chimeric eCLIP sequencing data

The pipeline for analysis of chimeric eCLIP datasets is available at https://github.com/YeoLab/chim-eCLIP. Briefly, final library fragments in chimeric eCLIP libraries contain a sequence of 10 random nucleotides at the 5′ end and a sequence of 9 (N9RiT22), 10 (CLASHn10RiL19bio), or 0 (CLASHRiL19bio) random nucleotides at the 3′ end of the insert sequence. These random sequences serve as unique molecular identifiers (UMIs) and were utilized for removal of PCR duplicates (Kivioja et al., 2011). Only the 5′ end UMI was used for processing of total chimeric and probe capture enriched libraries; the 3′ end UMI was used only for PCR duplicate removal in PCR Targeted Chimeric analysis where the 5’ UMI is lost. In the first steps of the analysis, the 10 nucleotide UMIs were pruned from the 5′ end of R1 read sequences using umi_tools (v1.0.0) (Smith et al., 2017) and saved in the read name. Next, aggressive adapter trimming was performed to remove not only adapter sequences but also adapter fragment concatemers with cutadapt (v2.5) using options -O 1 --times 3, -e 0.1, --quality-cutoff 6, -m 18, and 10nt fragments that step through the RNA adapter (-a AGATCGGAAG -a GATCGGAAGA - a ATCGGAAGAG -a TCGGAAGAGC -a CGGAAGAGCA -a GGAAGAGCAC -a GAAGAGCACA -a AAGAGCACAC -a AGAGCACACG -a GAGCACACGT -a AGCACACGTC -a GCACACGTCT -a CACACGTCTG -a ACACGTCTGA -a CACGTCTGAA -a ACGTCTGAAC -a CGTCTGAACT - a GTCTGAACTC -a TCTGAACTCC -a CTGAACTCCA -a TGAACTCCAG -a GAACTCCAGT –a AACTCCAGTC -a ACTCCAGTCA). This aggressive trimming due to high adapter concatemer presence was later observed to be linked to RNA adapters containing random nucleotides on the 5’ end, which can be alleviated with the CLASHRiL19bio adapter (Supplementary Table 5). Remaining reads shorter than 18 nucleotides in length were discarded. The final 9 or 10 nucleotides at the 3′ end of each read were then removed using cutadapt (v2.5) to ensure possible remaining random sequence at the 3′ end of the insert sequence was removed. Further analysis was performed separately for analysis of non-chimeric reads and analysis of chimeric reads.

### Non-chimeric (standard eCLIP) analysis

Standard eCLIP analysis was performed as previously described (Van Nostrand *et al*., 2016). Briefly, reads were mapped to a database of species-specific (human, mouse or rat depending on the experiment) repetitive elements from RepBase 18.05 using STAR (v2.7.6a) using the following parameter: --outFilterMultimapNmax 30. Reads that did not align to the repeats database were then mapped to the human (hg38), mouse (mm10), or rat (rn6) genomes using STAR (v2.7.6a) (Dobin et al., 2013) with options to require unique mapping (-- outFilterMultimapNmax=1) and end-to-end read alignment was forced (-- alignEndsType=EndToEnd). PCR duplicates were removed using umi_tools (v1.0.0) by utilizing the 5’ UMI sequences and mapping positions. Clusters of reads were identified within eCLIP samples using the cluster caller CLIPper (https://github.com/YeoLab/clipper/commit/61d5456) (Lovci *et al*., 2013) using transcript annotations for human (GENCODE v29; ENCODE accession ID ENCFF159KBI), mouse (GENCODE vM25), or rat (ENSEMBL release 98). For each cluster IP versus input fold enrichment values were then calculated as a ratio of counts of reads overlapping the cluster region in the IP and the input samples (read counts in each sample were normalized against the total number of genome mapped reads in the sample remaining after PCR duplicate removal). A p-value was calculated for each cluster by the Yates’ Chi-Square test (or Fisher Exact Test if the observed or expected read number was below 5). A pseudocount of 1 read was added to all read counts per cluster for input samples when calculating p-values and IP versus input log2 fold changes. Clusters were filtered using a list of excluded regions (ENCODE accession ID ENCFF269URO) and annotated using transcript information from GENCODE v29 (ENCODE accession ID ENCFF159KBI) and LNCipedia v5.0 (Volders et al., 2019) for human and vM25 for mouse (Frankish et al., 2019) using the following priority hierarchy to define the final annotation of overlapping features: protein coding transcript (CDS, UTRs, intron), followed by non-coding transcripts (exon, intron). Clusters passing cutoffs of IP vs. input fold enrichment ≥ 8 and p-value ≤ 0.001 were deemed significant and referred to as significantly enriched peaks.

Information content for significant peaks was calculated using the following formula, where *c_i_* = number of IP reads overlapping the peak, *i_i_* = number of input reads overlapping the peak, *n_i_* = total IP reads, and *m_i_* = total input reads.

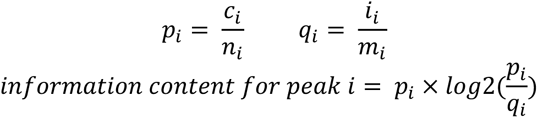

Inforrmation content was summed across peaks to calculate the total information at peaks overlapping various transcript annotation regions.

### Comparison of no-gel versus with-gel fold-enrichment

Peak-level fold-enrichment between samples was calculated by first taking all CLIPper clusters in the first (e.g. no-gel) sample and then calculating the fold-enrichment in IP versus input separate for each cluster in both samples (with-gel and no-gel), with a pseudocount of 1 added to the input read counts at each cluster. For the second (e.g. with-gel) sample in the comparison, if no reads in the IP overlapped the cluster, a pseudocount of 1 was used. The fold-enrichments for all clusters across both experiments were then plotted (Fig. 2E). This process was repeated for the inverse comparison (e.g. comparing fold-enrichment for all clusters identified in the with-gel experiment) (Sup. Fig. 2F).

### Chimeric read analysis

Identification of chimeric reads was adapted based on the method described by Moore et al. (Moore *et al*., 2015). Reads were first “reverse mapped” to a database of mature human or mouse miRNAs compiled from miRBase (v22) (Kozomara et al., 2019) by mapping the miRNA sequences to the reads using Bowtie (v1.2.2) (Langmead et al., 2009). Alignments of miRNA to sequencing reads were then filtered, keeping only positive strand alignments and prioritizing alignments with the fewest number of mismatches so that only one miRNA alignment was selected for each read. In each of these reads, sequences flanking the miRNA alignment were identified (the chimeric “target” portion in chimeric reads), and reads with target portions less than 18 nucleotides in length were discarded. As above, target portions were first mapped to a species-specific repeat element database, with mapped reads discarded. The remaining sequences were mapped to the human (hg38), mouse (mm10), or rat (rn6) genomes using STAR (v2.7.6a) and PCR duplicates were removed using umi_tools (v1.0.0). Annotation of target portions of chimeric reads was performed using transcript information for human (GENCODE v29), mouse (GENCODE vM25) or rat (RefGene) following the same hierarchy rules as listed for the annotation of peaks listed above.

For initial analyses, clusters were identified with CLIPper using all chimeric reads as input (with transcript annotations described above); for following analyses, chimeric reads were first separated by miRNA, and clusters were then identified (CLIPper) separately for each miRNA using the miRNA-specific set of chimeric reads. Reproducible clusters were identified with an approach based off the Irreproducible Discovery Rate (IDR) approach (Li et al., 2011) by ranking all clusters based on -log10 transformed CLIPper p-values, followed by running the IDR software (v 2.0.2). IDR-identified regions with IDR score ≥ 540 were used as ‘reproducible’ clusters.

### Calculation of miRNA and chimeric read abundance

As some miRNAs have multiple highly homologous genomic copies, quantification of miRNA abundance from only uniquely mapped reads yielded inaccurate estimates of expression. To address this for non-chimeric analyses, multimapping reads were also considered by taking reads that failed to map uniquely and redoing STAR mapping with the option ‘-- outFilterMultimapNmax=10000’ flag, then selecting the primary alignment for each read, and then performing PCR duplicate removal as described above for uniquely mapping reads. This approach may miss occasional PCR duplicates that are assigned different primary alignments from multiple equal alignments; analysis indicates that this typically alters expression estimates by <2%.

As appropriate, per-miRNA and per-cluster abundance was normalized using three alternative approaches: RPM (reads per million) calculated versus all uniquely mapped, non-PCR duplicate reads, RPMI (Reads Per Million Initial) calculated versus all sequenced reads (prior to any adapter trimming or other processing), and RPMC (Reads Per Million Chimeras) calculated versus all uniquely mapped, non-PCR duplicate chimeric reads. RPM of chimeric reads was calculated with a denominator of the sum of (uniquely mapped, PCR duplicate removed) chimeric reads and non-chimeric reads. RPM normalization of miRNA read counts in small RNA-seq was performed analogously to the calculation of non-chimeric RPM in chimeric eCLIP experiments.

### Analysis of human/rat mixing experiments

For human/rat mixing experiments, adapter trimming, mapping to human mature miRNAs (miRBase v22), removal of putative rRNA artifact miRNAs (Supplementary Table 1), and identification of potential chimeric reads was performed as described above (in pilot analyses, mapping to human miRNAs gave similar results as separate mapping to both human and rat miRNA annotations). After removing the miRNA region, the remaining putative chimeric fractions were first mapped to a repeat element database containing both human and rat elements (RepBase 18.05) with mapped fragments discarded. Next, remaining reads were separately mapped to human (hg38) and rat (rn6) genomes. For this analysis, genomic mapping was performed allowing multiple mapping (--outFilterMultimapNmax 99). Mapping was then compared between human and rat using STAR alignment score (‘AS’ flag), with ‘species-specific’ reads defined as those with a AS score at least 2 greater for one species than the other.

To identify species-specific miRNAs, first the number of putative chimeras (reads containing a miRNA plus at least 18nt of additional sequence either on the 5’ or 3’ end) were counted for all annotated miRNAs. Next, for the 3 with-gel pairs of human-only and rat-only chimeric eCLIP experiments, the fold-change in expression was calculated for each miRNA (after adding a pseudocount of 1 to both human and rat counts and normalizing against the total number for all miRNAs in the dataset). The set of ‘species specific’ microRNAs was then defined as those with a fold-change greater than or equal to 10 in all three human-only/rat-only pairs. Analysis of putative false-positive chimeras was performing using only chimeras for these miRNAs for which the chimeric fraction was also a species-specific map to human or rat (defined as reads that either only mapped to one of the two genomes, or for which the STAR mapping score (AS flag) was at least 2 larger for one species than the other). Annotations for rat transcripts used RefGene annotations from the UCSC Genome Browser (obtained 10/28/2019).

### Analysis of PCR-enriched chimeric eCLIP

DNA fragments in the targeted chimeric eCLIP libraries contain a 10nt UMI 3′ end of the insert sequence. These UMIs were pruned from the 5′ ends of R2 read sequences and saved by incorporating them into the read names in the R1 FASTQ files to be utilized in subsequent analysis steps. All subsequent steps were performed on R1 FASTQ files only. Next, 3′-adapters were trimmed from reads using cutadapt (v2.5) (Martin, 2011), and remaining reads than 18 nucleotides in length were removed. The final 9 nucleotides at the 3′ end of each read were trimmed using cutadapt (v2.5) to remove potential UMI sequence at the 3′ end. The miRNA primer sequence used to select for the miRNA of interest was then used to select for chimeras and trimmed from the 5′ end of reads, and reads with remaining length shorter than 18 nucleotides were removed. Reads were then processed using the same steps as in the “Non-chimeric reads” section above.

### Motif analysis of chimeric eCLIP clusters

First, each cluster was extended by 10nt on the 5’ side (to account for possible clusters that terminate at the protein-RNA crosslink site), and sequences were obtained from the appropriate species genome. Next, the presence or absence of every 6-mer was calculated for each extended cluster sequence, and percent frequency for each 6-mer was determined across either all clusters or subsets of clusters that met significance (CLIPper *p* ≤ 10^-5^ or IDR ≥ 540) as indicated. Seed 6- mer sequences complementary to miRNA positions 2-7, 3-8, 1-6, or A1-6 (an A at position 1 followed by miRNA positions 2-6) were obtained from the major annoted miRNA sequence from miRbase.

### RNA-seq analysis

Reads in the RNA-seq libraries contain a 10 nucleotide UMI at the 5′ end of each read, which was pruned from 5′ ends of R1 read sequences using umi_tools (v1.0.1). Next, 3′-adapters were trimmed from reads using cutadapt (v2.7), discarding reads less than 18 nucleotides remaining. Next, reads were mapped to a database of species-specific repetitive elements, as described above in methods for Non-chimeric (standard eCLIP) analysis. Reads that did not align to the repeats database were then mapped to the human genome (hg38) using STAR (v2.7.6a). PCR duplicates were removed using umi_tools (v1.0.0) by utilizing UMI sequences from the read names and mapping positions. Gene counts per sample were obtained using previously described pipelines for quantifying region-level coverage from eCLIP data (Van Nostrand *et al*., 2020a) as well as transcript information from GENCODE (V29). Differential expression analysis was performed using the R package DESeq2 (v 1.34.0) (Love *et al*., 2014).

### sRNA-seq analysis

Reads in sRNA-seq libraries contain the sequence “AACTGTAGGCACCATCAAT” followed by a 12 nucleotide UMI at the 3′ end of each read. The “AACTGTAGGCACCATCAAT” sequence was identified within each read using cutadapt (v2.7) and reads that did not contain this sequence were discarded. Next, the UMIs were appended to the read names, and the “AACTGTAGGCACCATCAAT” sequence as well as following 3′ sequence (this includes the UMI and sequencing adapter) using a custom python script. Reads were mapped to a database of human repetitive elements and rRNA sequences compiled from Dfam and Genbank. Reads that did not align to the repeat database were then mapped uniquely to the human genome (hg38) using STAR (v2.6.0c). PCR duplicates were removed using umi_tools (v1.0.1) by considering both UMI sequences and mapping positions. miRNA counts were obtained using a custom python script and miRNA annotations from Mirbase (v22). In order to allow relative comparison of abundances of miRNAs that are transcribed from multiple genomic locations, reads that failed to map uniquely were re-mapped allowing multimapping and selecting primary alignment as described above for miRNA quantification with non-chimeric reads.

### miRNA seed match analysis for miRNA over-expression

To assess frequency of miRNA seed matches in 3’ UTR of up- and down-regulated genes following miRNA transfections, a seed match was defined as presence of 6-mers complementing miR-124 or miR-1 mature sequences in positions [2:7] or [3:8]. 3’ UTR sequences were obtained by merging overlapping and bookending regions with 3’ UTR annotations into one contig encompassing all 3’ UTR sequences associated with a gene (with N characters inserted when joining non-adjacent sequences in order to avoid creation of subsequences not present in a real isoform). Next, genes were sorted based on the DESeq2 ‘stat’ value, and the Sylamer tool (van Dongen et al., 2008) was used to calculate a hypergeometric enrichment p-value for each 6-mer in growing bins that increase size in steps from the beginning of the list to the end of the list, using k-mer size of 6 nt (-k 6), bin growth step size of 50 genes (-grow 50) and making evaluation of 6- mer counts conditional on frequency of k-mers of up to 4 nt in length (-m 4).

### Comparison of miRNA targets with miRNA over-expression

To connect information about miRNA target sites with changes in the transcriptome, each gene was assessed for presence of miRNA-specific chimeric clusters identified in three non-overlapping features (3’ UTR, CDS or intronic) defined based on GENCODE (v29) annotations and feature hierarchy as described above in “Non-chimeric (standard eCLIP) analysis”. When miRNA-specific chimeric clusters overlapping a gene were identified, the average chimeric read coverage of the cluster (calculated by taking the mean of two biological replicates) was used to describe chimeric coverage of the cluster. If more than one cluster was present in the same annotation type of the gene, then the site with the greatest chimeric coverage was used. Cutoffs for different analyses were implemented as indicated, typically using genes with either all reproducible clusters (chimeric RPMC ≥ 0 and IDR ≥ 540), or subsets of these with increased chimeric RPMC cutoffs. For analysis of ‘probe-enriched only versus total and probe-enriched’ targets (Sup. Fig. 7J-K), ‘total and probe-enriched’ were defined as genes with clusters with chimeric RPMC ≥ 100 and IDR ≥ 540 in both probe-capture and total (non-enriched) experiments, and ‘probe-enriched only’ were all other genes with clusters with chimeric RPMC ≥ 100 and IDR ≥ 540 in the probe-capture experiment. For analysis of CDS versus 3’ UTR and intronic targets, genes were included in the ‘CDS only’ class if they contained a chimeric cluster (chimeric RPMC ≥ 100 and IDR ≥ 540) in the CDS region and lacked any clusters for both 3’ UTR and intronic regions (chimeric RPMC ≥ 0) (with equivalent rules for ‘3’ UTR only’ and ‘Intronic only’).

In addition to annotating genes based on chimeric eCLIP target sites in three different features (3’ UTR, CDS and intronic features), genes were also gropued based on predictions of 896 miR- 1-3p targets and 1,820 miR-124-3p targets by TargetScan version 7.2 (Agarwal *et al*., 2015), of which 862 of miR-1-3p and 1,769 of miR-124-3p targets overlapped GENCODE v29 gene identifiers. For analysis of overlap between TargetScan and chimeric eCLIP data, genes were first separated based on presence or absence of TargetScan predicted targets, and then based on whether they contained a cluster (chimeric RPMC ≥ 100 and IDR ≥ 540) in the probe-capture chimeric experiment.

### Published CLASH/chimeric CLIP datasets

For comparison (Sup. Fig. 7a), the number of miRNA chimeras were taken from prior publications and not re-processed.

## Supporting information

Supplemental Protocol

## Materials and Data Availability

Further information and requests for resources and reagents should be directed to and will be fulfilled by the Lead Contact, Eric Van Nostrand. This study did not generate new unique reagents. Chimeric eCLIP data has been deposited at the Gene Expression Omnibus (GEO ID GSE198251 ; reviewer token yruvagicbjixnin). The primary data processing pipeline for chimeric eCLIP data is available at https://github.com/YeoLab/chim-eCLIP. Other custom scripts are available upon reasonable request.

## Author Contributions

Conceptualization, A.A.S., G.W.Y and E.L.V.N.

Methodology, A.A.S. and E.L.V.N.

Investigation, S.A.M., A.A.S., B.A.Y., K.A.S., D.C.C., S.S.P., H.M.F., and E.L.V.N.

Software and Formal Analysis, S.A.M., B.A.Y., K.A.S., and E.L.V.N.

Writing – Original Draft, S.A.M. and E.L.V.N.

Writing – Review & Editing, S.A.M., A.A.S., B.A.Y., K.B.C., G.W.Y., and E.L.V.N.

Supervision, A.A.S., K.B.C, G.W.Y, and E.L.V.N.

Project Administration, A.A.S., K.B.C, G.W.Y, and E.L.V.N.

Funding Acquisition, A.A.S., K.B.C, G.W.Y, and E.L.V.N.

## Supplementary Files

**Supplemental Protocol 1** – Detailed protocol for chimeric eCLIP experiments described here

## Acknowledgements

We thank Jeanine Van Nostrand and Fernando Martinez for assistance with dissection of mouse liver tissue, and members of the Van Nostrand and Yeo labs for ongoing feedback during this work (particularly S. Aigner and M. Akinyi). GWY is supported by NIH grants U41 HG009889, U01 HG009417, R01 HL137223, R01 HG004659. ELVN is a CPRIT Scholar in Cancer Research (RR200040) and is supported by NHGRI (R00HG009530). AAS and KAS were supported by NHGRI (R43HG010603).

---

**Supplemental Figure 1.**
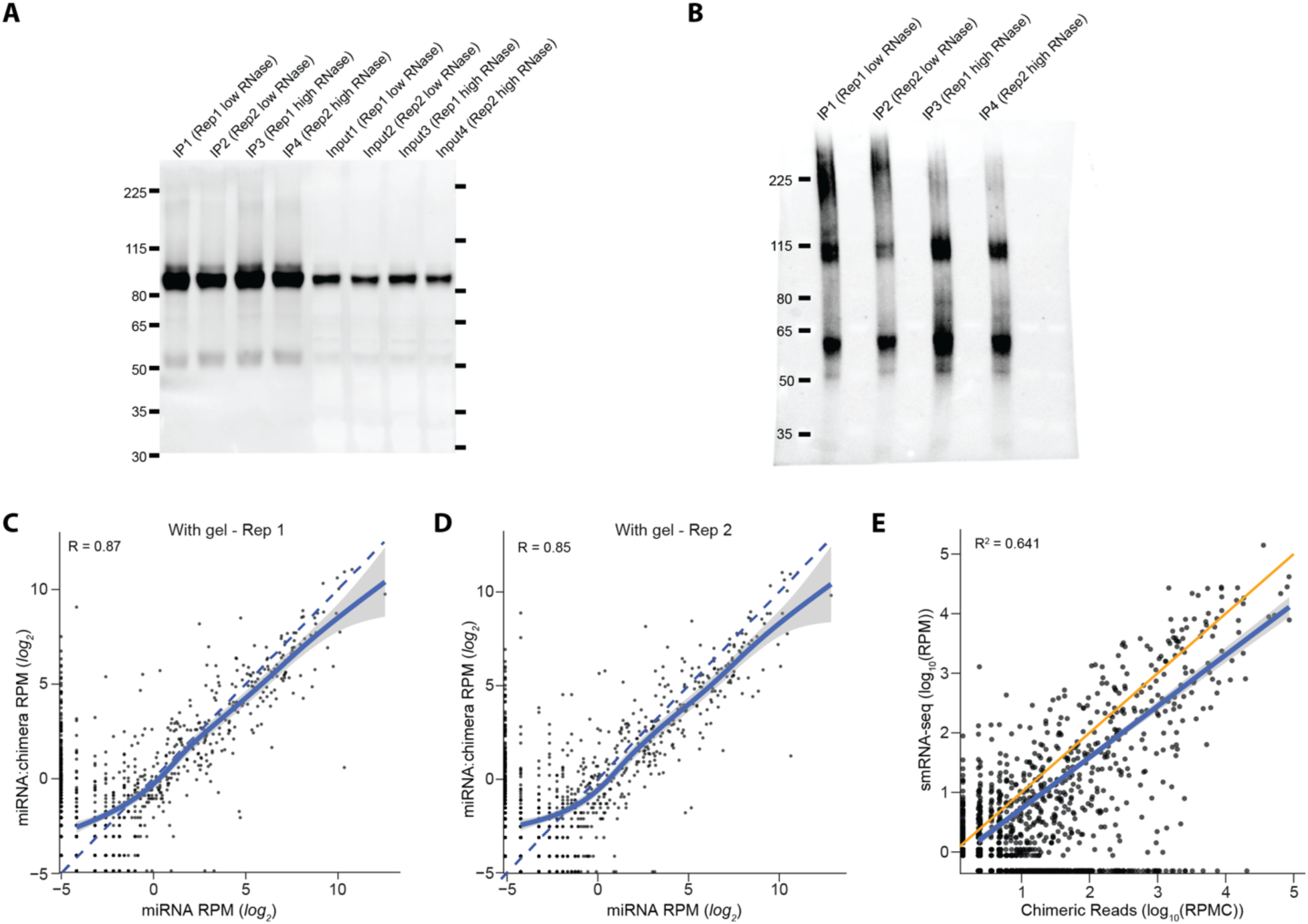
Identification of miRNA targets with chimeric eCLIP. (A-B) ImmunoPrecipitation (IP) Western Blot and RNA visualization for AGO2 chimeric eCLIP. Shown are low (3.3U) and high (100U) RNase conditions. (A) Western blot using anti-AGO2 primary antibody (Sino Biological) (B) RNA visualization blot for IP samples from (A). Biotin on the ligated RNA adapter was visualized with a biotin Chemiluminescent Nucleic Acid Detection Module Kit (Thermo Scientific). (C-D) For each miRNA, points indicate reads per million (RPM) of (x-axis) non-chimeric (miRNA-only) reads versus (y-axis) chimeric reads containing the miRNA per million mapped reads (RPM). The dashed line shows equality, and the blue line shows a generalized additive model (GAM) fit obtained using the geom_smooth function of R package ggplot2 (v3.3.5) with 0.95 confidence intervals in grey. Panels show data from (C) replicate 1 and (D) replicate 2. (E) Each point corresponds to a miRNA with values on x-axis showing mean chimeric reads per million chimeras (RPMC) from two replicates of with-gel chimeric eCLIP and values on y-axis showing mean reads per million (RPM) from two replicates of small RNA-seq HEK293T libraries. The orange line shows equality, and the blue line shows least squares linear fit with 0.95 confidence intervals.

**Supplementary Figure 2.**
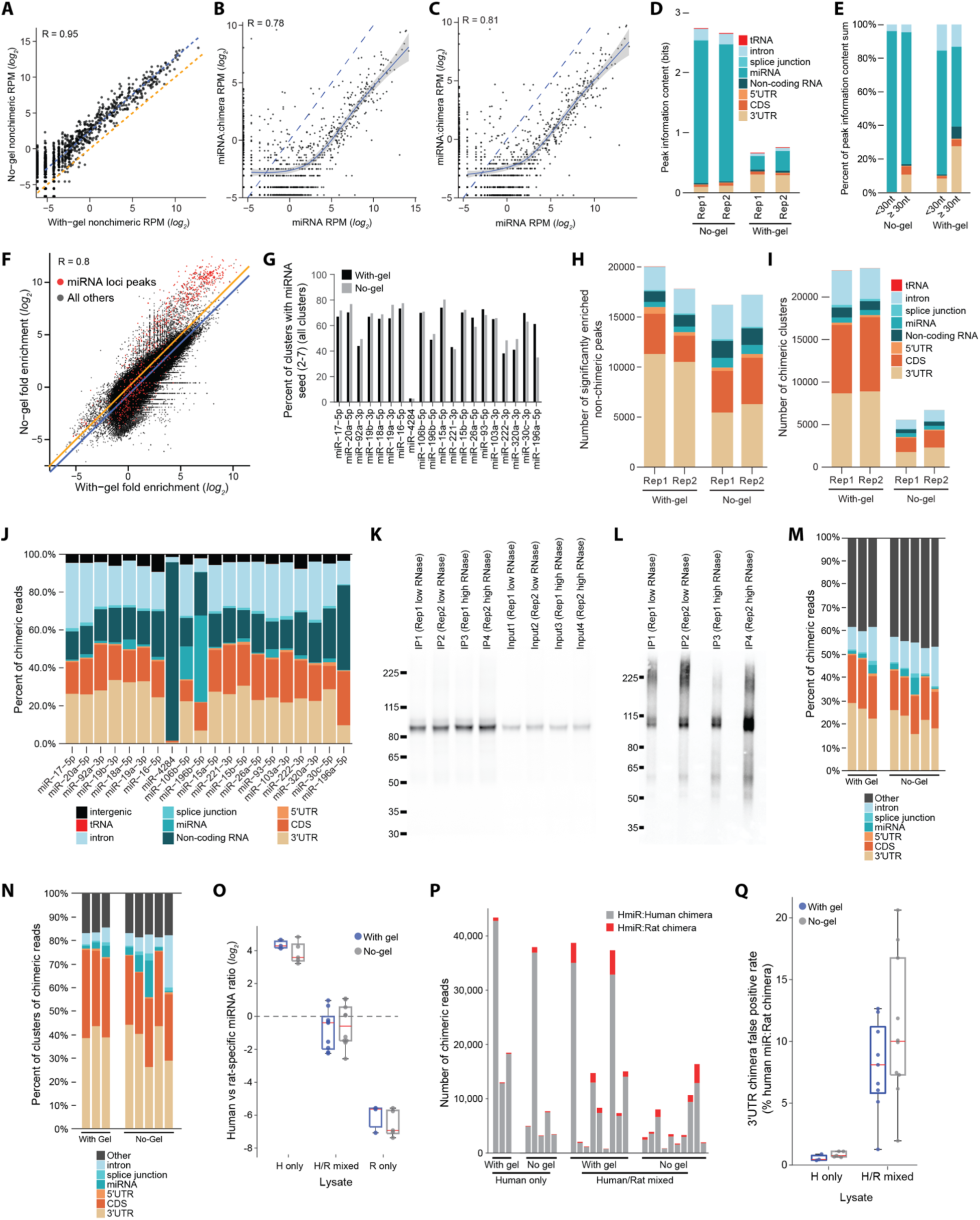
Improved recovery of chimeric miRNA fragments with no-gel chimeric eCLIP. (A) Points indicate miRNA abundance among non-chimeric reads for standard with-gel (x-axis) versus no-gel (y- axis) experiments. Blue line indicates least squares linear regression, and orange line indicates equality. (B-C) For each miRNA, points indicate reads per million (RPM) of (x-axis) non-chimeric (miRNA-only) reads versus (y-axis) chimeric reads containing the miRNA for no-gel chimeric eCLIP experiments. The dashed line shows equality, and the blue line shows a generalized additive model (GAM) fit obtained using the geom_smooth function of R package ggplot2 (v3.3.5) with 0.95 confidence intervals in grey. Panels show data from (B) replicate 1 and (C) replicate 2. (D-E) Chimeric eCLIP experiments in HEK293T cells analyzed with standard eCLIP peak calling. For significantly enriched peaks (*p* ≤ 10^-3^ and fold-enrichment ≥ 8 in immunoprecipitation (IP) versus size-matched input), stacked bars indicate (D) the sum of information content in significantly enriched peaks or (E) the percent of the total sum of information content, with peaks annotated based on overlapping transcript regions (indicated with colors). (F) For each of 353,280 clusters identified from non-chimeric eCLIP analysis of with-gel chimeric eCLIP replicate 1, points indicate fold-enrichment in (x-axis) with-gel chimeric eCLIP replicate 1 versus (y-axis) no-gel chimeric eCLIP replicate 1, with Pearson correlation indicated. Points indicated in red reflect peaks overlapping miRNA loci. (G) Bars indicate the percent of clusters containing miRNA 2-7 seed (the 6-mer complementary to each miRNA 2-7 position). Shown are the most abundant 20 miRNAs as in Fig. 2F. (H) Stacked bars indicate the number of significantly enriched peaks (*p* ≤ 10^-3^ and fold-enrichment ≥ 8 in immunoprecipitation (IP) versus size-matched input) from standard eCLIP (non-chimeric) analysis of chimeric eCLIP datasets. (I) Stacked bars indicate number of clusters identified from chimeric reads only, with overlapping transcript annotation indicated by colors. (J) Stacked bars indicate percent of all chimeric reads for the top 20 most abundant miRNAs that overlap indicated transcript annotations. (K-L) ImmunoPrecipitation (IP) Western Blot and RNA visualization for AGO2 chimeric eCLIP in rat C6 cells. Shown are low (3.3U) and high (100U) RNase conditions. (K) Western blot using anti-AGO2 primary antibody (Sino Biological) (L) RNA visualization blot for IP samples from (K). Biotin on the ligated RNA adapter was visualized with a biotin Chemiluminescent Nucleic Acid Detection Module Kit (Thermo Scientific). (M-N) Stacked bars indicate the percent of (M) chimeric reads or (N) clusters called from chimeric reads from with- and no-gel chimeric eCLIP experiments performed in rat C6 cells that overlap with the indicated transcript regions. (O) Points indicate the log_2_(ratio) of H (human HEK293T-specific miRNAs, defined as RPM_HEK293T_ / RPM_C6_ ≥ 10) versus R (rat C6 specific miRNAs, defined as RPM_C6_ / RPM_HEK293T_ ≥ 10). Standard (with-gel) experiments are indicated in blue and no-gel experiments are indicated in grey. Red line indicates median, and boxes indicate 25^th^ to 75^th^ percentiles. (P) For all replicates of human-only and human/rat mixed samples, stacked bars indicate the number of chimeric reads for human-specific miRNAs for which the chimeric region uniquely maps to (grey) human or (red) rat protein-coding transcriptomes (including mapping to 5’UTR, CDS, 3’UTR, and intronic regions). (Q) Points indicate ‘false positive’ rate per mixing experiment calculated from 3’UTR-mapping chimeras only, determined by identifying the subset of chimeras where the target fragment maps uniquely either to human or rat and calculating the ratio of miRNA_Human_:target_Rat_ target chimeras divided by miRNA_Human_:target_Human_ + miRNA_Human_:target_Rat_. Standard (with-gel) experiments are indicated in blue and no-gel experiments are indicated in grey. Red line indicates median, and boxes indicate 25^th^ to 75^th^ percentiles.

**Supplementary Figure 3.**
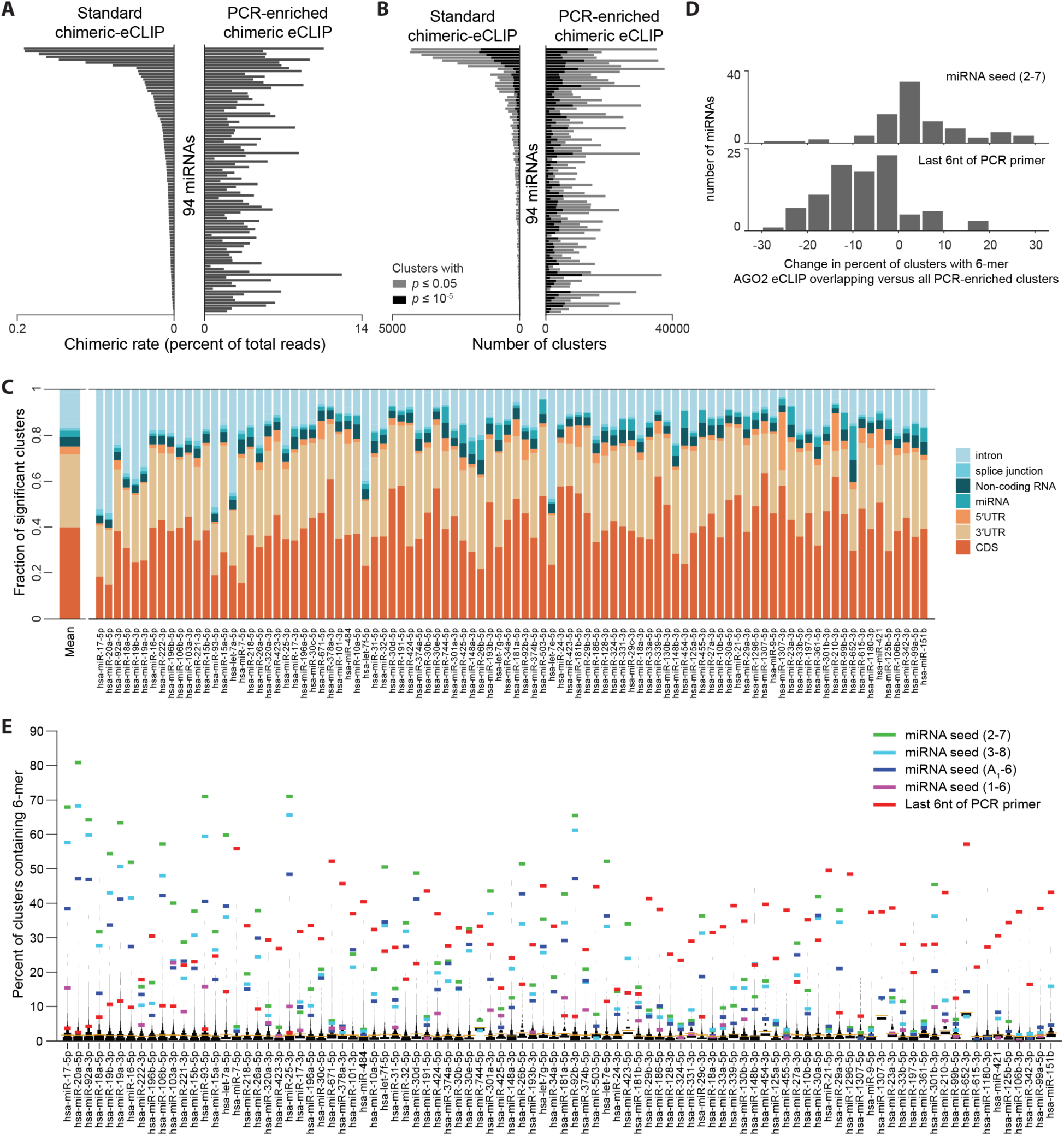
PCR enrichment of miRNAs of interest. (A) Bars indicate the percent of sequenced reads that are chimeric (containing both a miRNA and a uniquely mapped other fragment) for the 94 most abundant miRNAs profiled by (right) PCR-enriched chimeric eCLIP or (left) standard with-gel chimeric eCLIP in HEK293T cells. (B) Stacked bars indicate the number of (black) *p*≤10^-5^ or (grey) *p*<0.05 CLIPper-identified clusters using chimeric reads only for the 94 miRNAs. (C) From chimeric reads from PCR-enriched chimeric eCLIP for the 94 miRNAs, stacked bars indicate the fraction of CLIPper-identified clusters (with *p*≤10^-5^) overlapping the indicated transcript region annotations. (D) For each miRNA, the difference in the percent of clusters containing either (top) the miRNA seed (2-7) or (bottom) the last 6nt of the PCR primer was calculated for all clusters versus only those clusters which overlap significantly enriched (versus paired input) peaks from standard (non-chimeric) analysis of with-gel AGO2 chimeric eCLIP, with the histogram showing the distribution of these differences. (E) For each of 94 miRNAs profiled by PCR-enriched chimeric eCLIP, lines indicate the percent of significant clusters identified from chimeric reads containing either (colors) the indicated five 6-mers, or (black histogram) all other 6-mers. Orange line indicates mean of all other 6-mers. For this figure, only clusters that overlap significantly enriched peaks from standard (non-chimeric) analysis of with-gel AGO2 chimeric eCLIP were considered (see Fig. 3E for analysis of all clusters).

**Supplementary Figure 4.**
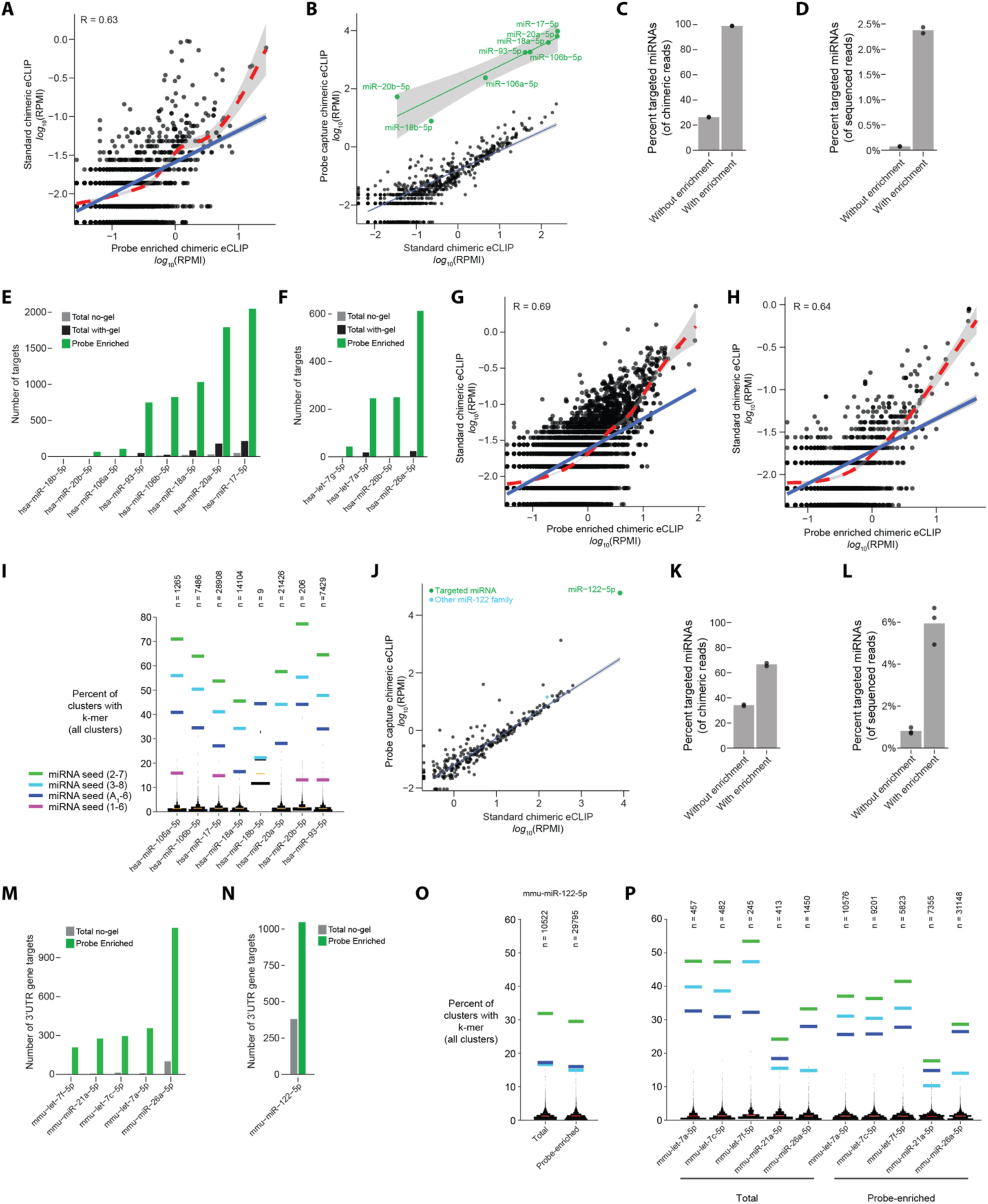
Probe-capture enrichment of miRNAs of interest. (A) Points indicate miR-221 chimeric reads in Reads Per Million Initial (RPMI) from (x-axis) probe-enriched or (y-axis) standard chimeric eCLIP for all clusters identified in either dataset (using one replicate for each). Red line indicates smoothed conditional mean predicted by a generalized additive model (GAM) fitted using geom_smooth function of ggplot2 (v3.3.5) package in R and blue line indicates an ordinary least squares model, with grey region indicating confidence intervals (0.95 level). (B) Points indicate chimeric reads in Reads Per Million Initial (RPMI) for all miRNAs in (x-axis) standard chimeric eCLIP, or (y-axis) probe capture-enriched chimeric eCLIP. Green points and line indicate targeted miRNAs with a least squares linear model fit showing 0.95 confidence intervals. Black points and a robust linear regression model fit showing 0.95 confidence intervals represent all other miRNAs. (C-D) Points indicate percent of chimeric reads out of (C) all chimeric reads or (D) all sequenced reads that represent the targeted miRNAs from (B), with bar indicating mean of two replicate experiments. (E-F) Bars indicate the number of genes with reproducible clusters in their 3’UTR regions for the indicated enriched miRNAs, for (E) the 8- miRNA enrichment experiment or (F) the 4-miRNA experiment (See Fig. 4H-J). (G-H) Points indicate (G) miR-17 or (H) miR-26a chimeric reads in Reads Per Million Initial (RPMI) from (x- axis) probe-enriched or (y-axis) standard chimeric eCLIP for all clusters identified in either dataset (using one replicate for each). Red line indicates smoothed conditional mean predicted by a generalized additive model (GAM) fitted using geom_smooth function of ggplot2 (v3.3.5) package in R and blue line indicates an ordinary least squares model, with grey region indicating confidence intervals (0.95 level). (I) For each of the targeted miRNAs, lines indicate the percent of clusters of probe capture enriched chimeric reads containing either (colors) the indicated five 6-mers, or (black histogram) all other 6-mers. Orange line indicates mean of all other 6-mers. Data shown are from one representative replicate. (J-L,N) miR-122-5p capture enrichment in mouse liver tissue. (J) Points indicate chimeric reads in Reads Per Million Initial (RPMI) for all miRNAs in (x-axis) standard chimeric eCLIP, or (y-axis) probe capture-enriched chimeric eCLIP performed in mouse liver. Green indicates miR-122-5p, and light blue indicates miRNAs that have the same seed as miR-122-5p but were not specifically targeted. Black points and a robust linear regression model fit showing 0.95 confidence intervals represent all other miRNAs except those with fewer than 20 chimeric reads in standard chimeric eCLIP experiment. (K-L) Points indicate percent of chimeric reads out of (K) all chimeric reads or (L) all sequenced reads that that represent the targeted miRNAs from (J), with bar indicating average from two replicate experiments. (M-N) As in (D-E) for (M) miR-122 enrichment experiment or (N) 5 miRNA enrichment experiment. (O-P) Motif frequency as in (I) shown for (O) miR-122 enrichment experiment or (P) 5 miRNA enrichment experiment (see Fig. 4K-L).

**Supplemental Figure 6.**
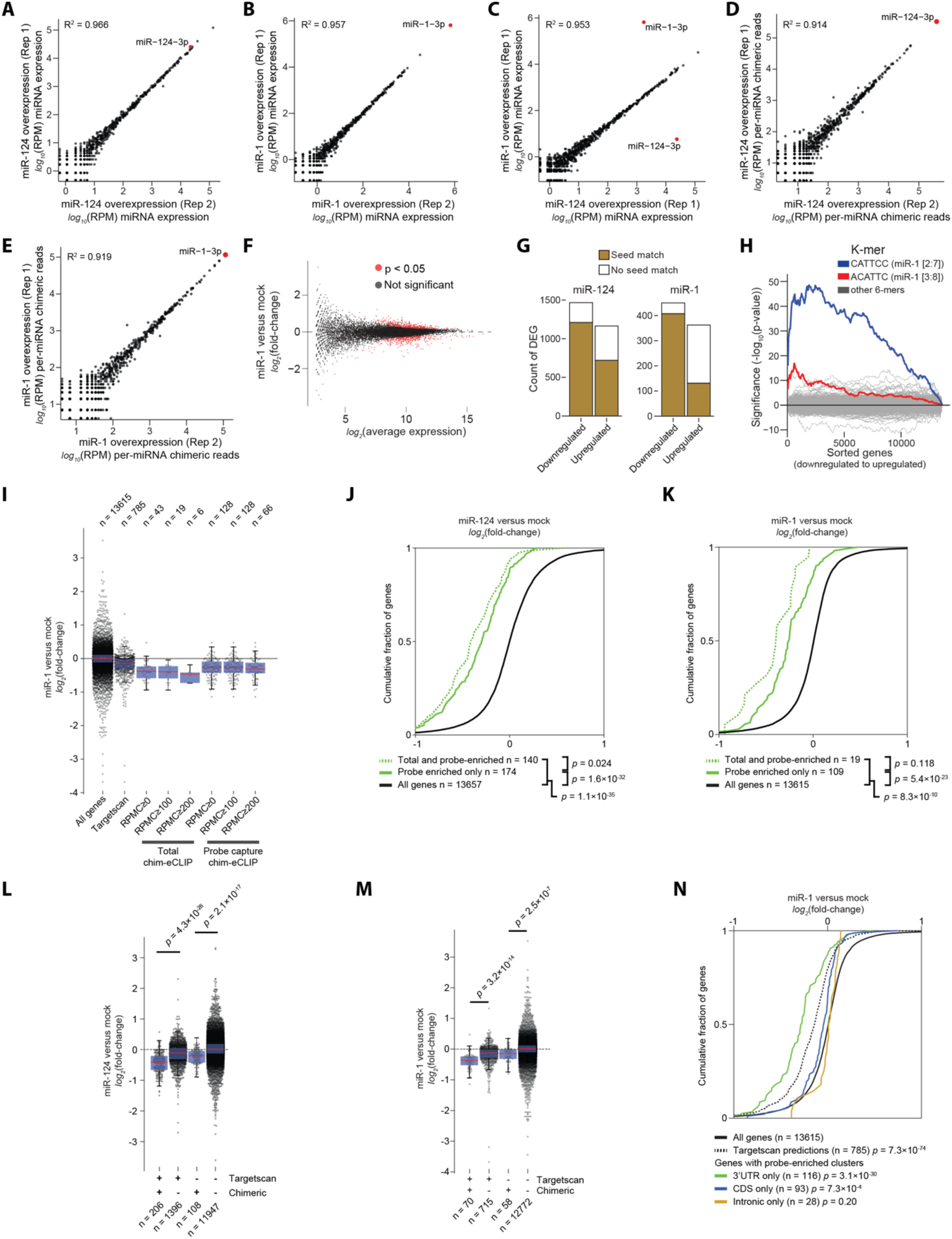
Validation of regulated miRNA targets with miRNA over-expression. (A-C) Points indicate per-miRNA abundance from small RNA-seq after miR over-expression. Shown are (A) two miR-124 replicates, (B) two miR-1 replicates, and (C) one replicate of miR-124 over-expression versus miR-1 over-expression. (D-E) Points indicate per-miRNA abundance within chimeric reads for chimeric eCLIP experiments performed after (D) two replicates of miR-124 over-expression or (E) two replicates of miR-1 over-expression. (F) Points indicate gene (x-axis) average expression (log_2_(DESeq2 normalized read counts)) and (y-axis) fold-change in miR-1 over- expression versus mock transfection, with significant differential expression indicated in red (DESeq2 p-value < 0.05). (G) Stacked bars indicate number of differentially expressed genes upon (left) miR-124 or (right) miR-1 over-expression, separated by whether they contain a match against the miRNA seed region (positions [2-7]). (H) Lines show -log_10_ transformed hypergeometric p-values of enrichment of each of 2,332 6-mers complementing miRBase human miRNAs in [2:7] and [3:8] positions in growing bins of 3’UTR sequences of all expressed genes. Genes were sorted from down- to up-regulated upon miR-1 over-expression using test statistic associated with expression fold-change. (I) For indicated gene classes, points indicate fold-enrichment in miR-1 over-expression. Red line indicates median, and blue boxes indicate 25^th^ to 75^th^ percentiles. Gene classes include those with TargetScan7.2-predicted miR-1 targets and genes containing chimeric eCLIP clusters meeting indicated reads per million (RPM) cutoff (average across two replicates) for either total (non-enriched) or miR-1 probe-enriched chimeric eCLIP. Plots were generated using the MATLAB ‘plotSpread’ package. (J-K) Cumulative empirical distribution plots show fold-enrichment in miRNA over-expression for (J) miRNA-124 and (K) miRNA-1. Lines indicate sets of genes (green, dotted line) with reproducible chimeric clusters (using a cutoff of average chimeric read RPMC≥100 and IDR > 540) from both probe- enriched and total chimeric eCLIP, (green, solid line) with clusters in probe-enriched eCLIP only, or (black) all genes. Significance was determined by two-tailed Kolmogorov-Smirnov test. (L-M) For indicated gene classes, points indicate fold-enrichment in (L) miR-124 or (M) miR-1 over- expression. Red line indicates median, and blue boxes indicate 25^th^ to 75^th^ percentiles. Gene classes include sets of genes with or without TargetScan7.2-predicted miRNA interaction sites versus sets of genes with or without clusters from chimeric eCLIP (using a cutoff of average RPMC≥100 and IDR > 540). Significance was determined by two-tailed Kolmogorov-Smirnov test. (N) Cumulative empirical distribution plots show fold-enrichment in miRNA over-expression for miRNA-1. Significance was determined by two-tailed Kolmogorov-Smirnov test. Lines indicate sets of genes with chimeric miR-1 clusters only in (green) 3’UTR, (blue) coding sequence, or (orange) intronic regions, versus (black) all genes or (dotted black) all genes with TargetScan7.2 predicted miR-1 targets.

**Supplementary Figure 7.**
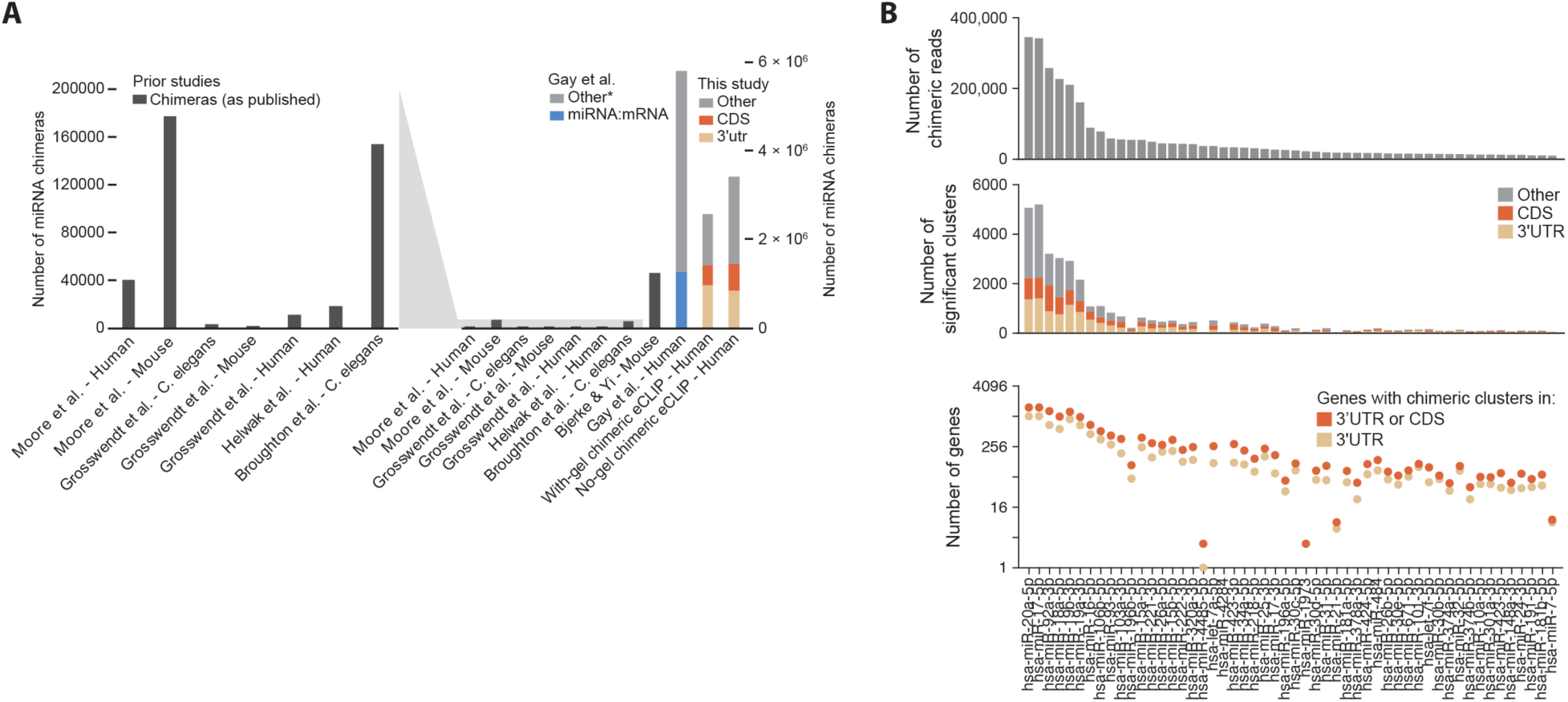
Deep profiling of miRNA interactomes in HEK293T. (A) Bars indicate the number of chimeric reads identified in prior chimeric CLIP publications or in this study, with zoomed in axis as indicated. Prior publications reflect the total number of chimeras as described in the publication. For Gay *et al*. (Gay *et al*., 2018), ‘Other’ includes both non-miRNA chimeras (e.g. mRNA:mRNA, rRNA, etc.) as well as miRNA:non-mRNA (e.g., intronic) chimeras. (B) Properties of the merged no-gel chimeric eCLIP data for the top 52 miRNAs (which each had more than 10,000 chimeric reads). (top) Bar indicates number of chimeric reads for each miRNA. (middle) Stacked bar indicates the number of significant (*p* ≤ 10^-5^) CLIPper clusters from chimeric reads for each miRNA, separated by overlapping annotation region. (bottom) On logarithmic scale, points indicate the number of unique genes containing significant clusters in either (tan) 3’UTR regions only or (orange) 3’UTR or CDS regions.

## Supplementary Tables

**Supplemental Table 1** – List of 15 miRBase annotated human miRNAs removed from analysis as rRNA artifacts

**Supplemental Table 2** – Expression of human miRNAs from small RNA-seq in HEK293T cells

**Supplemental Table 3** – List of HEK293T chimeric eCLIP datasets

**Supplemental Table 4** – Experimental details for datasets utilized in this study

**Supplemental Table 5** – Oligonucleotides utilized in chimeric eCLIP

## References

Agarwal, V., Bell, G.W., Nam, J.W., and Bartel, D.P. (2015). Predicting effective microRNA target sites in mammalian mRNAs. Elife 4. 10.7554/eLife.05005.

Ameres, S.L., and Zamore, P.D. (2013). Diversifying microRNA sequence and function. Nat Rev Mol Cell Biol 14, 475–488. 10.1038/nrm3611.

Baek, D., Villen, J., Shin, C., Camargo, F.D., Gygi, S.P., and Bartel, D.P. (2008). The impact of microRNAs on protein output. Nature 455, 64–71. 10.1038/nature07242.

Bartel, D.P. (2018). Metazoan MicroRNAs. Cell 173, 20–51. 10.1016/j.cell.2018.03.006.

Becker, W.R., Ober-Reynolds, B., Jouravleva, K., Jolly, S.M., Zamore, P.D., and Greenleaf, W.J. (2019). High-Throughput Analysis Reveals Rules for Target RNA Binding and Cleavage by AGO2. Mol Cell 75, 741–755 e711. 10.1016/j.molcel.2019.06.012.

Bjerke, G.A., and Yi, R. (2020). Integrated analysis of directly captured microRNA targets reveals the impact of microRNAs on mammalian transcriptome. RNA 26, 306–323. 10.1261/rna.073635.119.

Broughton, J.P., Lovci, M.T., Huang, J.L., Yeo, G.W., and Pasquinelli, A.E. (2016). Pairing beyond the Seed Supports MicroRNA Targeting Specificity. Mol Cell 64, 320–333. 10.1016/j.molcel.2016.09.004.

Chalfie, M., Horvitz, H.R., and Sulston, J.E. (1981). Mutations that lead to reiterations in the cell lineages of C. elegans. Cell 24, 59–69. 10.1016/0092-8674(81)90501-8.

Chi, S.W., Zang, J.B., Mele, A., and Darnell, R.B. (2009). Argonaute HITS-CLIP decodes microRNA-mRNA interaction maps. Nature 460, 479–486. 10.1038/nature08170.

Chu, Y., Yokota, S., Liu, J., Kilikevicius, A., Johnson, K.C., and Corey, D.R. (2021). Argonaute binding within human nuclear RNA and its impact on alternative splicing. RNA 27, 991–1003. 10.1261/rna.078707.121.

Cloonan, N., Brown, M.K., Steptoe, A.L., Wani, S., Chan, W.L., Forrest, A.R., Kolle, G., Gabrielli, B., and Grimmond, S.M. (2008). The miR-17-5p microRNA is a key regulator of the G1/S phase cell cycle transition. Genome Biol 9, R127. 10.1186/gb-2008-9-8-r127.

Cloonan, N., Wani, S., Xu, Q., Gu, J., Lea, K., Heater, S., Barbacioru, C., Steptoe, A.L., Martin, H.C., Nourbakhsh, E., et al. (2011). MicroRNAs and their isomiRs function cooperatively to target common biological pathways. Genome Biol 12, R126. 10.1186/gb-2011-12-12-r126.

DeVeale, B., Swindlehurst-Chan, J., and Blelloch, R. (2021). The roles of microRNAs in mouse development. Nat Rev Genet 22, 307–323. 10.1038/s41576-020-00309-5.

Dobin, A., Davis, C.A., Schlesinger, F., Drenkow, J., Zaleski, C., Jha, S., Batut, P., Chaisson, M., and Gingeras, T.R. (2013). STAR: ultrafast universal RNA-seq aligner. Bioinformatics 29, 15–21. 10.1093/bioinformatics/bts635.

Ebert, M.S., and Sharp, P.A. (2012). Roles for microRNAs in conferring robustness to biological processes. Cell 149, 515–524. 10.1016/j.cell.2012.04.005.

Erhard, F., Dolken, L., Jaskiewicz, L., and Zimmer, R. (2013). PARma: identification of microRNA target sites in AGO-PAR-CLIP data. Genome Biol 14, R79. 10.1186/gb-2013-14-7-r79.

Frankish, A., Diekhans, M., Ferreira, A.M., Johnson, R., Jungreis, I., Loveland, J., Mudge, J.M., Sisu, C., Wright, J., Armstrong, J., et al. (2019). GENCODE reference annotation for the human and mouse genomes. Nucleic Acids Res 47, D766–D773. 10.1093/nar/gky955.

Gangaraju, V.K., and Lin, H. (2009). MicroRNAs: key regulators of stem cells. Nat Rev Mol Cell Biol 10, 116–125. 10.1038/nrm2621.

Gay, L.A., Sethuraman, S., Thomas, M., Turner, P.C., and Renne, R. (2018). Modified Cross- Linking, Ligation, and Sequencing of Hybrids (qCLASH) Identifies Kaposi’s Sarcoma-Associated Herpesvirus MicroRNA Targets in Endothelial Cells. J Virol 92. 10.1128/JVI.02138-17.

Gebert, L.F.R., and MacRae, I.J. (2019). Regulation of microRNA function in animals. Nat Rev Mol Cell Biol 20, 21–37. 10.1038/s41580-018-0045-7.

Grimson, A., Farh, K.K., Johnston, W.K., Garrett-Engele, P., Lim, L.P., and Bartel, D.P. (2007). MicroRNA targeting specificity in mammals: determinants beyond seed pairing. Mol Cell 27, 91–105. 10.1016/j.molcel.2007.06.017.

Grosswendt, S., Filipchyk, A., Manzano, M., Klironomos, F., Schilling, M., Herzog, M., Gottwein, E., and Rajewsky, N. (2014). Unambiguous identification of miRNA:target site interactions by different types of ligation reactions. Mol Cell 54, 1042–1054. 10.1016/j.molcel.2014.03.049.

Gumienny, R., Jedlinski, D.J., Schmidt, A., Gypas, F., Martin, G., Vina-Vilaseca, A., and Zavolan, M. (2017). High-throughput identification of C/D box snoRNA targets with CLIP and RiboMeth- seq. Nucleic Acids Res 45, 2341–2353. 10.1093/nar/gkw1321.

Hafner, M., Landthaler, M., Burger, L., Khorshid, M., Hausser, J., Berninger, P., Rothballer, A., Ascano, M., Jr., Jungkamp, A.C., Munschauer, M., et al. (2010). Transcriptome-wide identification of RNA-binding protein and microRNA target sites by PAR-CLIP. Cell 141, 129–141. 10.1016/j.cell.2010.03.009.

He, L., Thomson, J.M., Hemann, M.T., Hernando-Monge, E., Mu, D., Goodson, S., Powers, S., Cordon-Cardo, C., Lowe, S.W., Hannon, G.J., and Hammond, S.M. (2005). A microRNA polycistron as a potential human oncogene. Nature 435, 828–833. 10.1038/nature03552.

Helwak, A., Kudla, G., Dudnakova, T., and Tollervey, D. (2013). Mapping the human miRNA interactome by CLASH reveals frequent noncanonical binding. Cell 153, 654–665. 10.1016/j.cell.2013.03.043.

Hodges, E., Xuan, Z., Balija, V., Kramer, M., Molla, M.N., Smith, S.W., Middle, C.M., Rodesch, M.J., Albert, T.J., Hannon, G.J., and McCombie, W.R. (2007). Genome-wide in situ exon capture for selective resequencing. Nat Genet 39, 1522–1527. 10.1038/ng.2007.42.

Ilik, I.A., Aktas, T., Maticzka, D., Backofen, R., and Akhtar, A. (2020). FLASH: ultra-fast protocol to identify RNA-protein interactions in cells. Nucleic Acids Res 48, e15. 10.1093/nar/gkz1141.

Iosub, I.A., van Nues, R.W., McKellar, S.W., Nieken, K.J., Marchioretto, M., Sy, B., Tree, J.J., Viero, G., and Granneman, S. (2020). Hfq CLASH uncovers sRNA-target interaction networks linked to nutrient availability adaptation. Elife 9. 10.7554/eLife.54655.

Khan, A.A., Betel, D., Miller, M.L., Sander, C., Leslie, C.S., and Marks, D.S. (2009). Transfection of small RNAs globally perturbs gene regulation by endogenous microRNAs. Nat Biotechnol 27, 549–555. 10.1038/nbt.1543.

Kingston, E.R., and Bartel, D.P. (2019). Global analyses of the dynamics of mammalian microRNA metabolism. Genome Res 29, 1777–1790. 10.1101/gr.251421.119.

Kivioja, T., Vaharautio, A., Karlsson, K., Bonke, M., Enge, M., Linnarsson, S., and Taipale, J. (2011). Counting absolute numbers of molecules using unique molecular identifiers. Nat Methods 9, 72–74. 10.1038/nmeth.1778.

Konig, J., Zarnack, K., Rot, G., Curk, T., Kayikci, M., Zupan, B., Turner, D.J., Luscombe, N.M., and Ule, J. (2010). iCLIP reveals the function of hnRNP particles in splicing at individual nucleotide resolution. Nat Struct Mol Biol 17, 909–915. 10.1038/nsmb.1838.

Kozomara, A., Birgaoanu, M., and Griffiths-Jones, S. (2019). miRBase: from microRNA sequences to function. Nucleic Acids Res 47, D155–D162. 10.1093/nar/gky1141.

Krek, A., Grun, D., Poy, M.N., Wolf, R., Rosenberg, L., Epstein, E.J., MacMenamin, P., da Piedade, I., Gunsalus, K.C., Stoffel, M., and Rajewsky, N. (2005). Combinatorial microRNA target predictions. Nat Genet 37, 495–500. 10.1038/ng1536.

Kudla, G., Granneman, S., Hahn, D., Beggs, J.D., and Tollervey, D. (2011). Cross-linking, ligation, and sequencing of hybrids reveals RNA-RNA interactions in yeast. Proc Natl Acad Sci U S A 108, 10010–10015. 10.1073/pnas.1017386108.

Langmead, B., Trapnell, C., Pop, M., and Salzberg, S.L. (2009). Ultrafast and memory-efficient alignment of short DNA sequences to the human genome. Genome Biol 10, R25. 10.1186/gb-2009-10-3-r25.

Lee, F.C.Y., Chakrabarti, A.M., Hänel, H., Monzón-Casanova, E., Hallegger, M., Militti, C., Capraro, F., Sadée, C., Toolan-Kerr, P., Wilkins, O., et al. (2021). An improved iCLIP protocol. bioRxiv. https://doi.org/10.1101/2021.08.27.457890.

Li, Q.H., Brown, J.B., Huang, H.Y., and Bickel, P.J. (2011). Measuring Reproducibility of High-Throughput Experiments. Ann Appl Stat 5, 1752–1779. doi:10.1214/11-AOAS466.

Li, X., Pritykin, Y., Concepcion, C.P., Lu, Y., La Rocca, G., Zhang, M., King, B., Cook, P.J., Au, Y.W., Popow, O., et al. (2020). High-Resolution In Vivo Identification of miRNA Targets by Halo-Enhanced Ago2 Pull-Down. Mol Cell 79, 167–179 e111. 10.1016/j.molcel.2020.05.009.

Lim, L.P., Lau, N.C., Garrett-Engele, P., Grimson, A., Schelter, J.M., Castle, J., Bartel, D.P., Linsley, P.S., and Johnson, J.M. (2005). Microarray analysis shows that some microRNAs downregulate large numbers of target mRNAs. Nature 433, 769–773. 10.1038/nature03315.

Lovci, M.T., Ghanem, D., Marr, H., Arnold, J., Gee, S., Parra, M., Liang, T.Y., Stark, T.J., Gehman, L.T., Hoon, S., et al. (2013). Rbfox proteins regulate alternative mRNA splicing through evolutionarily conserved RNA bridges. Nat Struct Mol Biol 20, 1434–1442. 10.1038/nsmb.2699.

Love, M.I., Huber, W., and Anders, S. (2014). Moderated estimation of fold change and dispersion for RNA-seq data with DESeq2. Genome Biol 15, 550. 10.1186/s13059-014-0550-8.

Majoros, W.H., Lekprasert, P., Mukherjee, N., Skalsky, R.L., Corcoran, D.L., Cullen, B.R., and Ohler, U. (2013). MicroRNA target site identification by integrating sequence and binding information. Nat Methods 10, 630–633. 10.1038/nmeth.2489.

Manakov, S.A., Grant, S.G., and Enright, A.J. (2009). Reciprocal regulation of microRNA and mRNA profiles in neuronal development and synapse formation. BMC Genomics 10, 419. 10.1186/1471-2164-10-419.

Martin, M. (2011). Cutadapt removes adapter sequences from high-throughput sequencing reads. 2011 *17*, 3. 10.14806/ej.17.1.200.

McGeary, S.E., Lin, K.S., Shi, C.Y., Pham, T.M., Bisaria, N., Kelley, G.M., and Bartel, D.P. (2019). The biochemical basis of microRNA targeting efficacy. Science 366. 10.1126/science.aav1741.

Miska, E.A., Alvarez-Saavedra, E., Abbott, A.L., Lau, N.C., Hellman, A.B., McGonagle, S.M., Bartel, D.P., Ambros, V.R., and Horvitz, H.R. (2007). Most Caenorhabditis elegans microRNAs are individually not essential for development or viability. PLoS Genet 3, e215. 10.1371/journal.pgen.0030215.

Moore, M.J., Scheel, T.K., Luna, J.M., Park, C.Y., Fak, J.J., Nishiuchi, E., Rice, C.M., and Darnell, R.B. (2015). miRNA-target chimeras reveal miRNA 3’-end pairing as a major determinant of Argonaute target specificity. Nat Commun 6, 8864. 10.1038/ncomms9864.

Okou, D.T., Steinberg, K.M., Middle, C., Cutler, D.J., Albert, T.J., and Zwick, M.E. (2007). Microarray-based genomic selection for high-throughput resequencing. Nat Methods 4, 907–909. 10.1038/nmeth1109.

Patton, R.D., Sanjeev, M., Woodward, L.A., Mabin, J.W., Bundschuh, R., and Singh, G. (2020). Chemical crosslinking enhances RNA immunoprecipitation for efficient identification of binding sites of proteins that photo-crosslink poorly with RNA. RNA 26, 1216–1233. 10.1261/rna.074856.120.

Pawlica, P., Yario, T.A., White, S., Wang, J., Moss, W.N., Hui, P., Vinetz, J.M., and Steitz, J.A. (2021). SARS-CoV-2 expresses a microRNA-like small RNA able to selectively repress host genes. Proc Natl Acad Sci U S A 118. 10.1073/pnas.2116668118.

Peng, Y., and Croce, C.M. (2016). The role of MicroRNAs in human cancer. Signal Transduct Target Ther 1, 15004. 10.1038/sigtrans.2015.4.

Quinlan, S., Kenny, A., Medina, M., Engel, T., and Jimenez-Mateos, E.M. (2017). MicroRNAs in Neurodegenerative Diseases. Int Rev Cell Mol Biol 334, 309–343. 10.1016/bs.ircmb.2017.04.002.

Rupaimoole, R., and Slack, F.J. (2017). MicroRNA therapeutics: towards a new era for the management of cancer and other diseases. Nat Rev Drug Discov 16, 203–222. 10.1038/nrd.2016.246.

Sarkozy, M., Kahan, Z., and Csont, T. (2018). A myriad of roles of miR-25 in health and disease. Oncotarget 9, 21580–21612. 10.18632/oncotarget.24662.

Shen, E.Z., Chen, H., Ozturk, A.R., Tu, S., Shirayama, M., Tang, W., Ding, Y.H., Dai, S.Y., Weng, Z., and Mello, C.C. (2018). Identification of piRNA Binding Sites Reveals the Argonaute Regulatory Landscape of the C. elegans Germline. Cell 172, 937–951 e918. 10.1016/j.cell.2018.02.002.

Siegfried, N.A., Busan, S., Rice, G.M., Nelson, J.A., and Weeks, K.M. (2014). RNA motif discovery by SHAPE and mutational profiling (SHAPE-MaP). Nat Methods 11, 959–965. 10.1038/nmeth.3029.

Smith, T., Heger, A., and Sudbery, I. (2017). UMI-tools: modeling sequencing errors in Unique Molecular Identifiers to improve quantification accuracy. Genome Res 27, 491–499. 10.1101/gr.209601.116.

Sternburg, E.L., Estep, J.A., Nguyen, D.K., Li, Y., and Karginov, F.V. (2018). Antagonistic and cooperative AGO2-PUM interactions in regulating mRNAs. Sci Rep 8, 15316. 10.1038/s41598-018-33596-4.

Tan, F.E., Sathe, S., Wheeler, E.C., Nussbacher, J.K., Peter, S., and Yeo, G.W. (2019). A Transcriptome-wide Translational Program Defined by LIN28B Expression Level. Mol Cell 73, 304–313 e303. 10.1016/j.molcel.2018.10.041.

Tewhey, R., Warner, J.B., Nakano, M., Libby, B., Medkova, M., David, P.H., Kotsopoulos, S.K., Samuels, M.L., Hutchison, J.B., Larson, J.W., et al. (2009). Microdroplet-based PCR enrichment for large-scale targeted sequencing. Nat Biotechnol 27, 1025–1031. 10.1038/nbt.1583.

van Dongen, S., Abreu-Goodger, C., and Enright, A.J. (2008). Detecting microRNA binding and siRNA off-target effects from expression data. Nat Methods 5, 1023–1025. 10.1038/nmeth.1267.

Van Nostrand, E.L., Freese, P., Pratt, G.A., Wang, X., Wei, X., Xiao, R., Blue, S.M., Chen, J.Y., Cody, N.A.L., Dominguez, D., et al. (2020a). A large-scale binding and functional map of human RNA-binding proteins. Nature 583, 711–719. 10.1038/s41586-020-2077-3.

Van Nostrand, E.L., Nguyen, T.B., Gelboin-Burkhart, C., Wang, R., Blue, S.M., Pratt, G.A., Louie, A.L., and Yeo, G.W. (2017a). Robust, Cost-Effective Profiling of RNA Binding Protein Targets with Single-end Enhanced Crosslinking and Immunoprecipitation (seCLIP). Methods Mol Biol 1648, 177–200. 10.1007/978-1-4939-7204-3_14.

Van Nostrand, E.L., Pratt, G.A., Shishkin, A.A., Gelboin-Burkhart, C., Fang, M.Y., Sundararaman, B., Blue, S.M., Nguyen, T.B., Surka, C., Elkins, K., et al. (2016). Robust transcriptome-wide discovery of RNA-binding protein binding sites with enhanced CLIP (eCLIP). Nat Methods 13, 508–514. 10.1038/nmeth.3810.

Van Nostrand, E.L., Pratt, G.A., Yee, B.A., Wheeler, E.C., Blue, S.M., Mueller, J., Park, S.S., Garcia, K.E., Gelboin-Burkhart, C., Nguyen, T.B., et al. (2020b). Principles of RNA processing from analysis of enhanced CLIP maps for 150 RNA binding proteins. Genome Biol 21, 90. 10.1186/s13059-020-01982-9.

Van Nostrand, E.L., Shishkin, A.A., Pratt, G.A., Nguyen, T.B., and Yeo, G.W. (2017b). Variation in single-nucleotide sensitivity of eCLIP derived from reverse transcription conditions. Methods 126, 29–37. 10.1016/j.ymeth.2017.08.002.

Volders, P.J., Anckaert, J., Verheggen, K., Nuytens, J., Martens, L., Mestdagh, P., and Vandesompele, J. (2019). LNCipedia 5: towards a reference set of human long non-coding RNAs. Nucleic Acids Res 47, D135–D139. 10.1093/nar/gky1031.

Ying, W., Riopel, M., Bandyopadhyay, G., Dong, Y., Birmingham, A., Seo, J.B., Ofrecio, J.M., Wollam, J., Hernandez-Carretero, A., Fu, W., et al. (2017). Adipose Tissue Macrophage-Derived Exosomal miRNAs Can Modulate In Vivo and In Vitro Insulin Sensitivity. Cell 171, 372–384 e312. 10.1016/j.cell.2017.08.035.

Zarnegar, B.J., Flynn, R.A., Shen, Y., Do, B.T., Chang, H.Y., and Khavari, P.A. (2016). irCLIP platform for efficient characterization of protein-RNA interactions. Nat Methods 13, 489–492. 10.1038/nmeth.3840.

Zhu, S., Rooney, S., and Michlewski, G. (2020). RNA-Targeted Therapies and High-Throughput Screening Methods. Int J Mol Sci 21. 10.3390/ijms21082996.

