## Supplemental Protocol for "Scalable and deep profiling of mRNA targets for individual microRNAs with chimeric eCLIP"

Eric Van Nostrand / Yeo Lab – contact &

### Chimeric eCLIP Experimental Procedures

---

P = user-prepared personal stock

S = stock from enzyme kit

P **iCLIP Lysis Buffer**
50 mM Tris-HCl pH 7.4
100 mM NaCl
1% NP-40 (Igepal CA630)
0.1% SDS
0.5% sodium deoxycholate (protect from light)

P **High Salt Wash Buffer**
50 mM Tris-HCl pH 7.4
1 M NaCl
1 mM EDTA
1% NP-40
0.1% SDS
0.5% sodium deoxycholate (protect from light)

P **Wash Buffer**
20 mM Tris-HCl pH 7.4
10 mM $MgCl_2$
0.2% Tween-20
5 mM NaCl

P **4x Mn SSIII/IV buffer**
**should be prepared fresh just before use*
200 mM Tris pH 8
300 mM KCl
12 mM $MnCl_2$

P **PKS Buffer**
100mM Tris-HCl pH 7.4
50mM NaCl
10mM EDTA
0.2% SDS

P **TT Elution Buffer**
10 mM Tris pH 7.5
0.01% Tween-20
0.1 mM EDTA

P **80% Ethanol**

P **1% Tween-20**

P **1M HCl**

P **1M NaOH**

P **RLTW Buffer**
1x Qiagen cat # 79216
0.025% Tween-20

P **Bead Elution Buffer**
5 mM Tris pH 7.5
0.01% Tween-20
0.05 mM EDTA

### Buffers & Solutions

#### Enzymes

| | | | |
|---|---|---|---|
| **Turbo DNase** | 2 U/µl | LifeTech | AM2239 |
| **RNase I** | 100 U/µl | LifeTech | AM2295 |
| **FastAP** | 1 U/µl | LifeTech | EF0652 |
| **Murine RNase Inhibitor** | 40 U/µl | NEB | M0314L |
| **T4 PNK** | 10 U/µl | NEB | M0201L |
| **T4 Polynucleotide Kinase (3' phosphatase minus)** | | NEB | M0236S |
| **T4 RNA ligase 1 high conc** | 30 U/µl | NEB | M0437M |
| **Proteinase K** | 0.8 U/µl | NEB | P8107S |
| **Q5 PCR Master Mix** | | NEB | M0492L |
| **Protease Inhibitor Cocktail III** | | EMD Millipore | 539134-1SET |
| **SuperScript IV Reverse Transcriptase** | | LifeTech | 18090010 |
| **Exo-SAP-IT** | | Affymetrix | 78201 |

#### Beads

| | | | |
|---|---|---|---|
| Dynabeads M-280 sheep anti-mouse | 10 mg/ml | LifeTech | 11201D |
| Dynabeads MyOne Silane | 40 mg/ml | LifeTech | 37002D |
| Agencourt AMPure XP beads | | Beckman Coulter | A63881 |

#### Antibodies

| | | | |
|---|---|---|---|
| Ago2 – IP | 200 µg/ml | Santa Cruz | sc-53521 |
| Ago2 – Western | 1:2000 or 1:4000 | Sino Biological | 50683-R036 |

### Primers

#### RNA oligos:

##### Original RNA adapters:

InvRiL19: /5Phos/rArGrArUrCrGrGrArArGrArGrCrArCrArCrGrUrC/iSpC3/3Bio/

CLASHRiL19bio /5Phos/rArGrArUrCrGrGrArArGrArGrCrArCrArCrGrUrC/iSpC3//3Bio/

(other alternatives:

N9RiT22 /5Phos/rNrNrNrNrNrNrNrNrNrArGrArUrCrGrGrArArGrArGrCrArCrArCrGrUrCrUrG/3SpC3/

CLASHn10RiL19bio /5Phos/rNrNrNrNrNrNrNrNrNrNrArGrArUrCrGrGrArArGrArGrCrArCrArCrGrUrC/iSpC3//3Bio/ )

**(Order 100 nmole RNA oligo, standard desalting; storage stock 200 µM; final concentration 1 µM (input), 4 µM (CLIP)).**

#### DNA oligos:

InvRand3Tr3: /5Phos/NNNNNNNNNNAGATCGGAAGAGCGTCGTGT/3SpC3/

**(Order 100 nmole DNA oligo, standard desalting; storage stock 200 µM; working stock 80 µM; final concentration 3 µM).**

InvAR17: CAGACGTGTGCTCTTCCGA **(25 nmole DNA oligo, standard desalting; storage stock 200 µM; working stock 10 µM; final concentration 0.5 µM).**

**(Below we order page-purified)**

| | |
|---|---|
| PCR_F_D501 | AATGATACGGCGACCACCGAGATCTACACTATAGCCTACACTCTTTCCCTACACGACGCTCTTCCGATCT |
| PCR_F_D502 | AATGATACGGCGACCACCGAGATCTACACATAGAGGCACACTCTTTCCCTACACGACGCTCTTCCGATCT |
| PCR_R_D701 | CAAGCAGAAGACGGCATACGAGATCGAGTAATGTGACTGGAGTTCAGACGTGTGCTCTTCCGATC |
| PCR_R_D702 | CAAGCAGAAGACGGCATACGAGATTCTCCGGAGTGACTGGAGTTCAGACGTGTGCTCTTCCGATC |

**(See Illumina customer service letter for D503-508, D703-712; any standard Illumina HT RNA-seq primers work fine)**

**(stock 100 uM; working 20 uM)**

### Notes

- For this protocol one 'experiment' is typically defined as 4 libraries: 2 eCLIP experiments on UV crosslinked biological replicate samples and 2 size-matched input control (taken from each of the two UV crosslinked samples).
    - For other experiments, one can modify this design to add the paired size-matched input control from the other replicate, add a non-UV crosslinked sample, or add additional library controls (IgG, FLAG or V5 pulldown on wild-type cells, etc.) if desired.
- Although we do not standardly do P32 labeling, we still use the "HOT" and "COLD" membranes nomenclature from iCLIP & other CLIP protocols as a shortcut. COLD = an analytical gel run on 10% of sample run as a standard Western blot; HOT = a preparative gel run on 80% of sample run for membrane cutting & RNA isolation
- When working with Protein G or other antibody-attached beads, be sure beads are never heated or dried (make sure master mixes are ready prior to removing supernatant from final washes)
- Bead washes should be done in the following order:
    1. Remove sample from magnet and buffer to beads
    2. Close tubes well and invert or flick tubes (do not vortex) to mix beads with buffer until solution is homogenous
    3. Transfer tubes to magnet and magnetically separate for 1-2 minutes. During this time it is recommended to slowly invert magnet with tubes a few times (this helps to collect any beads from tube caps).
    4. When the solution is transparent, remove most solution while ensuring the pipette tip does not touch the beads. Next, close the tubes and spin briefly for 1-3 seconds (desktop minicentrifuge), and place back on magnet. The remaining solution can then be removed using 200 µL (or smaller) pipette tips.
- Enzymes should be kept at -20C, preferentially in chilled enzyme coolers, keep them in cooler the whole time.
- We recommend the use of low-retention tips and tubes to ensure minimal sample loss
- Ensure proper hygiene for working with RNA samples (including separation from bacteria or other RNase-containing samples) as well as high-throughput sequencing libraries (we recommend physical separation of work performed on pre-amplified and post-PCR amplified material).

### DAY 0

##### Prepare iCLIP lysis mix

- Pre-chill iCLIP Lysis Buffer
- Per sample (3 million cells): add **5.5 µl 200x Protease Inhibitor Cocktail III** and **11 µl Murine RNase Inhibitor** to **500 uL iCLIP Lysis Buffer**, mix
    - ** Note: RNase Inhibitor may need to be further increased for particularly difficult samples (e.g. Pancreas).

##### Couple antibody to magnetic beads (do this first)

- **Beads and antibodies:**
    - Use **200 µl sheep anti-mouse Dynabeads** per sample
    - Use **5 µg AGO2 (eIF2C12 4F9) antibody (sc-53521, Santa Cruz)** per sample
- **Prepare beads:**
    - Magnetically separate beads, remove supernatant
    - Wash beads 2x in 500 µl cold **iCLIP Lysis Buffer**
    - Resuspend beads in cold **iCLIP Lysis Buffer,** 100 µl per sample
- **Bind antibody:**
    - Add antibody (5 µg per sample) to washed beads
    - Rotate at room temp, 45 min

##### Lyse cells (while ab and beads are binding)

- **Lyse cells:**
    - Retrieve cell pellets from -80°C freezer, immediately add 500 mL cold **iCLIP Lysis Buffer + Protease Inhibitor + RNase Inhibitor Mix** to each pellet, pipette to resuspend

      **2 Pellets per experiment:**
        - Sample 1: IP-A (UV-crosslinked batch #1)
        - Sample 2: IP-B (UV-crosslinked batch #2)
    - Lyse 5 mins on ice

##### RNase treat lysate (while ab+bead binding):

- Sonicate in Bioruptor on ‘low’ setting, 4°C, 5 min, 30sec on / 30 sec off
- Dilute RNase I in PBS** on ice; use 10 µl diluted RNase I per sample
    - ** This likely needs to be optimized for each cell type
    - 3 million 293Ts should start with 1:300
    - 20 million 293Ts we use 1:100
- To lysed sample(s), add 5 µl **Turbo Dnase** (mix immediately before use)
- To lysed sample(s), add 10 µl **diluted RNase I**, mix & immediately proceed to next step
- Incubate in Thermomixer at 1200 rpm, 37°C, 5 mins (exactly), place on ice
- Centrifuge 15,000g, 4°C, 3min
- Transfer cleared lysate to a new tube

##### Capture RBP-RNA complexes on beads

- Wash antibody beads 2x in 500 µl cold **iCLIP Lysis Buffer**
- Add cleared lysates to washed antibody beads
- Rotate 4°C, 2 h or overnight (in cold room)

### DAY 1

#### Step: SAVE INPUT SAMPLES

- Mix samples well
- To new tube, take 20 µL (2%) of Sample 1 (A-Input), 2 (B-Input) for 'HOT' (RNA) gel, store at 4°C
- To new tube, take 20 uL (2%) of Sample 1 (A-Input), 2 (B-Input) for 'COLD' (western) gel; store at 4°C

##### Wash beads

- Wash 3x with 500 µL cold **High Salt Wash Buffer**
- Wash 3x with 500 µL cold **Wash Buffer**

##### T4PNK Minus rxn (on-bead)

- Prepare **T4PNKminus mix** on ice; 100 µl per sample:
  - $H_2O$ 83.6 µl
  - 10x PNK pH 7 Buffer 10 µl
  - 0.1M ATP 1 µl
  - 5M NaCl 0.4 µl
  - Murine RNase Inhibitor 1 µl
  - T4 PNK Minus (M0236S) enzyme 3 µl
- Mix, add **100 µl** to each sample, incubate in Thermomixer at 1200 rpm, 37°C, 20 min

##### Wash beads

- Wash 1x with 500 µL cold **High Salt Wash Buffer**
- Wash 3x with 500 µL cold **Wash Buffer**

##### RNA chimeric ligation (on-bead)

- Prepare **RNA ligase mix** on ice; 180 µl per sample:
  - H2O 80.4 µl
  - 10x RNA Ligase Buffer 18 µl
  - 1% Tween-20 3.6 µl
  - DMSO 5.4 µl
  - 100 mM ATP 1.8 µl
  - 50% PEG-8000 54 µl
  - Murine RNase Inhibitor 2.4 µl
  - T4 RNA Ligase HC enzyme 14.4 µl
- Mix, add **180 µl** to each sample, incubate room temp in rotator for 1-2 hr.

##### Wash beads

- Wash 3x with 500 µL cold **High Salt Wash Buffer**
- Wash 3x with 500 µL cold **Wash Buffer**

##### FastAP treat beads (all samples except IgG)

- Prepare **FastAP reaction mix** on ice; 50 µl per sample:
  - $H_2O$ 38 µl
  - 10x FastAP Buffer 5 µl
  - Murine RNase Inhibitor 2 µl
  - Turbo DNase 2 µl
  - FastAP enzyme 3 µl
- Mix, add **50 µl** to each sample, incubate in Thermomixer at 1200 rpm, 37°C, 10 min

##### PNK treat beads

- While beads are incubating, prepare **PNK master mix** on ice; 150 µl per sample:
  - $H_2O$ 126 µl
  - 10x PNK pH 7 Buffer 20 µl
  - T4 PNK enzyme 4 µl
- Mix, add **150 µl** to each sample (don't remove FastAP mix), incubate at 1200 rpm, 37°C, 20 min

##### Wash beads

- Wash 1x with 500 µl cold **High Salt Wash Buffer**,
- Wash 3x with 500 µL cold **Wash Buffer**
- Prepare the 3' ligation master mix
- Just before adding the 3' ligation master mix, briefly spin tubes in minifuge, magnetically separate, remove residual liquid with fine tip

##### Ligate 3' RNA linker (on-bead)

- Prepare **3' ligation master mix** at room temperature (not on ice); 27 µl per sample:
  - $H_2O$ 10.4 µl
  - 10x RNA Ligase Buffer 3 µl
  - 1% Tween-20 0.6 µl
  - DMSO 0.9 µl
  - 100 mM ATP 0.3 µl
  - 50% PEG-8000 9 µl
  - Murine RNase Inhibitor 0.4 µl
  - RNA Ligase high conc. 2.4 µl
- Mix carefully by pipetting or flicking (do not vortex) and add **25 µl** to each sample
- To each sample, add
  - 0.3 µl TT elution buffer
  - 2.085 µl $H_2O$
  - 0.615 µl CLASHInvRiL19 (200 µM)
- Incubate at room temperature for 75 min; flick to mix every ~10 min or rotate constantly

##### Wash beads (resume IgG sample here)

- Wash 1x with 500 µL cold **Wash Buffer**, magnetically separate, remove supernatant
- Wash 1x with 500 µL cold **High Salt Wash Buffer**
- Wash 2x with 500 µL cold **Wash Buffer**

**Either Direct ProtK (Ago2 human/mouse antibody) or gel below:**

#### Direct ProtK:

- **Resuspend in ProtK solution:**
  + 65 uL PKS buffer
  + 15 uL Proteinase K
  Incubate 37°C for 20 min
  Incubate 50°C for 20 min
- **IF DOING PROBE-CAPTURE, SAVE SAMPLE HERE (see Appendix A)**

#### Zymo Column Cleanup

- Add 40 uL H2O (bring sample up to 120 uL total)
- Add 240 uL RNA binding buffer & pipette mix or shake well to mix
- Add 360 uL 100% EtOH and pipette mix well (at least 6 times)
- Add mixture to Zymo-Spin column
- Centrifuge 30 sec on benchtop minifuge or 5,000g
- Repeat column binding for sample: carefully pipette flow-through back onto column and centrifuge again, discard flow-through
- Add 400 uL RNA Wash Buffer, centrifuge for 30 sec, discard flow through
- Make DNase MM (gentle when mixing)
  - 17.5 uL Zymo DNA Digestion buffer
  - 2.5 uL Zymo DNase I
- Add 20 uL DNase MM directly to column
- Incubate at room temp for 15 mins
- Add 400 uL RNA Prep Buffer, centrifuge for 30 sec, discard flow through
- Add 480 uL RNA Wash buffer, centrifuge for 30 sec, discard flow through
- Repeat wash (add 480 uL RNA Wash buffer, centrifuge for 30 sec, discard flow through)
- Add 200 uL RNA Wash buffer, centrifuge 1 min 9000g in desktop centrifuge, discard flow through
- Centrifuge 2 additional minutes
- Transfer column to new 1.5 mL tube (avoid getting Wash Buffer on column)
- Let sample air dry for 2 min
- **Elute: Add 10 µl H2O to column, let sit for 1 min, centrifuge for 30 sec at 9,000g**
- **Repeat elution in same eluate: take the flow-through and pipette it onto the column again, sit for 1 minute, and centrifigue 30 sec at 9,000g**
- **Place IP samples in -80 C until reverse transcription**

**Potential -80 C stopping point**

**OR**

#### Prepare samples for gel loading

- **IP-Bead samples (HOT and COLD)**:
  **** Note: HOT & COLD are named relative to iCLIP gels; neither is radioactive in eCLIP**
  **HOT = CLIP gel** – for membrane transfer & RNA isolation
  **COLD = WESTERN gel** – for western imaging
  - **Remove s/n,** add **100** µl cold **Wash Buffer**, resuspend beads well
  - Move 20 µl to new tube #1 = **COLD IP Bead samples**
  - For **HOT IP Bead samples,** remove remaining 80 µl Wash Buffer and add 20 µl Wash Buffer

**Prepare input and IP samples for SDS-PAGE corresponding to following table:**

| Final Sample Composition for Gel Loading | | | | | | |
|---|---|---|---|---|---|---|
| **Buffer** | **IgG Bead *(if applicable)*** | **Cold 0.1% Input *(if applicable)*** | **Cold Inputs** | **Cold IP Beads** | **HOT Inputs** | **HOT IP Beads** |
| Wash Buffer | 100 µl | 20 µl *(add 18 µl to 2 µl)* | 20 µl | 20 µl | 20 µl | 20 µl |
| 4x NuPAGE LDS Buffer | 37.5 µl | 7.5 µl | 7.5 µl | 7.5 µl | 7.5 µl | 7.5 µl |
| 1M DTT | 15 µl | 3 µl | 3 µl | 3 µl | 3 µl | 3 µl |

- Denature all samples in Thermomixer, 1200 rpm, 70°C, 10 min
- Cool on ice 1 min, spin briefly in minifuge
- For **all samples**, magnetically separate prior to loading (IP AND Inputs have beads)

**Potential (not recommended) -20 C stopping point**

**Load and run gels** **your loading scheme may vary depending on conditions**

- **Load HOT (preparative gel for RNA transfer and isolation) (4-12% Bis-Tris, 10-well, 1.5 mm)**

| **HOT GEL** | 1 | 2 | 3 | 4 | 5 | 6 | 7 | 8 | 9 | 10 |
|---|---|---|---|---|---|---|---|---|---|---|
| Sample | M | **Input A** | m | **Input B** | m | **A-IP** | m | **B-IP** | M | m |
| Volume to Load | 5 µl | 30 µl | 1 µl | 30 µl | 1 µl | 30 µl | 1 µl | 30 µl | 5 µl | 1 µl |
| % of Sample Represented | | 2% | | 2% | | 80% | | 80% | | |

- **Load COLD (Western blot imaging gel) (4-12% Bis-Tris, 10-well, 1.5 mm)**

| **COLD GEL** | 1 | 2 | 3 | 4 | 5 | 6 | 7 | 8 | 9 | 10 |
|---|---|---|---|---|---|---|---|---|---|---|
| Sample | M | **A-IP** | **A-INPUT** | **B-IP** | **B-INPUT** | **M** | **-** | **-** | **-** | **-** |
| Volume to Load | 5 µl | 15 µl | 15 µl | 15 µl | 15 µl | 5 µl | | | | |
| % of Sample Represented | | 10% | 1% | 10% | 1% | | | | | |

- Save remaining 15 µl of COLD samples at -20 C for backup
- Run at 150V in 1X MOPS Running Buffer, 75 min or until dye front is at the bottom

**Transfer to membranes**

- **COLD imaging gel: iBlot transfer (see Quick Reference guide for pictures and more details)**
    - iBlot2 ‘mini’ stacks are for 1 gel, ‘regular’ stacks are for 2 gels
    - Unseal the Transfer Stack
    - Take the top stock off and place it to the side
    - Place bottom stack (in plastic tray) into the iBlot2 machine
    - Crack open the NuPAGE gel cassette and remove the comb and bottom ‘bump’ areas

- Carefully wet the gel in DI water and place it on top of the bottom iBlot stack
- Soak an iBlot Filter Paper in DI water, and place it on top of the bottom iBlot stack
- Remove air bubbles using the roller
- Remove the white plastic separator from the top stack and place the top stack over the filter paper
- Remove air bubbles using the roller
- Place an iBlot Absorbent Pad on top of the top stack (make sure the electrical contacts are aligned properly)
- Close the lid of the iBlot
- Run program (P0 is our typical program)
- After completion, check that ladder has properly transferred to membrane
- Place membrane in western membrane case with 10 mL of appropriate blocking buffer (either Azure Chemi or Fluorescent blocking buffer)
- Incubate on nutator at room temp, 30 min
- Probe with primary antibody: Replace blocking buffer with fresh buffer, and add 1:4000 µg/ml **primary AGO2 antibody (50683-R036, Sino Biological)**
- Incubate on nutator either at room temp for 1 hr or overnight at 4ºC

- **HOT library preparation gel:**
  - Have pre-prepared COLD (4 deg) transfer buffer with methanol: 1x NuPAGE transfer buffer, 10% methanol
  - Prepare Nitrocellulose membrane(s): incubate in transfer buffer for > 1 min
  - Wet sponges and Whatman papers in transfer buffer with methanol
  - Assemble transfer stacks, from bottom to top (black side of stack holder on bottom):

    1x sponge – 2x Whatman paper – gel – membrane – 2x Whatman paper – 1x sponge

  HOT gel: Nitrocellulose membrane from iBlot stack

- **Transfer:**
  - overnight 30V (preferred) <u>OR</u>
  - 2 hr 200 mA (if doing this, only hook up one transfer box per power supply)

### Day 2

- Remove HOT membrane, rinse quickly once with sterile 1X PBS, wrap in Saran wrap, store at -20C

##### Develop COLD membrane

- Block in Azure Chemi Blot Blocking Buffer (AC2148), room temp, 30 min
- Probe with primary antibody: 0.2-0.5 µg/ml in Azure Chemi Blot Blocking Buffer, room temp, 1 hr.
- Wash 3x with TBST, 5 min each
- Probe with secondary antibody: 1:4000 Mouse TrueBlot HRP in Azure Chemi Blot Blocking Buffer, room temp, 1 – 3 h
    - (Note: if western fails or signal is low, 1:1000 gives higher signal)
- Wash 3x with TBST, 5 min each
- Mix equal volumes of ECL Reagent 1 + Reagent 2 (or 40:1 of ECL Plus Substrate A to Substrate B), add to membrane and incubate (mix/rotate) for 1-5 min. (1ml final volume per membrane)
- Develop 30 sec & 5 min, then judge signal (15 min maximum; if 15 sec is still too bright, expose two films

##### Cut HOT membrane

- Note RBP band on film with respect to protein markers
- Place HOT membrane on clean glass/metal surface
- Using a fresh razor blade, cut lane from HOT membrane from 95-225 kDa
    - This starts at AGO2 (~85 kDa) + ~40nt RNA (~12 kDa) and includes chimeras up to ~400nt
- Slice membrane pieces into ~1-2 mm slices, use a fresh razor blade for each sample
- Transfer slices to Eppendorf tube and centrifuge – place tube on ice if doing many samples

##### Release RNA from membrane

- Prepare **Proteinase K SDS mix** on ice, 150 µl per sample:
    - PKS Buffer 130 µl
    - Proteinase K 20 µl
- Mix, add **150 µl** Proteinase K SDS mix to membrane slices, incubate in Thermomixer at 1200 rpm, 37 C, 20 min (make sure all membrane slices are submerged)
- Further incubate in Thermomixer at 1200 rpm, 50 C, for an additional 20 min
- Transfer all solution to a fresh 1.5 mL DNA loBind tube
- Rinse membrane with 55 µl of water, and add to supernatant above (giving 200 µL total)

##### Zymo column cleanup – RNA Clean & Concentrator-5 columns (Cat R1016)

- Add 400 µl (2x volumes) RNA binding buffer, pipette mix well
- Add 700 µl (3.5x starting volume) of 100% ethanol & pipette mix well (take care to avoid spilling of sample)
- Transfer 650 µl of mixed sample to Zymo-Spin column
- Centrifuge 30 sec on benchtop minifuge or 5,000g
- Repeat column binding for sample: carefully pipette flow-through back onto column and centrifuge again, discard flow-through
- Repeat by reloading an additional 650 µl volume until all sample has been spun through column
- Add 400 µl RNA Prep Buffer, centrifuge for 30 sec, discard flow through
- Add 500 µl RNA Wash Buffer, centrifuge for 30 sec, discard flow through
- Add 500 µl RNA Wash Buffer, centrifuge for 30 sec, discard flow through
- Add 200 µl RNA Wash Buffer, centrifuge for 1 minute at 9,000g, discard flow through
- Centrifuge additional 2 mins
- Transfer column to new 1.5 mL tube (avoid getting Wash Buffer on column)
- **Elute: Add 10 µl H2O to column, let sit for 1 min, centrifuge for 30 sec at 9,000g**
- **Repeat elution in same eluate: take the flow-through and pipette it onto the column again, sit for 1 minute, and centrifuge 30 sec at 9,000g**
- **Place IP samples in -80 C until reverse transcription**

**Potential -80 C stopping point**

### Day 3

**START Inputs only →**

#### FastAP treat input RNA

- Prepare **FastAP master mix**; 11 µl per sample:
  - 10X FastAP Buffer 2 µl
  - $H_2O$ 6 µl
  - Murine RNase Inhibitor 1 µl
  - FastAP enzyme 2 µl

- Mix, add **11 µl** to samples, mix, incubate in Thermomixer at 1200 rpm, 37 C, 10 min

#### PNK treat input RNA

- Prepare **PNK master mix**; 75 µl per sample:
  - $H_2O$ 61 µl
  - 10X PNK Buffer 9 µl
  - Turbo DNase 1 µl
  - PNK enzyme 4 µl

- Mix, add **75 µl** to samples, mix, incubate in Thermomixer at 1200 rpm, 37 C, 20 min

#### Zymo column cleanup – RNA Clean & Concentrator-5 columns (Cat R1016)

- Add 200 µl (2x volumes) RNA binding buffer directly to 95 µL repaired RNA sample, pipette mix well
- Add 300 µl (3x starting volume) of 100% ethanol & pipette mix well (avoid spilling of sample)
- Transfer all sample to Zymo-Spin column, centrifuge 30 sec at 5,000g
- Repeat column binding for sample: carefully pipette flow-through back onto column and centrifuge again, discard flow-through
- Add 400 µl RNA Prep Buffer, centrifuge for 30 sec, discard flow through
- Add 500 µl RNA Wash Buffer, centrifuge for 30 sec, discard flow through
- Add 500 µl RNA Wash Buffer, centrifuge for 30 sec, discard flow through
- Add 200 µl RNA Wash Buffer, centrifuge for 1 minute at 9,000g, discard flow through
- Centrifuge additional 2 mins
- Transfer column to new 1.5 mL tube (avoid getting Wash Buffer on column)
- **Elute: Add 10 µl H2O to column, let sit for 1 min, centrifuge for 30 sec at 9,000g**
- **Repeat elution in same eluate: take the flow-through and pipette it onto the column again, sit for 1 minute, and centrifuge 30 sec at 9,000g**

  **Potential -80 C stopping point**

##### 3’ linker ligate input RNA

- **Anneal adapter:**
    - Take 5 µl of RNA (from above) – remainder of input is kept at -80 C as backup
    - To each sample, add:
        - 0.2 µl TT elution buffer
        - 0.8 µl $H_2O$
        - 0.8 µl DMSO
        - 0.2 µl CLASHInvRiL19 (200 µM)
    - Incubate 65ºC, 2 min → place on ice >1 min

- **Prepare ligation master mix; 13.5 µl per sample** at room temperature (not on ice)**:**

| | |
|---|---|
| $H_2O$ | 2.8 µl |
| 50% PEG 8000 | 6 µl |
| 10X RNA Ligase Buffer | 2 µl |
| 1% Tween20 | 0.4 µl |
| DMSO | 0.6 µl |
| 100 mM ATP | 0.2 µl |
| Murine RNase Inhibitor | 0.3 µl |
| RNA Ligase High Conc | 1.2 µl |

- Flick/pipette mix, add **13.5 µl** to each sample, flick/pipette-mix, incubate at room temp for 60 min
- Flick to mix every ~15 min

##### Silane cleanup input RNA

- **Prepare beads:**
    - To 10 µl **MyONE Silane Beads** per sample, add 5x volume RLT (e.g. for 4 samples, use 40 µl of beads and add 200 µl of RLT)
    - Pipette mix, magnetically separate, and remove supernatant
    - Resuspend beads in 63 µl RLTW buffer (RLT + 0.025% Tween-20) **per sample (**e.g. for 4 samples, use 250 µl RLTW buffer). Mix well.

- **Bind RNA:**
    - Add 61 µl of bead/RLTW mixture above to each RNA sample, mix
    - Add 65 µl **100% EtOH** to each sample
    - Pipette mix 10 times, leave pipette tip in tube, pipette mix every ~3-5 min for 10 min

- **Wash beads:**
    - Magnetically separate and discard supernatant
    - Add 300 µl (PCR strip tubes) or 1 mL (1.5 mL tubes) **80% EtOH**, pipette resuspend
    - After 30 s, magnetically separate, remove supernatant
    - Repeat wash with 300 µl (PCR strip tubes) or 1 mL (1.5 mL tubes) **80% EtOH**
    - After 30 s, magnetically separate, remove supernatant
    - Wash 3$^{rd}$ time with 100 µl (PCR strip tubes) or 750 µl (1.5 mL tubes) **80% EtOH.**
    - Spin briefly in picoFuge, magnetically separate, remove residual liquid with fine tip
    - Dry beads well, i.e. until they stop “shining - no ethanol should be left on the bottom of strip. Beads are over-dry when they change color from brown to orange/rusty color.

- **Elute RNA:**
    - Resuspend in **9.5 µl Bead Elution Buffer**, let sit for 5 min
    - Magnetically separate, transfer supernatants to strip tube(s) (will be ~**9 µl**)

**Potential -80 C stopping point**

---

#### ←END Inputs only

---

All CLIP and INPUT samples are now synchronized.

#### Reverse transcribe RNA (ALL CLIP and INPUTS)

- **Anneal primer** in 8-well strip tubes:
  - To ~9µl of RNA, add:
    - 1 uL dNTP mix (10 mM each)
    - 0.5 uL RT primer InvAR17 (10 uM)
  - Heat 65°C for 2 min in pre-heated PCR block, place immediately on ice (do not cool down in PCR block)

- **Prepare RT master mix** on ice; 10 µl per sample:
  - 4x Mn SSIII/IV Buffer 5 µl
  - H2O 2.8 µl
  - 0.1M DTT 1 µl
  - Murine RNase Inhibitor 0.4 µl
  - Superscript IV Enzyme 0.8 µl

- Add 10 µl to each sample, mix, incubate 55 C, 20 min in pre-heated PCR block

#### Cleanup cDNA

- **ExoSAP Treatment**
  - Add **2.5µl ExoSAP-IT** to each sample, vortex, spin down
  - Incubate 37°C for 15 mins on PCR block
  - Add 1 µl **0.5M EDTA**, pipette-mix

- **RNA removal**
  - Add 3 µl of **1M NaOH**, pipette-mix
  - Incubate 70°C, 10 min on PCR block
  - Add 3 µl of **1M HCl**, pipette-mix (to fix pH)

#### Silane cleanup cDNA

- **Prepare beads:**
  - Take **5µl MyONE Silane beads** per sample and add 5x volume RLT buffer, mix well
  - Magnetically separate and remove supernatant
  - Resuspend beads in 93 µL **RLTW buffer per sample**

- **Bind cDNA:**
  - Add **90 µl beads+RLTW** to each sample
  - Add **108 µl 100% EtOH**
  - Pipette mix (10+ times), leave pipette tip in tube, pipette mix twice (every 5min) **for total incubation of 10 minutes at room temp**

- **Wash beads:**
  - Magnetically separate, remove supernatant
  - Wash 2× with 300 µl 80% EtOH (add 80% ethanol, move back and forth on magnet, magnetically separate, remove supernatant)
  - Wash 1x with 150 µl 80% EtOH (spin briefly in picoFuge, magnetically separate, remove residual liquid with fine tip)
  - Air-dry 5 min

##### 5' linker ligate cDNA (on-bead, in 10ul)

- **Add cDNA adapter mix**
  - To each sample, add:
    - 1.45 µl of TT Elution Buffer
    - 0.25 µl **Invrand10_3Tr3** adapter (200 uM)
    - 0.8 µl 100% **DMSO**
  - Heat at 70°C, 2 min, place immediately on ice for >1 min

- **Prepare ligation master mix** on bench:
  - $H_2O$ 1.4 µl
  - 10x NEB RNA Ligase Buffer (with DTT) 1 µl
  - 0.1M DTT 0.2 µl
  - 0.1M ATP 0.1 µl
  - 1% Tween-20 0.2 µl
  - 50% PEG 8000 3.6 µl
  - RNA Ligase high conc (M0437M) 1 µl
  - 5' Deadenylase (M0331S) 0.3 µl

- Flick to mix twice, spin down briefly, add **7.8 µl** to each sample: stir sample with pipette tip, then add master mix slowly with stirring; needs to be homogeneous
- Incubate at room temp overnight on rotator

#### Day 4

##### Silane cleanup linker-ligated cDNA

- To each sample **add 5 µl of Bead Elution Buffer**, making 15 µl total.

- **Prepare beads:**
  - Take **2.5 µl MyONE Silane beads** per sample, add 5x volume RLT
  - Magnetically separate and remove supernatant
  - Resuspend beads in 47 µL RLTW buffer per sample

- **Bind RNA:**
  - Add **45 µl** beads+**RLTW buffer mix** to each sample
  - Add **45 µl 100% EtOH**
  - Pipette mix, leave pipette tip in tube, pipette mix twice, for 10 min total

- **Wash beads:**
  - Magnetically separate, remove supernatant
  - Wash 2× with 300 µl 80% EtOH (add 80% ethanol, move back and forth on magnet, magnetically separate, remove supernatant)
  - Wash 1x with 150 µl 80% EtOH (spin briefly in picoFuge, magnetically separate, remove residual liquid with fine tip)
  - Air-dry 5 min

- **Elute ligated cDNA:**
  - Resuspend in 25 µl **Bead Elution Buffer**, let sit for 5 min
    - If performing gene-specific enrichment, elute ligated cDNA in **40 µl** of **Bead Elution Buffer & see Appendix B**
  - Magnetically separate, transfer **25 µl** sample to new tube

**Potential -80 C stopping point**

#### qPCR quantify cDNA

- Prepare **qPCR master mix**; 19 µl per sample:
  - Luna 2x qPCR master mix 10.0 µl
  - $H_2O$ 8.2 µl
  - qPCR primer mix 0.8 µl (10 uM each qPCR-grade D5x/D7x mix)
- Mix, dispense into 96-well qPCR plate, add **1 µl 1:10 diluted (in $H_2O$) cDNA**, seal, mix
- **qPCR conditions (preset protocol: 30 cycle, Luna qPCR, no melt)**
  - 95 C for 1 min
  - 95 C for 15 sec
  - 60 C for 30 sec -> take image

  30x

**Below is the conversion for Cielo6 qPCR Cq values to final PCR cycle values; on BioRad C384 machines, it was Ct minus 3 or 4 cycles**

**** Note: we use the automatically calculated Ct for this; these cycle numbers may change based on your lab setup, so for the first couple CLIPs it is best to err on the side of 1 or 2 extra PCR cycles. If final libraries are > 50 nM (especially if > 100 nM), you should decrease a couple cycles and redo PCR from remaining cDNA.**

#### PCR amplify cDNA

- Typical: Input 9 total cycles (6 + 3), CLIP 16 (6 + 10) total cycles
  - Note that 18 cycles will yield ~30-50% PCR duplicated libraries (further increasing above 18 cycles), which can be ok for RBPs with few specific targets but will be challenging for broad binders.
  - Cycle # for final PCR:

| Cq | uL cDNA | PCR cycles |
|---|---|---|
| 4 | 2.5 ul of 1:10 | 8 |
| 5 | 5 ul of 1:10 | 8 |
| 6 | 1 | 8 |
| 7 | 3 | 8 |
| 8 | 6 | 8 |
| 9 | 12 | 8 |
| 10 | 12 | 9 |
| 11 | 12 | 10 |
| 12 | 12 | 11 |
| 13 | 12 | 12 |
| 14 | 12 | 13 |
| 15 | 12 | 14 |

#### PCR amplify cDNA

- Typical: Input 9 total cycles (6 + 3), CLIP 16 (6 + 10) total cycles
  - Note that 18 cycles will yield ~30-50% PCR duplicated libraries (further increasing above 18 cycles), which can be ok for RBPs with few specific targets but will be challenging for broad binders.
  - Cycle # for final PCR: 3 cycles less than the qPCR Ct of the 1:10 diluted sample

- Prepare **PCR** on ice; 40 µl total per sample:

| | | |
|---|---|---|
| ○ Ligated cDNA | 12 µl | (save remaining 12 µl at -80 C as backup) |
| ○ $H_2O$ | 4 µl | |
| ○ 20 µM right primer (D50x) | 2 µl | |
| ○ 20 µM left primer (D70x) | 2 µl | |
| ○ 2x Q5 PCR master mix | 20 µl | |

- PCR conditions (cycle # depending on library):
  - 98ºC for 30 s
  - 98ºC for 15 sec -> 68ºC for 30 sec -> 72ºC for 40 sec (x6 cycles)
  - 98ºC for 15 sec -> 72ºC for 60 sec (x ? cycles)
  - 72ºC 1 min
  - 4ºC hold

#### SPRI cleanup library

- Resuspend **AmpureXP beads** well by vortexing
  a. (Note: beads should be incubated at room temp for 15 min prior to use)
- Add 72 µl bead suspension (do <u>not</u> separate) per 40 µl PCR reaction and pipette mix well
- Incubate at room temp for 10 min (pipette mix 2-3x during incubation)
- Magnetically separate, wash beads 3x with **80% EtOH**
- Dry beads for 5 min on magnet (do not over-dry, i.e. the pellet will 'crack')
- Resuspend in **20 µl of PCR Elution Buffer** and incubate for 5 minutes at room temperature
- Magnetically separate and transfer supernatant to new tubes

#### Gel-purify library

- **Prepare samples and run gel:**
  - Open E-gel EX 2% agarose gel, remove comb and load into E-gel system
  - Load 10 uL $H_2O$ or E-gel sample buffer into M (marker) lane – do not use this for ladder or sample (it tends to smear and run strangely)
  - Load 6 uL 50bp E-Gel ladder into lanes 1 and 10
  - Add 2 uL E-gel sample buffer to each sample and load samples into lanes 2-9
  - Run E-gel for 10-12 mins
  - Take picture using E-gel camera
- **Gel-extract library from gel:**
  - Crack open E-gel using spatula (take care to avoid tearing gel; if possible, keep gel on back faceplate so that the samples are in the same orientation as they were loaded)
  - Under blue light illumination, cut gel slices **195**-350 bp and place into 15 mL conical tubes, using fresh razor blades for each sample; keep cross-contamination to minimum
    - Keep in mind: adapter-dimer (including RNA adapter) is **156** bp, so chimeras (20nt miRNA + 20nt target) require at least 40nt of fragment

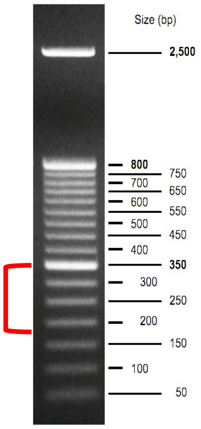


- **Cut & elute gel** using Qiagen MinElute gel extraction kit:
  - Weigh 15 mL conical with gel slice (blank with empty conical tube)
  - Calculate gel weight, add 6x volumes of **Buffer QG** to melt gel (e.g. for 100 mg gel, add 600 µl QG)
  - Melt gel at room temp (do <u>not</u> heat) on benchtop (can shake to help melt, but don't vortex)
  - After gel is melted, add 1x volume of original gel of **isopropanol** & <u>mix well</u> (100 mg gel = 100 µl isopropanol)
  - Load on column (750 µl per spin, can do multiple spins, all spins max speed 1 min)
    - **NOTE:** if gel weight is >400 mg, wash 1x with 500 µl Buffer QG after every 4 spins)
  - After all sample has been spun through, wash 1x with 500 µl **Buffer QG**
  - Add 1X with 750 µl **Buffer PE**, spin 1 min, pour out flow-through, spin again 2 min max speed

- Carefully move column to new 1.5 mL tube (avoid any carryover of PE – if any liquid is visible on the outside of the column redo 2 min max speed spin)
- Using a fine tip, pipette all remaining PE buffer from plastic purple rim of the MinElute column
- Air dry 2 mins
- Carefully add 12.5 µl **Buffer EB** directly to the center of the column, incubate 2 min room temp, spin max speed
- For improved yield – repeat the elution (take the flow-through and add it to the column again)

#### Quantitate library (D1000 DNA tapestation)

- 3 µl D1000 loading buffer, 1 µl sample
- Vortex to mix, spin down in microfuge
- Correctly quantify by adding a region to each sample and dragging boundaries to include the <u>entire</u> library peak (usually ~150-600bp).

**Example of a good trace:**

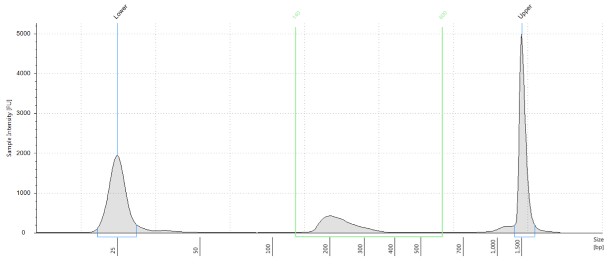


**Region Table**

| From [bp] | To [bp] | Average Size [bp] | Conc. [ng/µl] | Region Molarity [nmol/l] | % of Total | Region Comment | Color |
|---|---|---|---|---|---|---|---|
| 140 | 600 | 233 | 2.79 | 19.2 | 65.55 | | |

### Appendix A

### Probe capture for chimeric-eCLIP

**Required reagents** (beyond standard chimeric eCLIP):

| | |
|---|---|
| 0.5M TCEP | Sigma 646547 |
| 8M LiCl | Sigma L7026 |
| Dynabeads MyOne Streptavidin C1 Beads | Invitrogen 65001 |

**Buffers**

| **3x Hybridization buffer:** | **420 mL** | |
|---|---|---|
| | 1M Tris pH 7 | 44.1 mL |
| 150 mM Tris pH 7.5 | 1M Tris pH 8 | 18.9 mL |
| 1.5M LiCl | 8M LiCl | 78.4 mL |
| 15 mM EDTA | 0.5M EDTA | 12.6 mL |
| 0.3% NP-40 | 10% NP-40 | 12.6 mL |
| | $H_2O$ | 253.4 mL |

| **Probe resuspension buffer (final):** | **100 mL** | |
|---|---|---|
| 5 mM Tris pH 7.5 | 1M Tris pH 7.5 | 500 uL |
| 0.05 mM EDTA | 0.5M EDTA | 10 uL |
| 0.01% Tween-20 | 10% Tween-20 | 100 uL |
| | $H_2O$ | 99.4 mL |

**Key Notes:**

1. To perform this procedure, 80 uL chimeric eCLIP samples should be saved immediately following the proteinase K step (when they are in 65 uL ProtK SDS buffer + 15 uL ProtK) - **BEFORE** the RNA cleanup & concentrator column
2. To increase the fraction of chimeric fragments (which need to be ≥40nt), the ethanol ratio in Silane cleanup steps following reverse transcription and cDNA adapter ligation can be decreased:
   - **Silane cleanup cDNA (after RT, ExoSAP, NaOH/HCl)**
     - Add 90 µl beads+RLTW to each sample
     - Add **60 µl** 100% EtOH (was 108 uL)
   - **Silane cleanup linker-ligated cDNA**
     - Add 45 µl beads+RLTW buffer mix to each sample
     - Add **30 µl** 100% EtOH (was 45 uL)

**Probe design:**

1. Take miRNA sequence — e.g. miR-145 — guccaguuuucccaggaaucccu
2. Reverse complement — miR-145rc — agggattcctgggaaaactggac
3. Add biotin and flanking genomic sequences — miR-145probe
   /5Biosg/CTAagggattcctgggaaaactggacCGT
4. For small RNA captures, sequences can be duplicated to increase recovery:

miR-145probe2
/5Biosg/CTAagggattcctgggaaaactggacCGTCTAagggattcctgggaaaactggacCTA

5. Order from IDT as standard ssDNA probes (standard desalting)

**Prepare capture probe oligonucleotide**

1. Resuspend oligo (IDT) to 100uM in probe resuspension buffer

**Couple capture probes to beads**

1. To a clean 1.5mL tube, add 75uL of beads (Dynabeads MyOne Streptavidin C1 Beads) per capture sample (prepare separate tubes for each probe set being used)
2. Wash beads 1.5 times with 2 volumes of 3x hybridization buffer (3xHyB) and aspirate supernatant
3. Resuspend beads in 100uL of 3xHyB per sample
4. Add 4uL per capture sample of 100 uM probes to resuspended beads
5. Vortex for 3 seconds
6. Incubate beads in the thermomixer for 10 min at 65°C
7. Vortex for 1-2 seconds and place on magnet
8. Add 600uL of 3xHyB and pipet mix
9. Spin down in mini-fuge (5 sec at ~3,000-5,000 rpm)
10. Place on magnet and aspirate supernatant
11. Resuspend in 3xHyB, 38uL per sample
12. Add 2.4uL 500 mM TCEP per sample and flick to mix
13. Spin down in mini-fuge

**Perform probe capture on sample**

1. Thaw chimeric eCLIP RNA samples (these should be after the 'Direct Proteinase K' step)
2. Separately incubate 80 uL Direct ProtK samples and coupled beads in a thermomixer set to:
   a. 1250rpm
   b. Begin at 72°C and ramp down to 67°C (should take about 2 mins)
3. Keeping tubes in thermomixer at 67°C, one at a time quickly add 40uL of the appropriate coupled beads to each sample
4. To hybridize samples, incubate in thermomixer (using the thermomixer lid):

   60°C - 30 mins

   50°C - 45 mins

   45°C - 15 mins

5. Spin down in mini-fuge
6. Place on magnet and remove supernatant (save supernatant if desired to confirm successful enrichment)
7. Prepare 1x HyB buffer (dilute 3x Hyb with $H_2O$)
8. Add 500uL of 1x HyB to beads, put back at 45°C for 3-5 minutes
9. Spin down in mini-fuge
10. Place on magnet and aspirate supernatant
11. Add 500uL of 1x HyB, place back in thermomixer while preparing DNase reaction mix

**Degrade DNA probes**

1. Prepare Turbo DNase reaction mix:

| Component | Vol. (1 rxn) |
|---|---|
| $H_2O$ | 64 uL |
| Turbo DNase buffer | 8 uL |
| Murine RNase inhibitor | 2 uL |
| Turbo DNase | 6 uL |

2. Put samples on magnet
3. Flip magnetic holder with tube (to wash lid), settle beads and aspirate supernatant
4. Spin down in mini-fuge, place on magnet and remove residual liquid with fine pipette
5. Remove samples from magnet and add 80uL of DNase master mix
6. Flick to mix and spin down in mini-fuge
7. Incubate samples in thermomixer, 37°C for 15 minutes, 1250 rpm
   NOTE: your samples are now eluted
8. Spin down in mini-fuge, place on magnet
9. Save **eluted sample**: transfer supernatant to a new labeled tube
10. Repeat elution by adding 40uL of pH8 TE (10 mM Tris pH 8, 1 mM EDTA)
11. Flick to mix and spin down in mini-fuge
12. Incubate beads in thermomixer at 65°C for 2 minutes
13. Spin down in mini-fuge, place on magnet
14. **Combine supernatant (re-elution sample)** with the first elution (step 9)
15. Store samples at -80°C until proceeding to standard chimeric eCLIP reverse transcription.

### Appendix B

#### Gene probe capture for chimeric-eCLIP

**Required reagents** (beyond standard chimeric eCLIP):

| | |
|---|---|
| 0.5M TCEP | Sigma 646547 |
| 8M LiCl | Sigma L7026 |
| Dynabeads MyOne Streptavidin C1 Beads | Invitrogen 65001 |

**Buffers**

| **3x Hybridization buffer:** | | **420 mL** |
|---|---|---|
| | 1M Tris pH 7 | 44.1 mL |
| 150 mM Tris pH 7.5 | 1M Tris pH 8 | 18.9 mL |
| 1.5M LiCl | 8M LiCl | 78.4 mL |
| 15 mM EDTA | 0.5M EDTA | 12.6 mL |
| 0.3% NP-40 | 10% NP-40 | 12.6 mL |
| | $H_2O$ | 253.4 mL |

| **Bead elution buffer:** | | **100 mL** |
|---|---|---|
| 5 mM Tris pH 7.5 | 1M Tris pH 7.5 | 500 uL |
| 0.05 mM EDTA | 0.5M EDTA | 10 uL |
| 0.01% Tween-20 | 10% Tween-20 | 100 uL |
| | $H_2O$ | 99.4 mL |

**Key Notes:**

To perform this procedure, 40 uL chimeric eCLIP samples should be saved immediately following completion of the **Silane cleanup linker-ligated cDNA** step (after magnetic separation and transfer of sample to a new tube)

**Probe design:**

1. Take the full mRNA sequence and design primers for amplification of complete mRNA sequence (including 5'- and 3'-UTRs, excluding introns)
2. Add the T7 promoter on the 5'-end primer. When RNA probes are synthesized, these RNA probes should be the sense strand of RNA.
3. Generate polyA cDNA from the cell type or tissue of interest using standard oligo-dT priming
    a. Note: Synthetic Gene Fragments can be used instead if convenient
4. PCR amplify the full mRNA sequence from cDNA using primers from (2).
5. Generate biotinylated RNA probes using an in-vitro transcription kit (e.g. NEB E2040S) with 90% of CTP and 10% biotin-CTP as well as 90% of UTP and 10% biotin-UTP.
6. After completion of in-vitro transcription, digest DNA template using standard Turbo DNAse reaction
7. Purify RNA using NEB's Monarch® RNA Cleanup Kit (500 µg) (T2050S)
8. Measure RNA probe concentration using NanoDrop or TapeStation.

**Prepare probe-coupled capture beads**

1. Dilute probes (10 ug per sample, up to 40 ug per reaction) in 475 uL $H_2O$ in a 1.5 mL tube
2. Pre-warm Thermomixer to 95ºC
3. Add 25 uL 0.5M TCEP to diluted probes
4. Incubate diluted probes in Thermomixer at 95ºC for 2 minutes and place on ice until **step 14**
5. Adjust Thermomixer to 60ºC
6. Vortex MyOne Streptavidin C1 Dynabeads to resuspend
7. Per sample (maximum 4 samples per tube):
    a. Aliquot 100 uL MyOne Streptavidin C1 Dynabeads to a new 1.5 mL tube
    b. Add 100 uL 3x Hybridization buffer
8. Place tube on magnet and allow beads to separate until solution is clear
9. Aspirate supernatant and discard
10. Resuspend beads in 1 mL 3x Hybridization buffer
11. Place tube on magnet and allow beads to separate until solution is clear
12. Aspirate supernatant and discard
13. Resuspend beads in 100 uL 3x Hybridization buffer
14. Add beads to diluted probes (step 4) and mix by inverting the tube 5 times
15. Incubate in Thermomixer at 60ºC for 10 mins (with interval shaking at 1250 rpm, 15s on / 15s off)
16. Place tube on magnet and allow beads to separate until solution is clear
17. Adjust Thermomixer to 90ºC
18. Aspirate supernatant and discard
19. Resuspend beads in 1 mL 3x Hybridization buffer
20. Place tube on magnet and allow beads to separate until solution is clear
21. Aspirate supernatant and discard
22. Spin tube in minifuge, place on magnet, and remove all residual supernatant
23. Resuspend probe-coupled beads in 20 uL 3x Hybridization buffer
24. Keep probe-coupled beads at room temperature until use.

**Perform probe capture on sample**

1. Pre-heat Thermomixer to 90ºC
2. Perform capture:

    Notes: Keep tubes in Thermomixer block at 1200 rpm during entire protocol

    Do not place samples on ice or allow samples to cool and renature

    a. Take cDNA samples from **Silane cleanup linker-ligated cDNA** step and place in pre-heated 90ºC Thermomixer block for 1 min
    b. Decrease temperature to 72ºC
    c. When temperature reaches 72ºC, place probe-coupled beads (step 24 above) on the thermomixer for 1 minute, then add 20 uL of probe-coupled beads to each sample
    d. Incubate 72ºC for 1 additional minute
    e. Incubate 65ºC for 10 minutes
    f. Incubate 60ºC for 20 minutes
    g. Incubate 50ºC for 60 minutes
3. Prepare 1.2 mL 1x Hybridization buffer per sample
    a. Dilute 3x Hybridization in $H_2O$
    b. Pre-warm 1x Hybridization buffer at 50ºC
4. Wash beads:
    a. Take sample from 50ºC and place on magnetic rack
    b. As soon as beads separate, aspirate supernatant and discard
    c. Add 500 uL of pre-warmed 1x Hybridization buffer and invert to mix

   d. Place sample back on Thermomixer at 50ºC and incubate for 5 mins
5. Repeat step 4 (wash) 1 time for a total of 2 washes
6. Place tube on magnet and allow beads to separate until solution is clear
7. Aspirate supernatant and discard
8. Resuspend beads in 55 uL Bead Elution Buffer
9. Elute captured cDNA:
   a. Add 6 uL RNase enzyme mix
   b. Vortex samples for 2 sec and spin down in minifuge
   c. Incubate 37ºC for 10 mins at 1200 rpm
   d. Adjust Thermomixer to 70ºC
   e. Add 4 uL of 2M NaOH to each sample
   f. Vortex samples for 2 sec and spin down in minifuge
   g. Incubate 70ºC for 10 mins at 1200 rpm
   h. Spin tube in minifuge and place on magnet
   i. Transfer supernatant (**containing captured cDNA**) to a new 1.5 mL tube
10. Repeat elution:
   a. To beads, add 30 uL 100 mM NaOH and pipette mix to resuspend
   b. Incubate 70ºC for 2 mins at 1200 rpm
   c. Spin tube in minifuge and place on magnet
   d. Combine supernatant (**containing captured cDNA**) with the previous supernatant (step 9i above)
11. Add 3.67 uL 3M HCl to neutralize pH
12. Flick tube to mix
13. Add 21 uL $H_2O$ to bring final volume to 120 uL

**Silane cleanup captured cDNA**

- **Prepare beads:**
  - To 10 µl **MyONE Silane Beads** per sample, add 5x volume RLT (e.g. for 4 samples, use 40 µl of beads and add 200 µl of RLT)
  - Pipette mix, magnetically separate, and remove supernatant
  - Resuspend beads in 365 µl RLTW buffer (RLT + 0.025% Tween-20) **per sample**. Mix well.

- **Bind cDNA:**
  - Add 360 µl of bead/RLTW mixture above to each 120 uL cDNA sample (**step 13** above) and mix
  - Add 480 µl **100% EtOH** to each sample
  - Close tube well and place on rotator for 10 minutes

- **Wash beads:**
  - Magnetically separate and discard supernatant
  - Add 1 mL (1.5 mL tubes) **80% EtOH**, pipette resuspend
  - After 30 s, magnetically separate, remove supernatant
  - Repeat wash with 1 mL (1.5 mL tubes) **80% EtOH**
  - After 30 s, magnetically separate, remove supernatant
  - Wash 3$^{rd}$ time with 750 µl (1.5 mL tubes) **80% EtOH.**
  - Spin briefly in picoFuge, magnetically separate, remove residual liquid with fine tip
  - Dry beads well, i.e. until they stop "shining - no ethanol should be left on the bottom of strip. Beads are over-dry when they change color from brown to orange/rusty color.

- **Elute cDNA:**
  - Resuspend in **21 µl Bead Elution Buffer**, let sit for 5 min

- Magnetically separate, transfer supernatants to strip tube(s)
- Proceed to qPCR in standard chimeric eCLIP protocol
